# Postmortem brain MRI reveals differential associations of subcortical and limbic volumes with cortical thinning and neurodegenerative pathologies

**DOI:** 10.64898/2025.12.30.697140

**Authors:** Pulkit Khandelwal, Michael Tran Duong, Lisa M. Levorse, Winifred Trotman, Alejandra Bahena, Sydney A. Lim, Amanda E. Denning, Eunice Chung, Christopher A. Olm, Hamsanandini Radhakrishnan, Ranjit Ittyerah, Karthik Prabhakaran, Gabor Mizsei, Theresa Schuck, Sheina Emrani, Joaquin A. Vizcarra, John Robinson, Daniel T. Ohm, Jeffrey S. Phillips, Jesse Cohen, Laura E.M. Wisse, John A. Detre, Ilya M. Nasrallah, Christopher A. Brown, Sandhitsu R. Das, Edward B. Lee, M. Dylan Tisdall, David J. Irwin, Corey T. McMillan, David A. Wolk, Paul A. Yushkevich

**Author notes:** Correspondence: Pulkit Khandelwal 149 13th Street Charlestown, MA 02129, United States and Michael Tran Duong 3400 Spruce Street Philadelphia, PA 19104, United States and. Authors contributed equally (Pulkit Khandelwal, Michael Tran Duong).

## Abstract

**Introduction:** The impact of different neuropathologies on deep brain structures remains to be understood. We examine subcortical and limbic volumetry in neurodegenerative diseases involving p-tau, α-synuclein and TDP-43.

**Methods:** We acquired neuropathological measures and brain segmentations from postmortem analysis of 132 donors with Alzheimer’s disease (AD), Lewy body disease (LBD), Frontotemporal Lobar Degeneration with TDP-43 (FTLD-TDP) and FTLD-Tau.

**Results:** LBD had the least subcortical, limbic and cortical atrophy compared to AD, FTLD-TDP and FTLD-Tau. In donors with both AD and LBD pathologies, primary LBD was associated with less atrophy than primary AD. While AD had cortico-subcortical and cortico-limbic morphometric associations, LBD had more limited parieto-occipital cortico-limbic associations. FTLD-TDP had cortico-subcortical while FTLD-Tau had cortico-subcortical and cortico-limbic associations. In AD and FTLD-Tau, hippocampal volumes correlated with p-tau burden, neuron loss and gliosis. In LBD, thalamic α-synuclein severity was associated with subcortical/limbic volumes.

**Discussion:** Postmortem neuroimaging reveals disease- and region-specific structure-pathology relationships.

## 1. Background

Deep within the brain, subcortical and limbic structures harbor essential neural circuits with critical functions and susceptibilities to different neuropathologies such as neurofibrillary phosphorylated tau (p-tau), α-synuclein and Transactive Response DNA-binding Protein 43 kDa (TDP-43). Applying postmortem imaging and pathological evaluation can further parse distinct profiles of subcortical, limbic and cortical atrophy in structure-structure correspondences and structure-pathology relationships in different neurodegenerative diseases. Thus far, studies converge on structural changes in several key limbic and subcortical structures such as hippocampus, amygdala, thalamus and striatum in the majority of neurodegenerative diseases [1, 2, 3, 4]. These include Alzheimer’s disease (AD) [5, 6, 7, 8, 9, 10, 11, 12, 13, 14, 15], Lewy Body disease (LBD) [14, 16, 17, 18], Frontotemporal Lobar Degeneration with TDP-43 pathology (FTLD-TDP) [19, 20, 21, 22, 23] and FTLD-Tau [10, 22, 24, 25, 26].

Nonetheless, substantial biological heterogeneity and polypathology exists between patients [27, 28] with the majority of patients harboring more than one neuropathological diagnosis at autopsy [3, 29, 30]. Therefore, to characterize relationships between regional subcortical/limbic volume loss, cortical atrophy and different neuropathologies, we need accurate measures of brain morphology and pathological burden across patients and diagnostic categories. Several prior studies already investigated structure-pathology relationships by comparing antemortem imaging with postmortem pathology, but the variable time gap between these assessments may confound observed associations. In this study, we instead examine cortical, subcortical and limbic associations between postmortem brain magnetic resonance imaging (MRI), histopathology and immunohistochemistry. Postmortem imaging offers remarkable advantages over antemortem MRI for linking regional neuroanatomy and morphometry, enabling both higher spatial resolution and correlations between brain structure and pathological drivers at the same point in time [6, 8, 12, 31, 32, 33, 34, 35, 36].

Here, we introduce a unique dataset of 132 postmortem specimens with primary neuropathological diagnoses of AD, LBD, FTLD-TDP and FTLD-Tau, along with histopathological markers and high-resolution postmortem MRI. With a deep learning model SuperSynth [37, 38], we obtained segmentations for five subcortical structures (caudate, putamen, nucleus accumbens, thalamus, pallidum) and two limbic regions (hippocampus, amygdala). Cortical segmentations were generated with purple-mri [35, 36].

With this data-rich cohort, we sought to juxtapose subcortical and limbic volumes with cortical thickness and pathology markers to clarify and expand on prior observations of differential volume loss patterns, structure-structure correspondences and structure-pathology relationships across diseases. For instance, we predicted that donors with FTLD-TDP and FTLD-Tau have more subcortical and limbic volume loss than AD, given that non-AD p-tau and TDP-43 pathologies start early in subcortical and limbic regions [10, 22]. We hypothesized that donors with primary LBD would have less subcortical and limbic atrophy than other groups, especially AD, given that neuronal α-synuclein inclusions are linked to changes in neurotransmission and neuronal metabolism and less associated with volume loss [16, 39, 40, 41]. We aimed to uncover cortical thinning patterns associated with total subcortical and limbic volume loss across diseases. Importantly, we posited that regional burden of primary neuropathology would correlate with subcortical and limbic volume loss, as it does with cortical atrophy [3, 17, 19, 33]. Finally, we proposed that regional histologic neuronal loss and gliosis are associated with gross volume loss. Overall, by integrating postmortem imaging and pathology, our approach enables us to substantiate these predictions across neurodegenerative diagnoses.

## 2. Methods

### 2.1. Patient donor cohort

We analyze a dataset of 132 donors from the Penn Integrated Neurodegenerative Disease Database [42]. Patients were evaluated at the Penn Frontotemporal Degeneration Center or Alzheimer’s Disease Research Center and followed to autopsy at the Penn Center for Neurodegenerative Disease Research (CNDR) [25, 43, 44]. Human brain specimens were obtained with informed consent from next of kin at time of death. Patients were evaluated per standard antemortem diagnostic criteria and postmortem autopsy and imaging [42]. **Table 1** details primary and secondary neuropathological diagnoses. Groups included AD, LBD, FTLD-TDP and FTLD-Tau. Here, we denote FTLD-Tau to include 3R tauopathies such as Pick’s disease and 4R tauopathies such as Progressive Supranuclear Palsy (PSP) and Corticobasal Degeneration (CBD). Disease-associated genetic variant data was acquired. Study procedures were performed with written informed consent per Institutional Review Board guidelines and the Declaration of Helsinki.

**Table 1.**
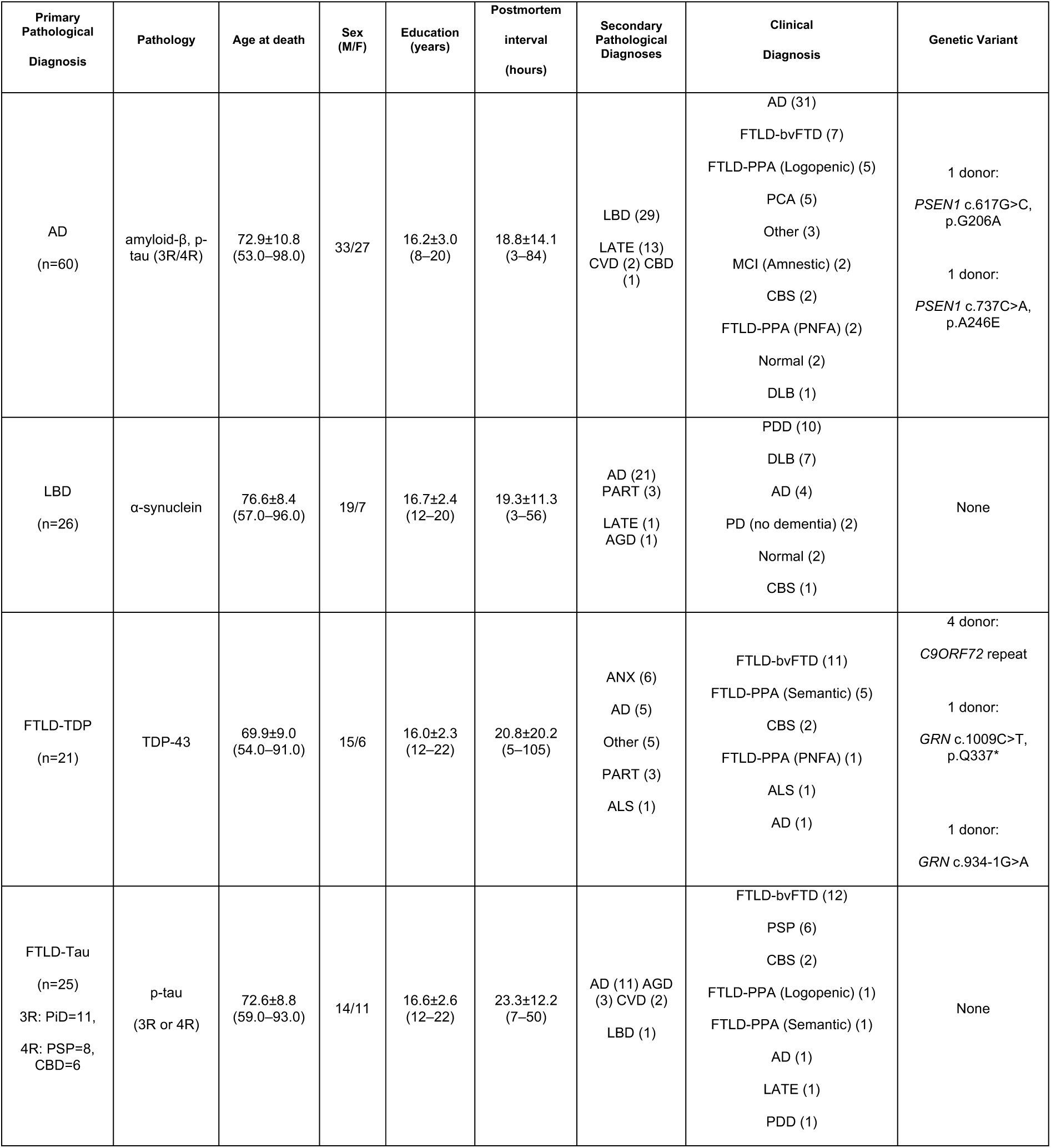
Clinical characteristics of the donor cohort. The table summarizes group-level characteristics for donors with neuropathological diagnoses. Abbreviations: AD Alzheimer’s disease. AGD Argyrophilic Grain Disease. ALS Amyotrophic Lateral Sclerosis. ANX Annexinopathy. bvFTD behavioral variant Frontotemporal Dementia. CAA Cerebral Amyloid Angiopathy. CBD Corticobasal Degeneration. CBS Corticobasal Syndrome. CVD Cerebrovascular Disease. *C9ORF72* Chromosome 9 Open Reading Frame 72 gene. DLB Dementia with Lewy bodies. F female. FTLD-TDP Frontotemporal Lobar Degeneration with Transactive response DNA binding Protein-43 kiloDaltons (TDP-43) pathology. *GRN* progranulin gene. LATE Limbic-Predominant Age-Related Transactive response DNA binding Protein-43 kiloDaltons (TDP-43) Encephalopathy. LBD Lewy Body Disease. M male. MCI Mild Cognitive Impairment. PART Primary Age-Related Tauopathy. PCA Posterior Cortical Atrophy. PD Parkinson’s disease. PDD Parkinson’s disease dementia. PiD Pick’s disease. PNFA Progressive Nonfluent Aphasia. PSP Progressive Supranuclear Palsy. PPA Primary Progressive Aphasia. *PSEN1* presenilin 1 gene. 3R 3-repeat tau. 3R/4R mixed 3-repeat/4-repeat tau. 4R 4-repeat tau.

### 2.2. Image acquisition

#### 2.2.1. Antemortem imaging procedure

Antemortem T1-weighted brain MRI scans enabled intracranial volume (ICV) calculation to normalize regional volumes. Imaging parameters varied because scans were acquired between 2006 and 2021, across a range of scanners with varying field strengths 1.5T (n=11), 3T (n=118), or 7T (n=3) (Magnetom Sonata, Verio, Trio, TIM Trio, Prisma, Prisma Fit, and Terra; Siemens Healthineers, Erlangen, Germany). Median (range) parameters included: repetition time 2300 ms (5-3000), echo time 2.98 ms (0.002-7.72), echo numbers 1.0 (0.0-4.0), flip angle 9.0 [10, 14, 22, 23, 24, 34, 37, 45, 46, 47, 48, 49, 50, 51, 52, 53, 54, 55, 56], and number of averages 1.0 (0-4.0) [3].

#### 2.2.2. Postmortem imaging procedure

After removing cerebellum and brainstem, one hemisphere was immersed in 10% neutral buffered formalin for >4 weeks before imaging, with formalin replaced weekly to ensure complete fixation. Meninges and vasculature were removed 2-4 days before scanning. The hemisphere was removed from fixative one day before scanning and desiccated for 3 hours. After fixation, samples were placed in Fomblin (California Vacuum Technology; Freemont, CA) and enclosed in custom-built 3D printed holders within a sealed bag. Samples were left to rest to allow air bubbles to escape from tissue. Samples were scanned using a custom-built small solenoid coil or a custom-modified quadrature birdcage coil (Varian, Palo Alto, CA, USA) [43, 57]. Then, samples were placed into a whole-body 7T Siemens Terra scanner with foam shims under the coils to raise them off the table and near the isocenter. T2w images were acquired using a 3D-encoded T2 SPACE sequence with 0.3×0.3×0.3 mm³ isotropic resolution, repetition time 3 s, echo time 383 ms, turbo factor 188, echo train duration 951 ms, bandwidth 348 Hertz/pixel in 2-3 hours per measurement. Image reconstruction applied vendor reconstruction software, combining signal averages in k-space, and producing magnitude images for each echo. Four repeat measurements were acquired per sample and averaged to generate the final image.

### 2.3. Image analysis

Volumes of nucleus accumbens, caudate, putamen, thalamus, globus pallidus (pallidum), hippocampus, and amygdala were computed via automated segmentation of postmortem MRI by SuperSynth. For cortical thickness measurements, parcellations based on the Desikan-Killiany-Tourville (DKT) [55] atlas were obtained for each postmortem MRI with purple-mri. SuperSynth has more subcortical and limbic regions included while purple-mri has a complete cortical parcellation. All segmentations underwent quality control.

### 2.4. Semi-quantitative neuropathology and histology measures

Brain hemispheres were split for imaging and immunohistochemistry. Tissue blocks of approximately 1.5×1.5×0.5cm^3^ were extracted from the contralateral hemisphere and embedded in paraffin which were then sectioned into 6 μm slices for immunohistochemistry using monoclonal antibodies to detect amyloid-β (NAB228, 1:8000, CNDR), p-tau (PHF-1, 1:1000, Dr. Peter Davies), TDP-43 (TAR5P-1D3, 1:500, Drs.

Manuela Neumann and E. Kremmer) and α-synuclein (Syn303, 1:16,000, CNDR). Hematoxylin and Eosin staining was performed to assess neuronal and glial density. Semi-quantitative scores of p-tau, TDP-43, α-synuclein, gliosis, neuronal loss, and global vascular measures were visually assigned by expert neuropathologists (Drs. Edward B. Lee and John Trojanowski) on a scale of 0–3, from 0 none, 0.5 rare, 1 mild, 2 moderate and 3 severe. These ratings are based on consensus criteria agreed upon by the authors and facilitated by visual exemplars. Neuronal loss ang gliosis scores were determined by visual semiquantitative assessment of different cell morphologies (neurons vs. glia) in each field of view relative to the total tissue imaged. Pathology burdens for subcortical and limbic regions were available for tissue samples derived from caudate, putamen, thalamus, pallidum, amygdala and hippocampus (computed from a mean between CA1/subiculum, dentate gyrus and entorhinal cortex). Pathology ratings were not available for nucleus accumbens. Pathology ratings were recorded for additional regions including precentral gyrus, postcentral gyrus, middle frontal gyrus, angular gyrus, superior middle temporal gyrus, cingulate gyrus, occipital cortex, entorhinal cortex, midbrain, substantia nigra, pons, locus coeruleus, brainstem and cerebellum. The ratings across all these regions allowed neuropathologists to comprehensively evaluate the magnitude and distribution of different neuropathologies and adjudicate between primary and secondary diagnoses. **Figure 1** shows the distribution of the neuropathology ratings across different subcortical and limbic regions along with global markers.

**Figure 1.**
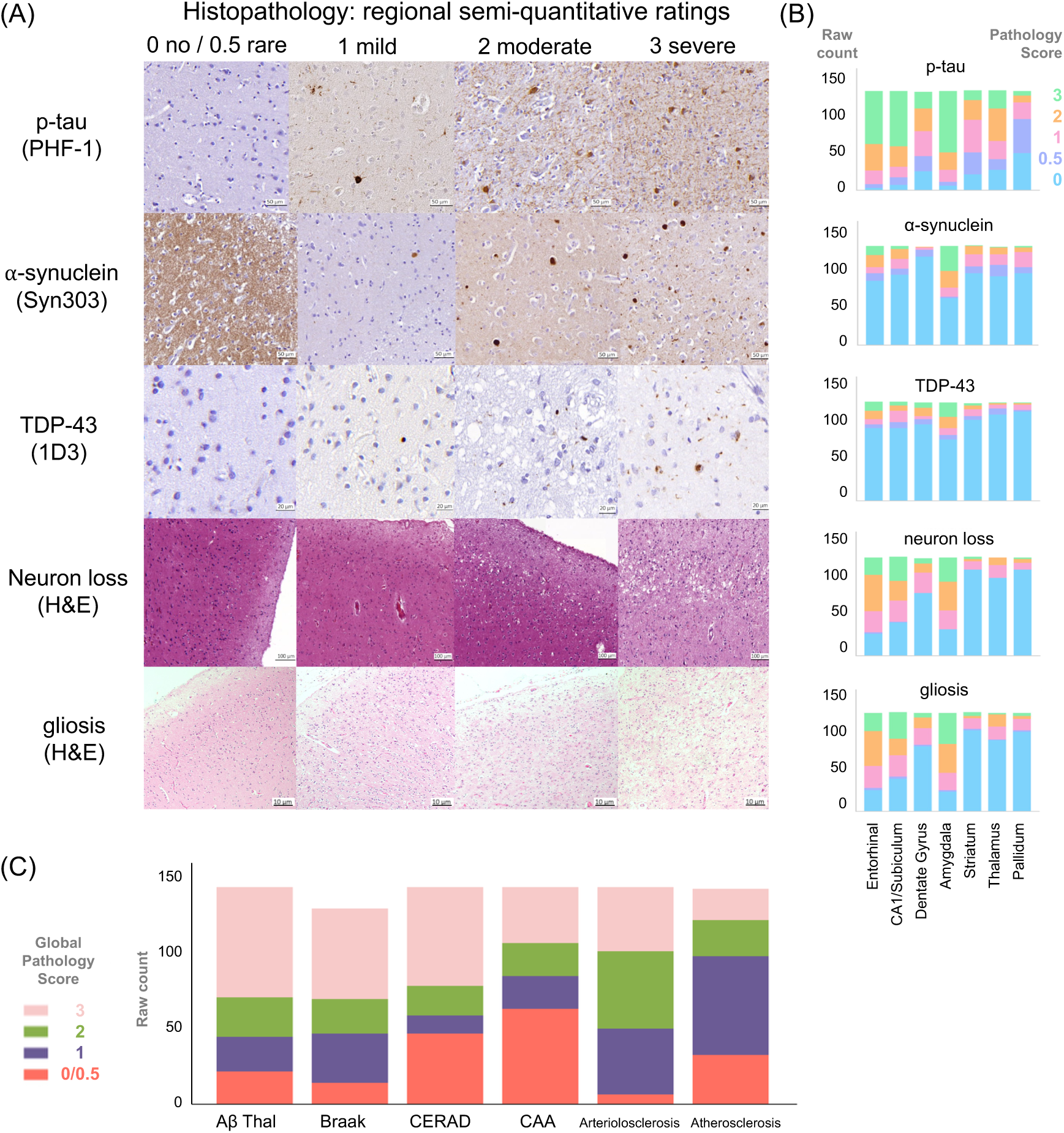
Histopathology ratings across the autopsy cohort. (**A**) Example immunohistochemistry images. Distributions via stacked bar plots of raw counts for pathology markers, including (**B**) proteinopathy and histological markers (p-tau, TDP-43, α-synuclein, neuronal loss and gliosis) across entorhinal cortex, CA1/subiculum, dentate gyrus, amygdala, striatum (caudate/putamen), thalamus and pallidum, as well as (**C**) additional global histopathology markers (Aβ Thal phase, Braak neurofibrillary tangle stage, CERAD neuritic plaque stage, cerebral amyloid angiopathy (CAA), arteriolosclerosis, and atherosclerosis measures).

### 2.5. Statistical analysis

To evaluate volumetric and pathological relationships across disease groups, we performed a series of group-level analyses (**Figure 2**). For all analyses, regional volumetric measures from postmortem MRI were normalized by ICV to reduce inter-subject variability. All models included covariates of age at death, sex, education, and postmortem interval to fixation. Antemortem comparison models included antemortem imaging interval rather than postmortem interval. All statistical tests were two-tailed, with significance set at p<0.05 and corrected for multiple comparisons using Benjamini–Hochberg false-discovery-rate (FDR) procedure. All statistical analyses were conducted using statsmodel [58] in Python. We subdivide the FTLD-tau group into 3R tauopathy (Pick’s disease) and 4R tauopathy (PSP and CBD) for further comparative analysis.

**Figure 2.**
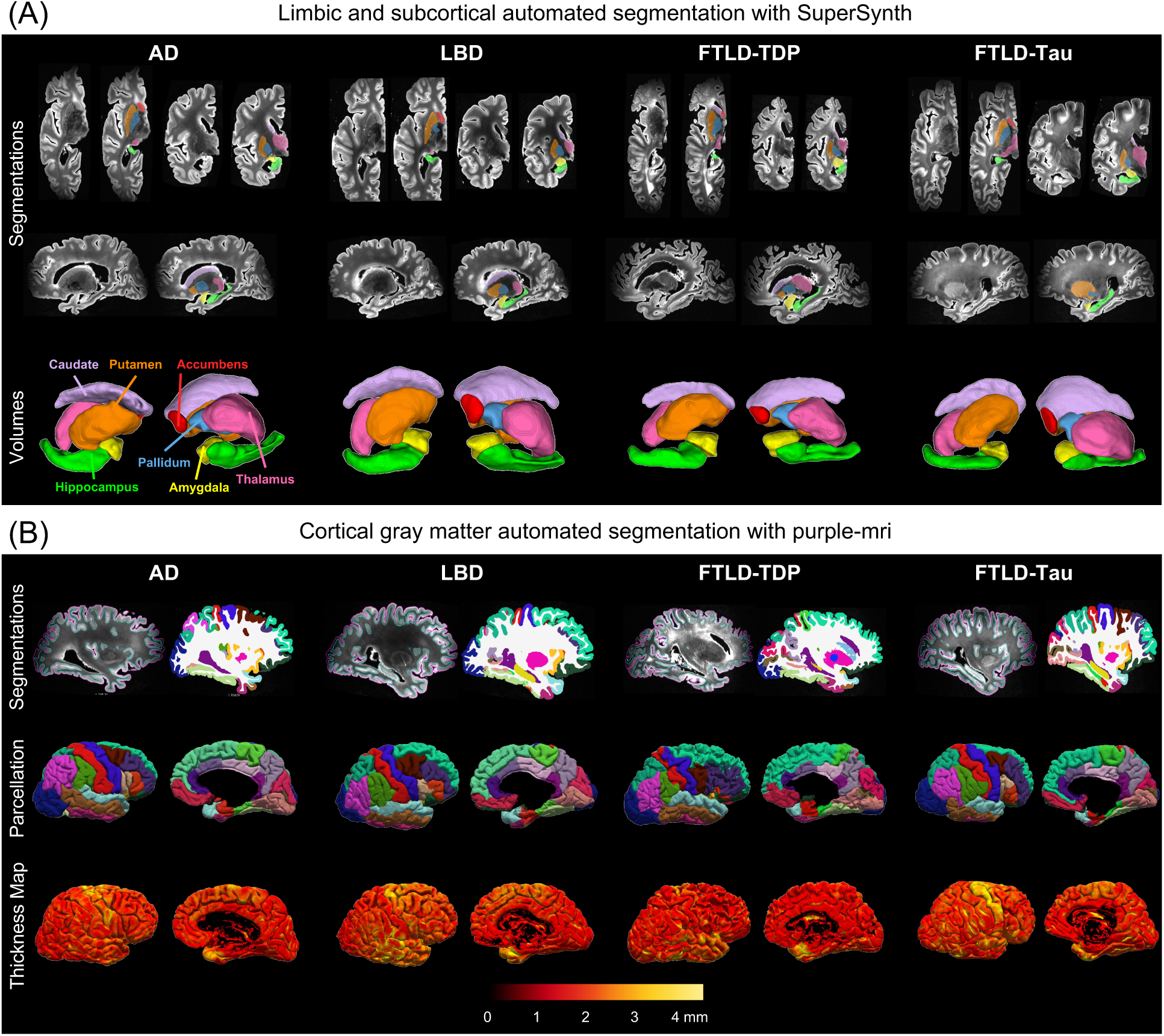
Deep learning 3D subcortical and limbic structure segmentations and whole-hemisphere surface-based parcellation. (**A**) The automated segmentations for hippocampus, amygdala, nucleus accumbens, putamen, caudate, thalamus and globus pallidus were obtained via SuperSynth [37, 38]. Shown are MRIs with segmentation overlaid for axial, coronal, and sagittal planes along with 3D renderings for the lateral and medial views for four specimens on T2w postmortem MRI. (**B**) Cortical gray matter was parcellated based on the Desikan-Killiany-Tourville (DKT) atlas with purple-mri [35, 36] followed by cortical thickness estimation in native subject-space resolution. Shown are gray matter (pink) and white matter (cyan) surfaces on the left along with the cortical parcellation on the right and computed cortical thickness [72].

#### 2.5.1. Structural group differences and subcortical-cortical associations

To assess the differential volumetric patterns of hippocampus, amygdala, nucleus accumbens, thalamus, caudate, putamen and pallidum, we computed ANOVA group comparisons and then pairwise within-group differences with likelihood-ratio tests via nested linear models across groups. To examine regional relationships between cortical and subcortical matter, we computed partial Spearman correlations between mean thickness of each DKT atlas-based cortical region and the total weighted composite subcortical and limbic volume.

#### 2.5.2. Structure-pathology associations

To determine relationships between regional semi-quantitative neuropathology ratings and subcortical/limbic volumes within each diagnostic group, we fit an independent ordinary least squares (OLS) regression model with volume as the dependent variable and the region’s predominant primary disease-specific pathology measure as the independent variable accounting for covariates as above, with slope coefficients β and residual error ε. For primary pathology models, AD and FTLD-Tau were compared with p-tau, FTLD-TDP with TDP-43, and LBD with α-synuclein regional burden. Next, we fitted several polypathology OLS models to examine the effects of volume loss on secondary concomitant molecular pathologies [3].

Primary pathology model:

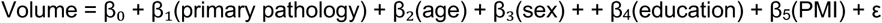

Polypathology model:

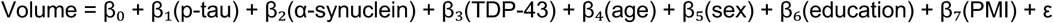

#### 2.5.3. Structure-histology feature correlations

To test if MRI-derived volumetry reflects histopathological markers of neurodegeneration independent of pathology type, we fit an OLS model with volume as the dependent variable and regional neuronal loss and gliosis measures as the independent variable accounting for covariates as above. Furthermore, we performed mediation analyses to determine if neuronal loss and gliosis mediated the interaction between primary disease-specific regional pathology and its effects on structural volumetry. Finally, we assessed relationships between global vascular disease burden and structural volume for each structure by computing OLS regression adjusting for covariates between volume and global pathology scores in CAA, arteriolosclerosis, and atherosclerosis.

## 3. Results

### 3.1. Pipeline for postmortem imaging, segmentation and histopathology

Our dataset comprised 132 postmortem donor specimens spanning four primary neuropathological diagnoses: AD (n=60), LBD (n=26), FTLD-Tau (n=25), and FTLD-TDP (n=21) (**Table 1**). While most donors had sporadic disease, 9 donors had disease-associated genetic variants. We found 2 donors with familial AD associated with *PSEN1* exon mutations, 4 donors with FTLD-TDP due to *C9ORF72* hexanucleotide repeat expansion and 2 donors with FTLD-TDP due to *GRN* non-coding mutations. Across the cohort, mean age at death was 73±10 years with an average of 16±3 years of education. The postmortem interval (PMI) averaged 20±14 hours. MRI segmentations were performed for subcortical and limbic regions via SuperSynth along with the cortical gray matter DKT atlas-based volumetric segmentations and surface-based parcellations via purple-mri (**Figure 2**).

### 3.2. AD, FTLD-TDP and FTLD-Tau have lower subcortical and limbic volumes than LBD

First, we compared relationships between postmortem subcortical/limbic volumes and diagnostic groups adjusting for covariates and multiple comparisons (**Figure 3A, Supplementary Table S1.1**). Donors with AD had significantly lower volumes than those with LBD in all subcortical and limbic regions studied (p<0.05) except caudate and globus pallidus. Donors with FTLD-Tau and FTLD-TDP had lower postmortem volumes than those with LBD across all subcortical and limbic regions queried (p<0.05). Donors with FTLD-TDP displayed the lowest volumes across the groups and had significantly worse volume loss than those with AD for caudate, putamen and nucleus accumbens (p<0.05). When we split the FTLD-Tau group into 3R tauopathy and 4R tauopathy, we found significant relationships of worse subcortical volume loss compared to AD in these same regions (**Supplementary Figure S2.1**). Postmortem findings paralleled antemortem imaging (**Supplementary Figure S1.1**). In comparisons that covaried for age, sex, education and antemortem imaging interval, we similarly found that patients with LBD had higher antemortem volumes across subcortical and limbic regions than AD, FTLD-Tau and FTLD-TDP.

**Figure 3.**
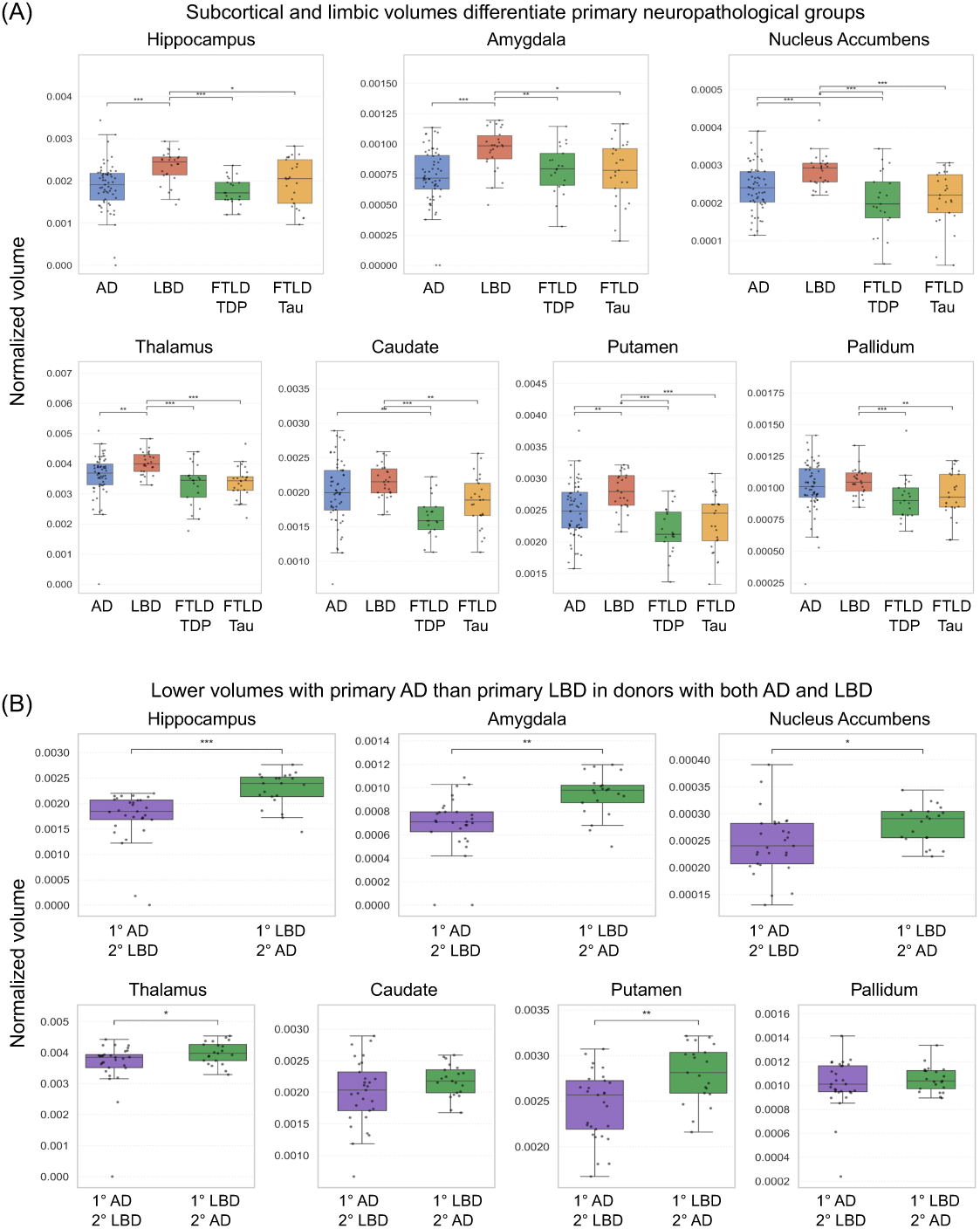
Postmortem limbic and subcortical volumes across neuropathological groups. (**A**) Boxplots show ICV-normalized postmortem single hemisphere volumes for the seven subcortical and limbic regions across four primary diagnostic categories: Alzheimer’s disease (AD), Lewy body disease (LBD), Frontotemporal Lobar Degeneration with TDP-43 pathology (FTLD-TDP), and FTLD-tau. (**B**) Boxplots compare ICV-normalized postmortem volumes across regions for primary AD and secondary LBD (n = 29) vs. primary LBD and secondary AD (n = 21) in donors with both AD and LBD pathology. Pairwise differences between groups were assessed using likelihood-ratio tests adjusting for age at death, sex, education, and postmortem interval. Statistically significant pairwise comparisons with multiple test adjustment are denoted with *p<0.05, **p<0.01, ***p<0.001. In each boxplot, the center line indicates median, box edges represent 25th–75th percentiles (IQR), whiskers extend to 1.5×IQR, and overlaid points correspond to individuals. Detailed statistical results are reported in **Supplementary Table S1.1**.

### 3.3. Primary AD is linked to worse subcortical and cortical atrophy than primary LBD in donors with both AD and LBD copathologies

Since AD and LBD pathologies often coexist, so we aimed to better delineate the relationships between volume loss and primary AD vs. primary LBD pathology (**Figure 3B**). Our autopsy cohort includes 29 donors with primary AD and secondary LBD and 21 donors with primary LBD and secondary AD. Similar to the omnibus findings, donors with primary AD and secondary LBD had significantly lower volumes than donors with primary LBD and secondary AD in all subcortical and limbic regions studied (p<0.05) except globus pallidus and caudate. We also assessed postmortem cortical thicknesses from purple-mri. Across almost all cortical regions except medial occipital cortex, sensorimotor cortex, superior parietal lobule and pars orbitalis, donors with primary AD and secondary LBD had lower postmortem cortical thickness than donors with primary LBD and secondary AD (**Supplementary Figure S1.2**).

Comparing cortical thickness across the four main disease groups, we found LBD also had less cortical atrophy than FTLD-Tau and FTLD-TDP in nearly all 35 cortical regions (**Supplementary Figure S1.3**). Intriguingly, we found that superior temporal gyrus and insula had significantly lower postmortem thickness in FTLD-TDP than FTLD-tau, suggesting these regions may help differentiate these clinical entities. Overall, these results strengthen the notion that AD more so than LBD pathology drives cortical, subcortical and limbic atrophy, even in patients with mixed AD and LBD.

### 3.4. Subcortical and limbic volume loss tracks with cortical thinning differently in each disease

Since volume changes occurred across several subcortical and limbic structures in a fairly similar manner for these diseases, we first assessed how cortical thinning correlated with a composite subcortical and limbic meta-region (comprised of hippocampus, amygdala, nucleus accumbens, thalamus, caudate, putamen and pallidum) (**Figure 4A**). Composite subcortical and limbic volume loss showed positive pathology-volume correlations with thinning of nearly all cortical regions in donors with AD (p<0.05). However, donors with FTLD-TDP and FTLD-Tau showed positive correlations limited to frontal regions (such as superior frontal gyrus and cingulate gyrus), and inferior parietal lobule (p<0.05). Donors with LBD had less atrophy such that associations between cortical thinning and composite subcortical/limbic volume loss were not found. Separately evaluating the 3R and 4R tauopathy subgroups individually also did not show significant correlations between subcortical volume and cortical thinning (**Supplementary Figure S2.2**).

**Figure 4.**
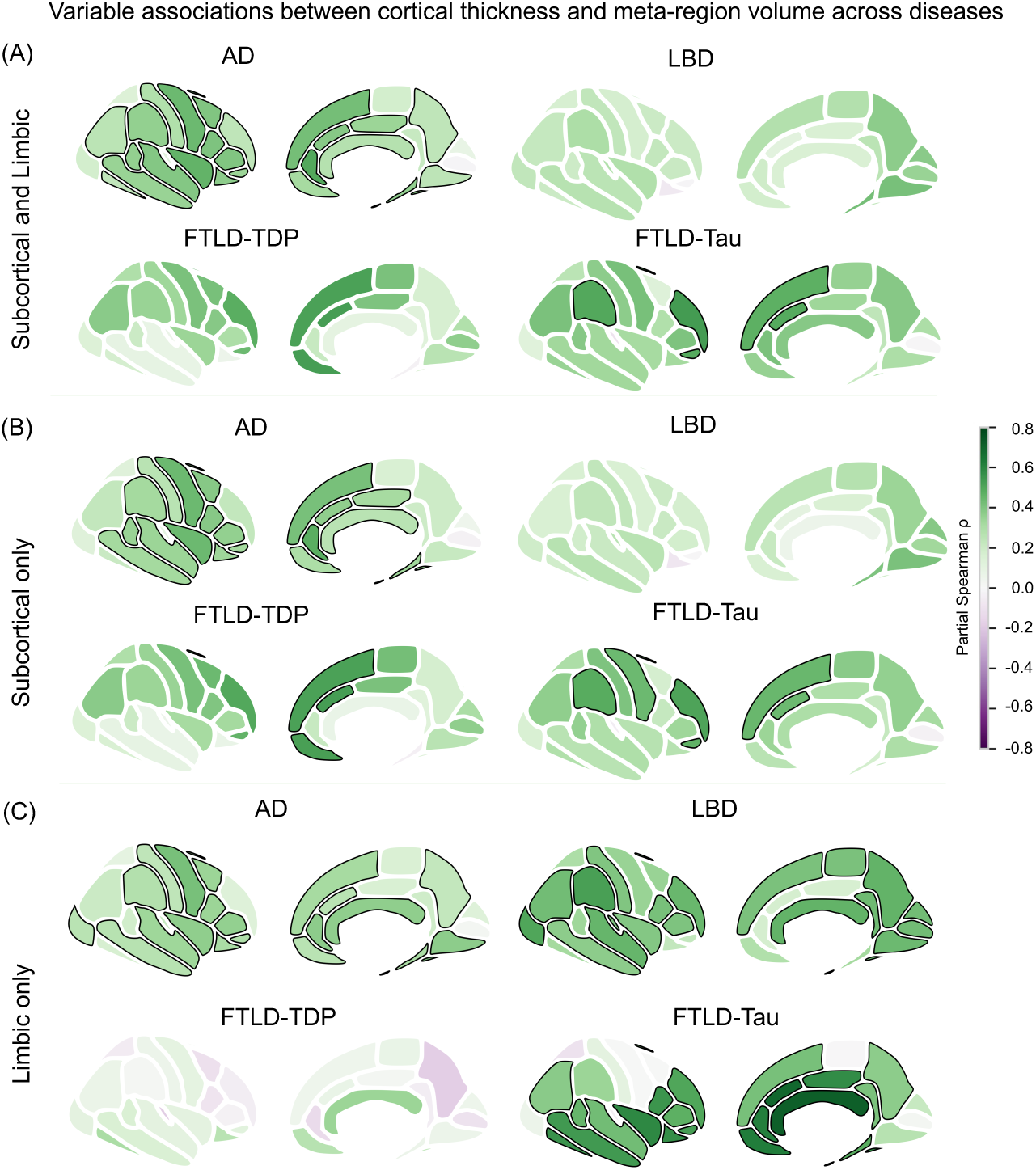
Associations between Desikan-Killiany-Tourville atlas-based regional mean thickness and weighted subcortical and limbic volumes. Disease-specific regional correlations between cortical thickness and (**A**) a meta-region of subcortical and limbic structures computed by a volume-weighted, ICV-normalized composite (including caudate, putamen, pallidum, thalamus, nucleus accumbens, hippocampus and amygdala), (**B**) subcortical structures only (caudate, putamen, pallidum, thalamus, nucleus accumbens) and (**C**) limbic structures only (hippocampus and amygdala). For each neuropathological group, partial Spearman correlations were calculated between mean ROI-level Desikan-Killiany-Tourville cortical thickness and normalized weighted volume, adjusting for age at death, sex, education, and postmortem interval. Partial Spearman coefficients (ρ) are reported with regions outlined in black indicating p<0.05.

To ascertain whether the correspondence between cortical thinning and total subcortical/limbic volume loss was largely driven by variation in the medial temporal lobe, we then compared regional cortical thickness with volume for a second subcortical-only meta-region that includes only subcortical regions (caudate, putamen, accumbens, thalamus, pallidum) (**Figure 4B**). This modified subcortical composite volume showed significant correlations with most frontal, parietal and temporal regions in AD (p<0.05) and frontal and lateral parietal regions in FTLD-TDP and FTLD-Tau (p<0.05). LBD did not show cortical-subcortical associations were not significant.

Closing the loop, we tested how cortical thinning patterns change with limbic volume loss across diseases with a limbic only region (hippocampus and amygdala) (**Figure 4C**). We recapitulated the finding of diffuse cortical-limbic associations in AD (p<0.05). Interestingly, compared to the subcortical volumes, donors with LBD had strong associations between frontal and parieto-occipital atrophy and limbic volume loss (p<0.05). Moreover, we observed robust significant cortico-limbic morphometric associations in FTLD-Tau (p<0.05) but not FTLD-TDP, juxtaposed to the significant cortico-subcortical associations in FTLD-TDP.

### 3.5. Primary neuropathology is associated with subcortical and limbic volume loss

Next, we assessed how semi-quantitative primary pathology burden is associated with subcortical and limbic volumes across regions in their corresponding disease group (p-tau in AD and FTLD-Tau, α-synuclein in LBD, and TDP-43 in FTLD-TDP) with regression models with covariates (**Figure 5, Supplementary Figure S1.4, Supplementary Table S1.2**). In AD, we found that regional p-tau burden had significant negative correlations with volumes for both hippocampus (p=0.002) and putamen (p=0.002) after multiple comparison adjustments. In LBD, thalamic α-synuclein pathology correlated to volumes of thalamus, putamen and pallidum (p’s<0.02), after multiple test adjustment (**Supplementary Figure S1.5**). For FTLD-Tau, regional p-tau ratings had inverse correlations with hippocampus and amygdala volumes but did not withstand multiple comparison correction. For FTLD-TDP, TDP-43 severity had an inverse relationship with putamen volume but not after multiple test adjustment. Additionally, 3R and 4R tauopathy subgroups did not show significant correlations seen between semi-quantitative p-tau measures and subcortical/limbic volumes (**Supplementary Figure S2.3**).

**Figure 5.**
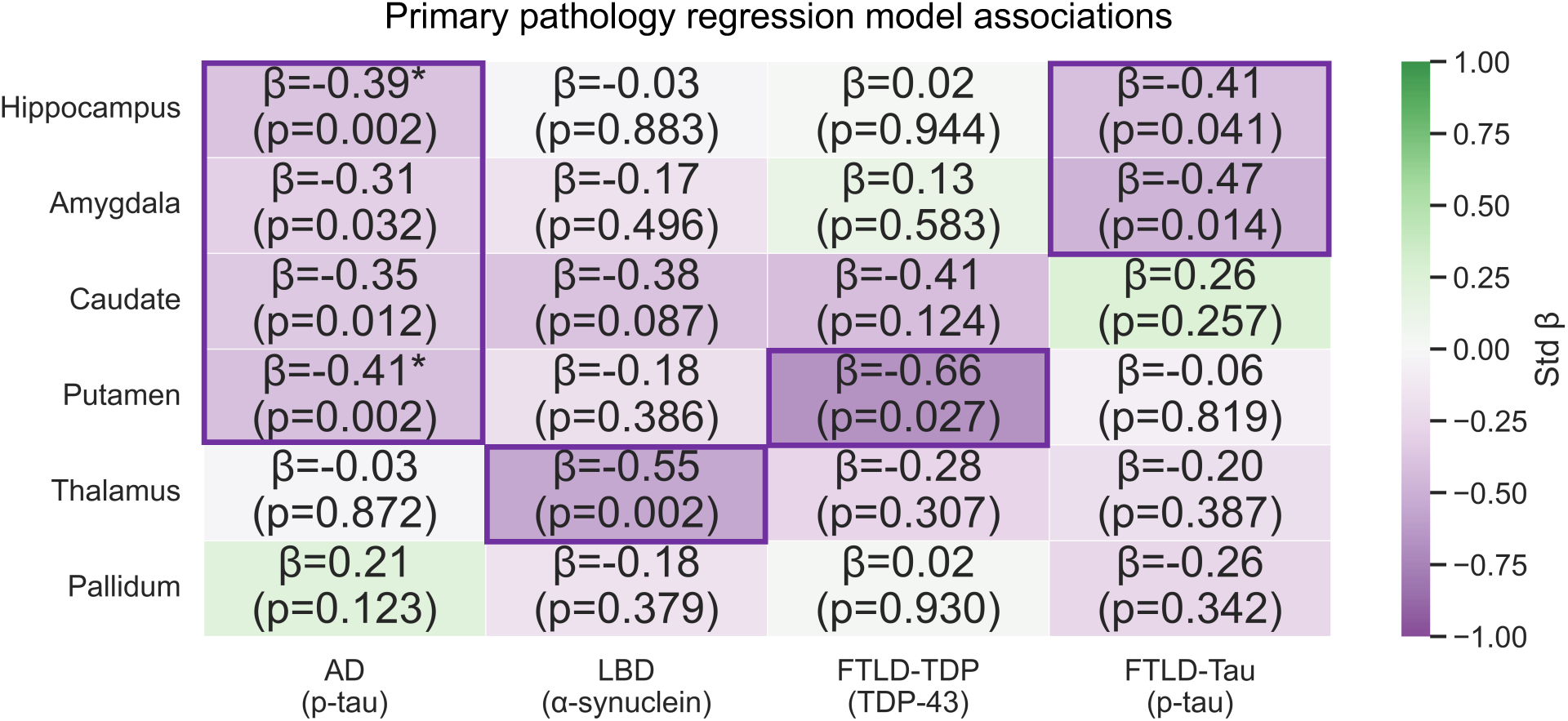
Primary pathology model with ordinary least squares model regression analysis between postmortem volumes and primary neuropathological burden. Heatmap shows standardized β coefficients from regression models on the relation between normalized postmortem subcortical and limbic matter volumes and corresponding regional predominant molecular pathology measures within each diagnostic group: AD (p-tau), LBD (α-synuclein), FTLD-TDP (TDP-43), and FTLD-tau (p-tau). Correlations were adjusted for age at death, sex, postmortem interval, and years of education. For the OLS regression analysis, an independent linear model was fit for each structure within each diagnostic group, including the region’s predominant pathology measure together with the same covariates and multiple comparisons adjustment. Each box displays standardized β coefficients from OLS models predicting each structure’s volume from its predominant regional pathology measure while adjusting for the same covariates with asterisks denoting significance after false discovery rate correction applied separately within each diagnostic group (*p<0.05 (and outlined), **p<0.01, ***p<0.001). Pathology burden ratings were not available for the nucleus accumbens. **Supplementary Table S1.2** tabulates detailed statistical results respectively.

### 3.6. Polypathologic models suggest primary pathology mostly drives subcortical and limbic volume loss

Approximately 82% (108/132) of donors had secondary neuropathological diagnoses, so we assessed a polypathology regression with multiple pathologies (tau, α-synuclein and TDP-43) for each primary diagnostic group [3], also correcting for covariates (**Figure 6, Supplementary Table S1.3**). For the AD polypathology models, hippocampal p-tau pathology had significant correlations with hippocampal volume (p=0.007), which did not remain after multiple comparison adjustments. Donors with AD also exhibited partial correlations between amygdala volumes and TDP-43 burden that did not survive multiple test corrections. Regional p-tau burden associated with hippocampal and amygdala volume in FTLD-Tau and regional α-synuclein pathology associated with thalamic volume in LBD, though these correlations did not withstand multiple test adjustments. Similarly to results from the primary pathology models, for FTLD-TDP, putamen TDP-43 severity and volume had a trend that did not survive multiple test adjustment. Distinguishing between 3R and 4R tauopathy in the polypathology model highlighted trends in lower pallidum volume with greater regional p-tau and α-synuclein involvement, though these did not remain after multiple test correction (**Supplementary Figure S2.4**). These findings remain similar in polypathology models with additional covariates of neuronal loss and gliosis (**Supplementary Figure S1.6**).

**Figure 6.**
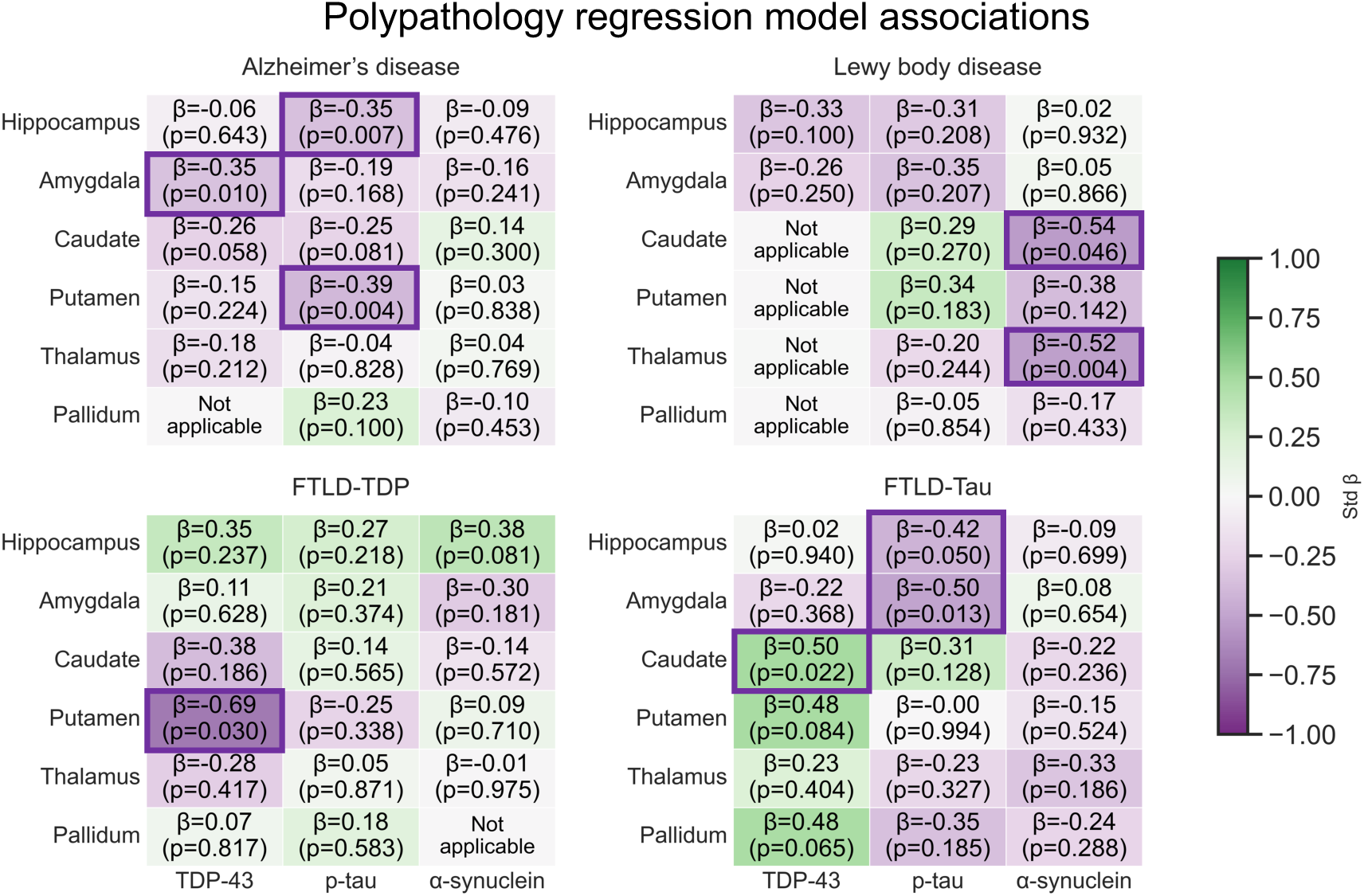
Polypathology ordinary least squares models for structure-pathology relationships. Heatmaps show standardized β coefficients from OLS models quantifying the association between regional postmortem MRI volumes (rows) and multiple semi-quantitative pathology ratings (columns) across the four diagnostic groups (Alzheimer’s disease, Lewy body disease, FTLD-TDP, FTLD-Tau). Each cell reports the standardized effect size (β) and the corresponding uncorrected p value; boxes highlight uncorrected p<0.05, and asterisks denote significance after false discovery rate correction using the Benjamini–Hochberg procedure applied separately within each diagnostic group across all structures and pathology predictors (*p<0.05 (also outlined), **p<0.01, ***p<0.001). All models included age at death, sex, years of education, and postmortem interval as covariates. Pathology burden ratings were not available for the nucleus accumbens. **Supplementary Table S1.3** shows detailed statistical results. <u>Note:</u> some predictors appear as “Not applicable” because several histology markers exhibited minimal or zero variability within disease groups. Predictors lacking sufficient within-group variance cannot yield interpretable regression estimates and may otherwise produce β values near zero with spuriously small p-values. Such predictors were removed prior to model fitting and excluded from FDR correction.

### 3.7. Histologic neuronal loss and gliosis mediate relationships between subcortical and limbic pathology and volume loss

Because neurodegeneration is classically associated with neuron loss and gliosis [27], we compared gross subcortical and limbic volumes with semi-quantitative histologic measures of gliosis and neuronal loss accounting for covariates (**Figure 7A, Supplementary Figure S1.7**). In AD, microscopic neuronal loss ratings significantly corresponded to macroscopic volume loss in amygdala (p=0.004), caudate (p=0.029) and putamen (p=0.020). Similarly in AD, local gliosis scores correlated to volume loss in amygdala (p=0.025). In FTLD-Tau, significant associations were found between microscopic neuronal loss and volume loss in hippocampus (p<0.001) and (p<0.001). In parallel, for FTLD-Tau, gliosis was associated with volume loss in hippocampus (p<0.001) and amygdala (p<0.001). This finding did not remain after splitting FTLD-Tau into 3R and 4R tauopathy subgroups (**Supplementary Figure S2.5**). Donors with LBD and FTLD-TDP had no significant relationships between these microscopic and macroscopic measures.

**Figure 7.**
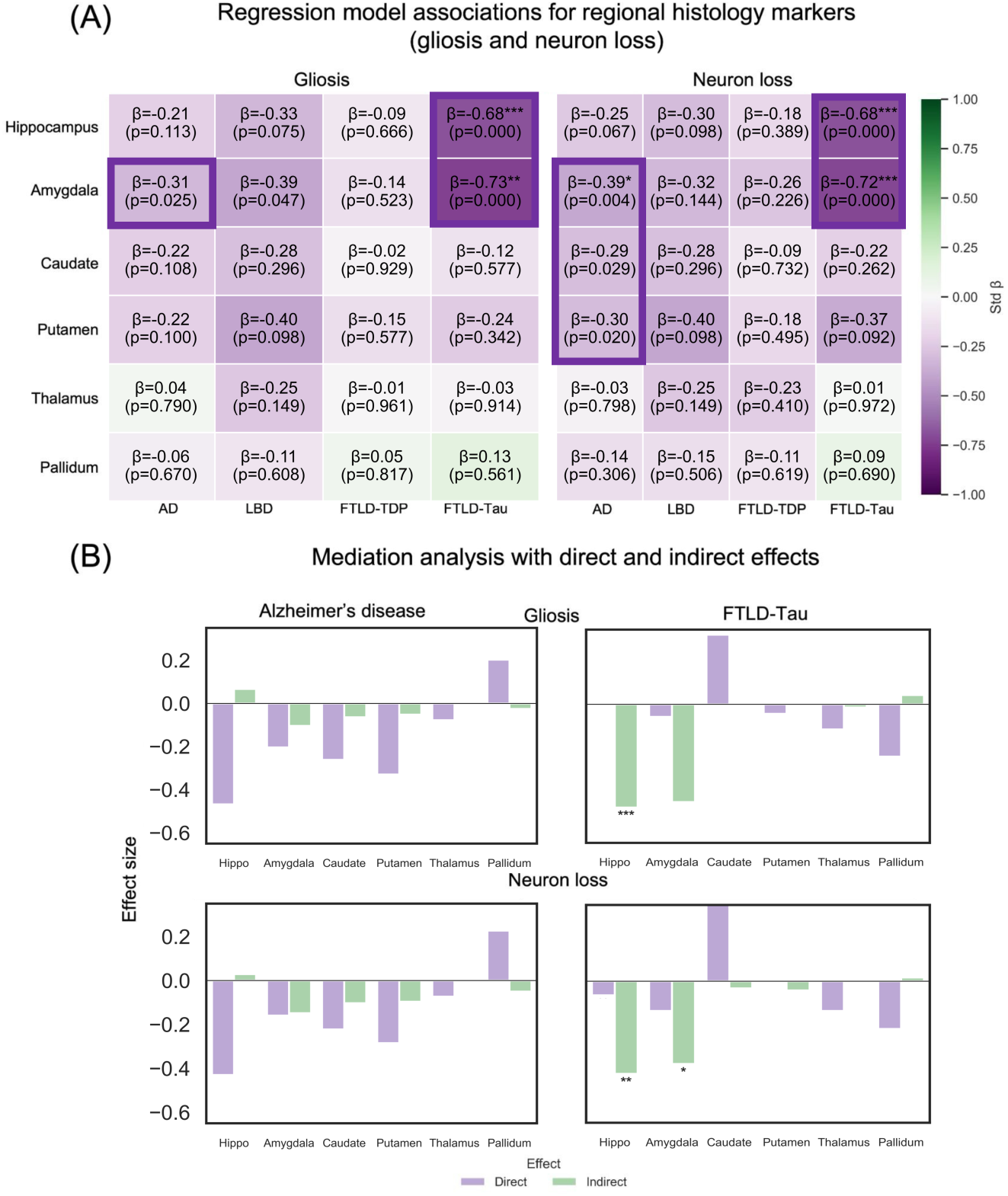
Structure-histology relationships and mediation analyses for gliosis and neuronal loss. (**A**) Correlations between postmortem MRI volumes and regional mediation markers were calculated between postmortem subcortical and limbic MRI volumes and semi-quantitative ratings of gliosis and neuronal loss for each group. All correlations were adjusted for age at death, sex, postmortem interval and education. Heatmap colors reflect slope (β) with corresponding uncorrected p value (in parentheses, boxes for p<0.05) with asterisks denoting significance after FDR correction applied within group (*p<0.05, **p<0.01, ***p<0.001). (**B**) Mediation analysis with the bar plots showing effect sizes for *direct* (purple) and *indirect* (green) effects from regional pathology burden to subcortical/limbic volume, mediated by gliosis (top row) or neuronal loss (bottom row). Pathology burden ratings were not available for the nucleus accumbens, so this structure is not shown. **Supplementary Tables S1.4** and **S1.5** tabulates detailed statistical results respectively.

Mediation analyses show both direct and indirect relationships between subcortical/limbic neuropathology and volume loss, mediated in part by neuronal loss and gliosis, even after adjusting for covariates and multiple comparisons (**Figure 7B**). For FTLD-Tau, the indirect associations between local p-tau pathology and volume loss were mediated by hippocampal neuronal loss (p<0.001), hippocampal gliosis (p<0.001) and amygdala neuronal loss (p<0.05).

### 3.8. Cerebrovascular pathology is not associated with subcortical and limbic volume loss

Given that subcortical regions are common sites of ischemic injury, we investigated how microvascular features may relate to deep brain atrophy. Vascular pathology, as measured by global cerebral amyloid angiopathy, atherosclerosis and arteriolosclerosis scores, did not significantly correlate with subcortical or limbic volumes in groups after multiple test correction (**Supplementary Figures S1.4, S2.6**).

## 4. Discussion

We examined structure-structure and structure-pathology relationships between subcortical, limbic and cortical regions and p-tau, a-synuclein, and TDP-34 neuropathologies by incorporating postmortem imaging and histopathology. With our novel pipeline, we found that brain donors with FTLD-Tau and FTLD-TDP had lower postmortem volumes across most subcortical and limbic structures than donors with either AD or LBD. Similarly, striatum and accumbens had lower volume in FTLD-TDP than in AD. Strikingly, AD had diffuse cortico-subcortical and cortico-limbic morphometric associations, while LBD had limited parieto-occipital cortico-limbic associations. FTLD-Tau exhibited frontal cortico-limbic and cortico-subcortical structural associations while FTLD-TDP only showed cortico-subcortical associations. These findings in FTLD-Tau remained significant after stratifying by 3R and 4R tauopathies. In AD and FTLD-Tau, hippocampal volume loss correlated with p-tau, neuronal loss and gliosis measures, but not for TDP-43 or α-synuclein. In LBD, regional α-synuclein was associated with thalamic volume, independent of other pathologies. In FTLD-TDP, local TDP-43 had no significant volume associations. Overall, our study demonstrates the utility of postmortem imaging to reveal relationships between deep brain structure and pathologies across neurodegenerative diseases.

A prior study applied postmortem 3T MRI with shape analysis and pathology in AD and LATE cohorts to uncover structure-pathology insights [11].We build on this work with ultra-high-resolution 7T MRI and deep learning segmentations of limbic, subcortical and cortical regions in a diverse cohort with AD, LBD, FTLD-TDP, and FTLD-Tau. Our cohort also compared copathologies, histologic measurements and genetic data in donors with sporadic and genetic disease.

Subcortical and limbic volume loss was significantly associated with diffuse cortical atrophy in AD and a more restricted atrophy pattern limited to frontal regions and supramarginal gyrus in FTLD-TDP and FTLD-Tau. This is consistent with prior studies suggesting that pathologies arise early in specific subcortical and limbic regions and spread through neural circuits to specific cortical regions in amyotrophic lateral sclerosis (ALS-TDP) and perhaps FTLD-TDP [3, 48, 49], but not necessarily in other cases such as FTLD-Tau [25, 59, 60, 61]. FTLD-TDP harbored cortico-subcortical associations, in keeping with evidence of subcortical-to-cortical spread, while FTLD-Tau portrayed both cortico-subcortical and cortico-limbic morphometric associations. Some of this lack of correlation in FTLD-TDP may be attributed to a floor effect in hippocampal volumes in FTLD-TDP compared to FTLD-Tau, the latter of which had greater variation to derive associations. Nevertheless, these findings suggest differing regional patterns involved in misfolded protein propagation specific to TDP-43 vs. p-tau. LBD demonstrated parieto-occipital cortico-limbic associations but not cortico-subcortical relationships. Since most of our LBD group had a secondary AD neuropathology (21/26), this cortico-limbic association is concordant with prior studies [39, 41]and could represent mixed AD with LBD pattern of atrophy.

We contrasted histopathological markers and subcortical/limbic volume loss with regression analyses for primary pathology and polypathology models. Power was limited in our polypathology analyses, leading to observed relationships between volume and primary pathology and trends for secondary pathologies that did not survive multiple comparison adjustments. Donors with AD had significant partial correlations of p-tau burden with volumes for hippocampus, amygdala, caudate and putamen after accounting for covariates. In LBD, thalamic α-synuclein pathology correlated to all subcortical and limbic volumes. For FTLD-Tau, regional p-tau burden did not correlate with subcortical or limbic volumes after multiple tests adjustment. For FTLD-TDP, TDP-43 severity scores did not significantly correlate with subcortical and limbic volumes. From modeling regressions of primary pathology and polypathology, the association between hippocampal structure and p-tau pathology was supported in AD, with a hint towards comorbid TDP-43 pathology contributing to atrophy. The observed relationship between hippocampal shrinkage and p-tau pathology in AD was mediated by hippocampal neuronal loss and gliosis, but not atherosclerosis or arteriolosclerosis. Although many donors had cerebrovascular pathology, vascular disease may reflect a pathological factor that is not differentially associated with unique subcortical volume loss patterns. These findings support a strong tandem relationship between p-tau burden and cellular injury (neuron loss, gliosis and gross volume) [47, 51, 54, 62, 63], though not be as clear-cut for α-synuclein.

Compared to tau-related diseases, LBD had less subcortical and limbic volume loss and less robust relationships between cortical and subcortical neurodegeneration. Moreover, thalamic α-synuclein pathology in LBD correlated with subcortical and limbic volumes in primary pathology models, though this finding did not extend to the polypathology model after multiple test adjustment. Indeed, in donors with both AD and LBD pathology, those with primary LBD and secondary AD diagnoses had less atrophy than those with primary AD and secondary LBD diagnoses, across limbic, subcortical and cortical regions (except for caudate, globus pallidus, calcarine cortex and a few additional areas). The presence of neuronal α-synuclein pathology is associated with hypometabolism and neurotransmission changes but not much atrophy [16, 18, 39, 41, 50]. This suggests that LBD has less subcortical and limbic volume loss than other diseases, and/or non-LBD pathology may contribute to volume loss more than α-synuclein does in LBD [17, 53, 64], though we did not find specific evidence of this in our dataset. Evaluation of the effects of LBD pathology on subcortical structures may require different imaging tools, such as metabolism or synaptic density, as well as different queries of α-synuclein regional patterns [65].

The lack of significant partial correlations between TDP-43 burden and limbic/subcortical volume in FTLD-TDP or other groups has several explanations. First, this group may be underpowered. Secondly, similar studies demonstrate low magnitude correlations between TDP-43 severity and cortical region thickness [8, 33]. However, some groups find moderate associations between TDP-43 burden and volume loss [19, 20, 21, 23]. TDP-43 load may be less associated with cortical and limbic volume loss in primary pathology in FTLD-TDP than in limbic-predominant age-related TDP-43 encephalopathy (LATE) or comorbid AD with

LATE [52]. While the lack of evidence may be due to the absence of biological effects, additional technical sources can contribute, such as sampling error. This is relevant for TDP-43 pathology, which has low dynamic range and is sparse even in advanced stages. To this point, intraneuronal TDP-43 inclusions disappear once affected neurons die, unlike ghost tau tangles that remain after the neuron disintegrates.

The study had several limitations and next steps. Although the cohort included 132 donors, some groups were likely underpowered, particularly FTLD-TDP and polypathology models in which volume-pathology relationships were subdivided by diagnosis. This reduced our ability to detect additional polypathology associations that prior studies tested in omnibus cohorts [3]. Pathology and imaging were obtained from different hemispheres, a common practice in postmortem studies [6, 12, 33, 34, 35, 66], because many neuropathologies are bilateral, though not universally so [19, 67]. In AD, prior work found largely similar MTL thickness correlations with ipsilateral and contralateral p-tau scores [8]. Any unmeasured hemispheric asymmetry in pathology or volume would reduce power, likely most affecting FTLD-Tau. Another limitation is reliance on semi-quantitative pathology scores, which are subjective, variable across readers, and may not scale linearly with true burden. Cerebrovascular measures were whole-brain measures and may not capture local subcortical effects. We are now obtaining same-hemisphere immunohistochemistry and developing machine-learning–based quantitative ratings [46] to validate these findings. Monoaminergic nuclei are dysregulated in neurodegenerative disease [1], including basal forebrain [66, 68], substantia nigra [69], and locus coeruleus [70], so segmentation of additional subcortical regions is planned. Future work should examine more granular clinicopathological entities, including LATE [28, 30], FTLD-TDP type C with annexin A11 co-aggregates [45, 71],and genetic variants. We will investigate cortico-limbic versus cortico-subcortical associations in FTLD-Tau and FTLD-TDP and local shape deformations of subcortical and limbic structures.

Overall, our study provides a comprehensive postmortem framework for linking brain tissue structure with pathological markers of neurodegenerative disease across primary and secondary diagnoses, brain regions, and antemortem versus postmortem imaging. We find evidence of limbic, subcortical, and cortical volume loss in primary AD over primary LBD, even among donors with comorbid AD and LBD. We also demonstrate selective cortico-limbic associations in LBD and cortico-subcortical associations in FTLD-TDP, compared with more widespread associations in AD and, to a lesser extent, FTLD-Tau. Regional p-tau burden was associated with atrophy, partly mediated by neuron loss and gliosis. Together, these findings advance the characterization of the pathological basis of structural imaging biomarkers and neurodegenerative disease mechanisms.

## Acknowledgements

The authors thank the brain donors and their families for their generosity and support without which this scientific endeavor would not be possible. We thank the lab members of the Penn ADRD Trajectories, Copathology and Heterogeneity Lab, the Penn Image Computing Science Lab and the Penn Frontotemporal Degeneration Center for helpful discussions. This manuscript is dedicated to all patients with neurodegenerative disease and their families, including the grandparents of M.T.D.

## Consent Statement

Human brain specimens were obtained in accordance with local laws and regulations and include informed consent from next of kin at time of death or where possible, pre-consent during life.

## References

1. Ehrenberg AJ, Kelberman MA, Liu KY, et al. Priorities for research on neuromodulatory subcortical systems in Alzheimer’s disease: Position paper from the NSS PIA of ISTAART. Alzheimers Dement 2023;19(5):2182–2196.

2. van der Velpen IF, Vlasov V, Evans TE, et al. Subcortical brain structures and the risk of dementia in the Rotterdam Study. Alzheimers Dement 2023;19(2):646–657.

3. Phillips JS, Robinson JL, Cousins KAQ, et al. Polypathologic Associations with Gray Matter Atrophy in Neurodegenerative Disease. J Neurosci 2024;44(6):e0808232023.

4. Rosbergen MT, Van der Veere P, Claus JJ et al. Subcortical gray matter volumes and 5-year dementia risk in individuals with subjective cognitive decline or mild cognitive impairment: A multi-cohort analysis. Alzheimer’s & Dementia 2025;21(7):e70413.

5. Frigerio I, Boon BD, Lin CP, et al. Amyloid-β, p-tau and reactive microglia are pathological correlates of MRI cortical atrophy in Alzheimer’s disease. Brain communications 2021;3(4):fcab281.

6. Yushkevich PA, López MM, de Onzoño Martin MMI, et al. Three-dimensional mapping of neurofibrillary tangle burden in the human medial temporal lobe. Brain 2021;144(9):2784–2797.

7. Yushkevich PA, Ittyerah R, Li Y, et al. Morphometry of medial temporal lobe subregions using high-resolution T2-weighted MRI in ADNI3: Why, how, and what’s next? Alzheimers Dement 2024;20(11):8113–8128.

8. Ravikumar S, Wisse LEM, Lim S, et al. Ex vivo MRI atlas of the human medial temporal lobe: characterizing neurodegeneration due to tau pathology. Acta Neuropathol Commun 2021;9:173.

9. Wearn A, Tremblay SA, Tardif CL, et al. Neuromodulatory subcortical nucleus integrity is associated with white matter microstructure, tauopathy and APOE status. Nat Commun 2024;15(1):4706.

10. Banerjee A, Yang F, Dutta J, Cacciola A, et al. Atrophy patterns of deep gray matter nuclei in Alzheimer’s disease and frontotemporal dementia. J Alzheimers Dis 2025:13872877251390386.

11. Saifullah K, Makkinejad N, Yasar MT, et al. Neuropathological Correlates of Volume and Shape of Deep Gray Matter Structures in Community-Based Older Adults. Hum Brain Mapp 2025;46(10):e70273.

12. Salman Y, Goloubeva J, Huyghe L et al. Specific atrophy patterns distinguish tau and TDP-43 pathology: a longitudinal MRI ante-mortem study. Acta Neuropathologica Communications 2025: in press.

13. Wesseling A, Calandri IL, Bouwman MM, et al. Amygdalar and hippocampal volume loss in limbic-predominant age-related TDP-43 encephalopathy. Brain 2025;148(11):3913–3923.

14. Bouwman MM, Frigerio I, Lin CP, Reijner N, van de Berg WD, Jonkman LE Hippocampal subfields: volume, neuropathological vulnerability and cognitive decline in Alzheimer’s and Parkinson’s disease. Alzheimer’s Research & Therapy 2025;17(1):121.

15. Reijner N, Frigerio I, Bouwman MMA, et al. Clinical phenotypes of Alzheimer’s disease: investigating atrophy patterns and their pathological correlates. Alzheimer’s Research & Therapy 2025;17(1):93.

16. Surmeier DJ, Obeso JA, Halliday GM. Selective neuronal vulnerability in Parkinson disease. Nature Reviews Neuroscience 2017;18(2), pp.101–113.

17. Spotorno N, Coughlin DG, Olm CA, et al. Tau pathology associates with in vivo cortical thinning in Lewy body disorders. Annals of Clinical and Translational Neurology 2020;7(12):2342–2355.

18. Duong MT, Das SR, Khandelwal P, et al. Discordance of dopaminergic dysfunction and subcortical atrophy by α-synuclein status in sporadic and genetic Parkinson’s Disease. Movement Disorders 2025: in press.

19. Irwin DJ, McMillan CT, Xie SX, et al. Asymmetry of post-mortem neuropathology in behavioural-variant frontotemporal dementia. Brain 2018;141(1):288–301.

20. Giannini LA, Xie SX, McMillan CT, et al. Divergent patterns of TDP-43 and tau pathologies in primary progressive aphasia. Ann Neurol 2019;85(5), pp.630–643.

21. Yu, L, Boyle PA, Dawe RJ, Bennett DA, Arfanakis K, Schneider JA. Contribution of TDP and hippocampal sclerosis to hippocampal volume loss in older-old persons. Neurology 2020;94(2):e142–e152.

22. Bocchetta M, Malpetti M, Todd EG, Rowe JB, Rohrer JD. Looking beneath the surface: the importance of subcortical structures in frontotemporal dementia. Brain Commun 2021;3(3):fcab158.

23. Burke SE, Phillips JS, Olm CA, et al. Phases of volume loss in patients with known frontotemporal lobar degeneration spectrum pathology. Neurobiol Aging, 2022;113:95–107.

24. Dickson DW, Kouri N, Murray ME, Josephs KA. Neuropathology of frontotemporal lobar degeneration-tau (FTLD-tau). J Mol Neurosci 2011;45(3):384–389.

25. Irwin DJ, Brettschneider J, McMillan CT, et al. Deep clinical and neuropathological phenotyping of Pick disease. Ann Neurol 2016;79:272–287.

26. Miletić S, Bazin PL, Isherwood SJS, et al. Charting human subcortical maturation across the adult lifespan with in vivo 7 T MRI. Neuroimage 2022;249:118872.

27. Jack CR, Andrews JS, Beach TG, et al. Revised criteria for diagnosis and staging of Alzheimer’s disease: Alzheimer’s Association Workgroup. Alzheimer’s Dement 2024;20(8):5143–5169.

28. Wolk DA, Nelson PT, Apostolova L, et al. Clinical criteria for limbic-predominant age-related TDP-43 encephalopathy. Alzheimers Dement 2025;21(1):e14202.

29. Nichols E, Merrick R, Hay SI, et al. The prevalence, correlation, and co-occurrence of neuropathology in old age: harmonisation of 12 measures across six community-based autopsy studies of dementia. Lancet Healthy Longev 2023;4(3):e115–e125.

30. Nelson PT, Brayne C, Flanagan ME, et al. Frequency of LATE neuropathologic change across the spectrum of Alzheimer’s disease neuropathology: combined data from 13 community-based or population-based autopsy cohorts. Acta Neuropathol 2022;144:27–44.

31. Wisse LEM, Ravikumar S, Ittyerah R, et al. Downstream effects of polypathology on neurodegeneration of medial temporal lobe subregions. Acta Neuropathol Commun 2021;9(1):128.

32. Li K, Rashid T, Li J, et al. Postmortem Brain Imaging in Alzheimer’s Disease and Related Dementias: The South Texas Alzheimer’s Disease Research Center Repository. J Alzheimers Dis 2023;96(3):1267–1283.

33. Ravikumar S, Denning AE, Lim S, et al. Postmortem imaging reveals patterns of medial temporal lobe vulnerability to tau pathology in Alzheimer’s disease. Nat Commun 2024;15(1):4803.

34. Denning AE, Ittyerah R, Levorse LM, et al. Association of quantitative histopathology measurements with antemortem medial temporal lobe cortical thickness in the Alzheimer’s disease continuum. Acta Neuropathol 2024;148(1):37.

35. Khandelwal P, Duong MT, Sadaghiani S, et al. Automated deep learning segmentation of high-resolution 7 Tesla postmortem MRI for quantitative analysis of structure-pathology correlations in neurodegenerative diseases. Imaging Neurosci (Camb) 2024;2:1–30.

36. Khandelwal P, Duong MT, Levorse L, et al. Surface-Based Parcellation and Vertex-wise Analysis of Ultra High-resolution ex vivo 7 tesla MRI in Alzheimer’s disease and related dementias. In: Bathula, D.R., et al. Machine Learning in Clinical Neuroimaging. MLCN 2024. Lecture Notes in Computer Science, vol 15266. 2025. Springer, Cham.

37. Billot B, Greve DN, Puonti O, et al. SynthSeg: Segmentation of brain MRI scans of any contrast and resolution without retraining. Med Image Anal 2023;86:102789.

38. Liu P, Puonti O, Hu X, et al. A Modality-agnostic Multi-task Foundation Model for Human Brain Imaging. arXiv 2025;2509.00549.

39. Harper L, Bouwman F, Burton EJ, et al. Patterns of atrophy in pathologically confirmed dementias: a voxelwise analysis. J Neurol Neurosurg Psychiatry 2017;88(11):908–916.

40. Silva-Rodríguez J, Labrador-Espinosa MA, Moscoso A, et al. Characteristics of amnestic patients with hypometabolism patterns suggestive of Lewy body pathology. Brain 2023a;146(11):4520–4531.

41. Duong MT, Das SR, Khandelwal P, et al. Hypometabolic mismatch with atrophy and tau pathology in mixed Alzheimer’s and Lewy body disease. Brain 2024;148(5):1577–1587.

42. Toledo JB, Van Deerlin VM, Lee EB, et al. A platform for discovery: The University of Pennsylvania Integrated Neurodegenerative Disease Biobank. Alzheimers Dement 2014;10(4):477–484.e1.

43. Tisdall MD, Ohm DT, Lobrovich R, et al. Ex vivo MRI and histopathology detect novel iron-rich cortical inflammation in frontotemporal lobar degeneration with tau versus TDP-43 pathology. Neuroimage Clin 2022;33:102913.

44. Arezoumandan S, Xie SX, Cousins KAQ, et al. Regional distribution and maturation of tau pathology among phenotypic variants of Alzheimer’s disease. Acta Neuropathol 2022;144(6):1103–1116.

45. Arseni D, Nonaka T, Jacobsen MH, et al. Heteromeric amyloid filaments of ANXA11 and TDP-43 in FTLD-TDP Type C. Nature 2024;634(8034):662–668.

46. Athalye C, Bahena A, Khandewal P, et al. Operationalizing postmortem pathology-MRI association studies in Alzheimer’s disease and related disorders with MRI-guided histology sampling. Acta Neuropathol Commun 2025;13(1):120.

47. Bejanin A, Schonhaut DR, La Joie R, et al. Tau pathology and neurodegeneration contribute to cognitive impairment in Alzheimer’s disease. Brain 2017;140(12):3286–3300.

48. Braak H, Brettschneider J, Ludolph AC, Lee VM, Trojanowski JQ, Tredici KD. Amyotrophic lateral sclerosis—a model of corticofugal axonal spread. Nat Rev Neurol 2013;9(12):708–714.

49. Brettschneider J, Del Tredici K, Lee VM, Trojanowski JQ. Spreading of pathology in neurodegenerative diseases: a focus on human studies. Nat Rev Neurosci 2015;16(2):109–20.

50. Brookner SM, Pasquini J, Choi SH, et al. Clinical and imaging characteristics of Parkinson’s disease with negative alpha-synuclein seed amplification assay. medRxiv 2025.

51. Brown C, Das S, Xie L, et al. Medial temporal lobe gray matter microstructure in preclinical Alzheimer’s disease. Alzheimers Dement 2024;20(6):4147–4158.

52. Buciuc M, Martin PR, Tosakulwong N, et al. TDP-43-associated atrophy in brains with and without frontotemporal lobar degeneration. Neuroimage Clin 2022;34:102954.

53. Chu Y, Hirst WD, Federoff HJ, Harms AS, Stoessl AJ, Kordower JH. Nigrostriatal tau pathology in parkinsonism and Parkinson’s disease. Brain 2024;147(2):444–457.

54. Das SR, Lyu X, Duong MT, et al. Tau-Atrophy Variability Reveals Phenotypic Heterogeneity in Alzheimer’s Disease. Ann Neurol 2021;90(5):751–762.

55. Desikan RS, Ségonne F, Fischl B, et al. An automated labeling system for subdividing the human cerebral cortex on MRI scans into gyral based regions of interest. Neuroimage 2006;31(3):968–980.

56. Duong MT, Das SR, Lyu X, et al. Dissociation of tau pathology and neuronal hypometabolism within the ATN framework of Alzheimer’s disease. Nat Commun 2022;13(1):1495.

57. Edlow BL, Mareyam A, Horn A, et al. 7 Tesla MRI of the ex vivo human brain at 100 micron resolution. Sci Data 2019;6(1):244.

58. Seabold, Skipper, and Josef Perktold. “Statsmodels: econometric and statistical modeling with python.” SciPy 7.1. 2010:92–96.

59. Kovacs GG, Lukic MJ, Irwin DJ, et al. Distribution patterns of tau pathology in progressive supranuclear palsy. Acta Neuropathol 2020;140(2):99–119.

60. Ling H, Kovacs GG, Vonsattel JPG, et al. Astrogliopathy predominates the earliest stage of corticobasal degeneration pathology. Brain 2016;139(12):3237–3252.

61. Giannini LA, Ohm DT, Rozemuller AJ, et al. Isoform-specific patterns of tau burden and neuronal degeneration in MAPT-associated frontotemporal lobar degeneration. Acta Neuropathol 2022;144(6):1065–1084.

62. Gómez-Isla T, Hollister R, West H, et al. Neuronal loss correlates with but exceeds neurofibrillary tangles in Alzheimer’s disease. Ann Neurol 1997;41(1):17–24.

63. Lyu X, Duong MT, Xie L, et al. Tau-neurodegeneration mismatch reveals vulnerability and resilience to comorbidities in Alzheimer’s continuum. Alzheimers Dement 2023;20(3):1586–1600.

64. Popescu A, Lippa CF, Lee VM, Trojanowski JQ. Lewy bodies in the amygdala: increase of alpha-synuclein aggregates in neurodegenerative diseases with tau-based inclusions. Arch Neurol 2004;61(12):1915–9.

65. Mastenbroek SE, Vogel JW, Collij LE, et al. Disease progression modelling reveals heterogeneity in trajectories of Lewy-type α-synuclein pathology. Nature Commun 2024;15(1):5133.

66. Wisse LEM, Spotorno N, Rossi, et al. MRI Signature of α-Synuclein Pathology in Asymptomatic Stages and a Memory Clinic Population. JAMA Neurol 2024;81(10):1051–1059.

67. Tremblay C, Serrano GE, Intorcia AJ, et al. Hemispheric asymmetry and atypical lobar progression of Alzheimer-type tauopathy. J Neuropathol & Exp Neurol 2022;81(3):158–171.

68. Labrador-Espinosa MA, Silva-Rodriguez J, et al. Cortical hypometabolism in Parkinson’s disease is linked to cholinergic basal forebrain atrophy. Mol Psychiatry 2025;30(6):2372–2380.

69. Silva-Rodríguez J, Labrador-Espinosa MA, Moscoso Aet al. Differential effects of tau stage, Lewy body pathology, and substantia nigra degeneration on FDG-PET patterns in clinical Alzheimer disease. J Nucl Med 2023b;64:274–280.

70. Freeze WM, Van Veluw SJ, Jansen WJ, Bennett DA, Jacobs HII. Locus coeruleus pathology is associated with cerebral microangiopathy at autopsy. Alzheimers Dement 2023;19(11):5023–5035.

71. Robinson JL, Suh E, Xu Y, et al. Annexin A11 aggregation in FTLD-TDP type C and related neurodegenerative disease proteinopathies. Acta Neuropathol 2024;147(1):104.

72. Fischl B. FreeSurfer. Neuroimage 2012;62.2:774–781.

