## Supplementary File 1 for "Postmortem brain MRI reveals differential associations of subcortical and limbic volumes with cortical thinning and neurodegenerative pathologies"

### Supplementary File 1: Four disease diagnostic groups

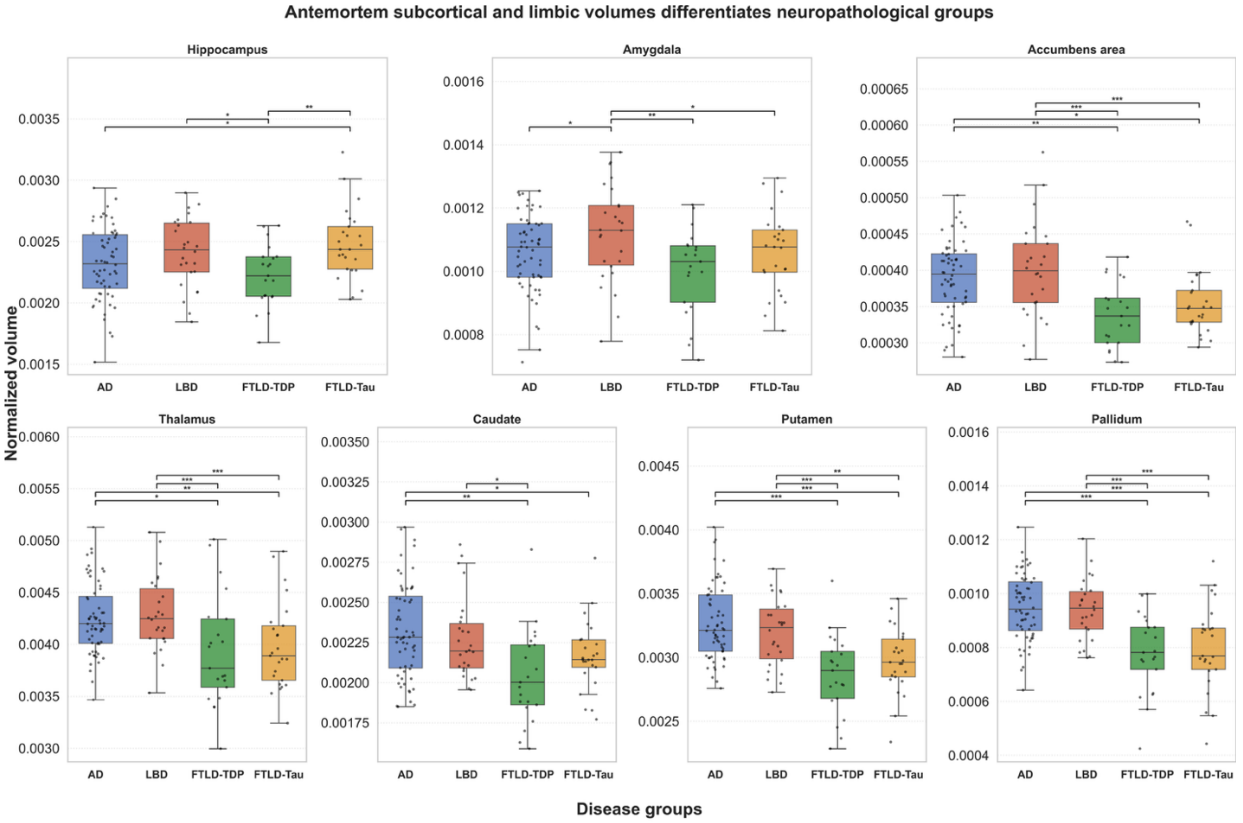


**Supplementary Figure S1.1. <u>Antemortem</u> limbic and subcortical volumes across neuropathological groups.** Boxplots show ICV-normalized antemortem volumes (mean of the two hemispheres) for the seven subcortical and limbic regions across four primary diagnostic categories: Alzheimer's disease (AD), Lewy body disease (LBD), Frontotemporal Lobar Degeneration with TDP-43 pathology (FTLD-TDP), and FTLD-tau. Pairwise differences between groups were assessed using likelihood-ratio tests adjusting for age at death, sex, education, and <u>antemortem</u> interval. Statistically significant pairwise comparisons with multiple test adjustment are denoted with *p<0.05, **p<0.01, ***p<0.001. In each boxplot, the center line indicates median, box edges represent 25th–75th percentiles (IQR), whiskers extend to 1.5×IQR, and overlaid points correspond to individuals.

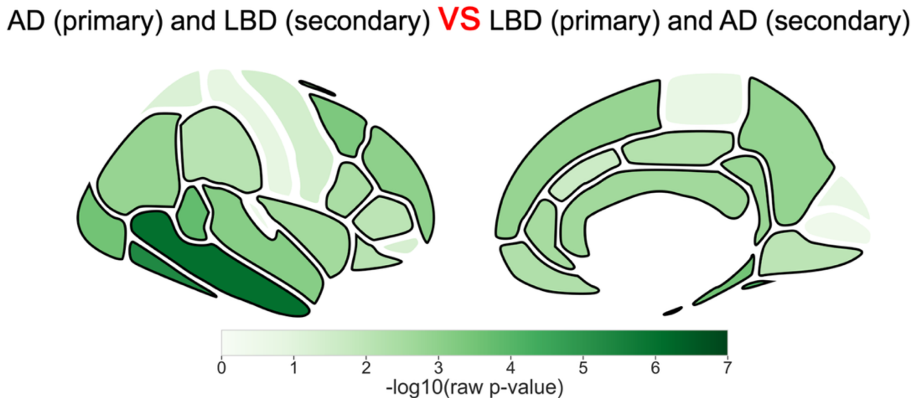


**Supplementary Figure S1.2. Postmortem regional mean cortical thickness across major neuropathological groups with both AD and LBD pathologies.** Mean postmortem cortical thickness for DKT atlas-based cortical regions across primary AD and secondary LBD (n = 29) vs. primary LBD and secondary AD (n = 21) in donors with both AD and LBD pathologies were assessed using likelihood-ratio tests adjusting for age at death, sex, education, and postmortem interval. Shown are the $-\log_{10}(p)$ for an easier interpretation. Statistically significant pairwise comparisons with multiple test adjustment are denoted with regions outlined in black indicating $p<0.05$.

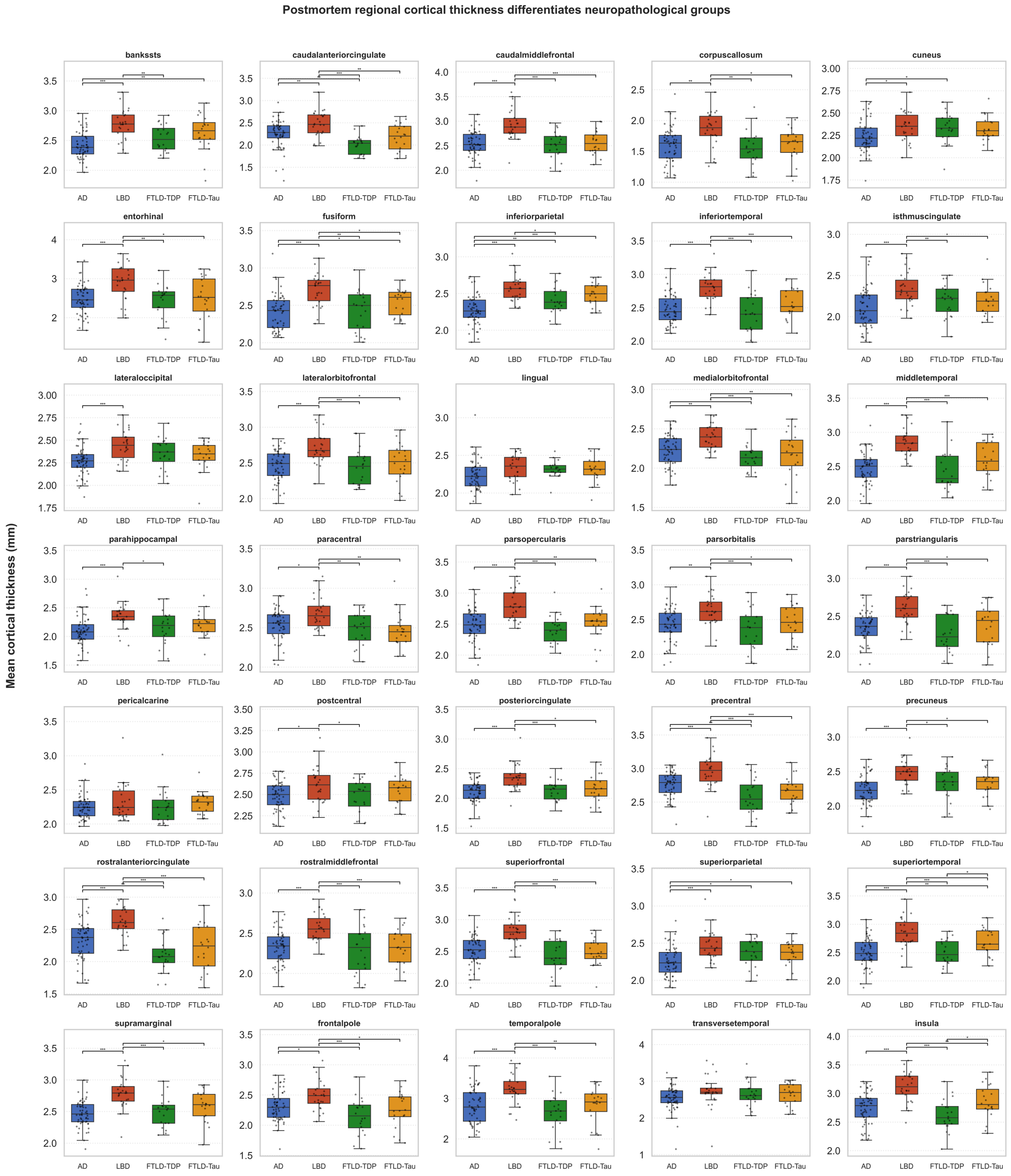


**Supplementary Figure S1.3. Postmortem regional mean cortical thickness across major neuropathological groups.** Boxplots show mean postmortem cortical thickness for DKT atlas-based cortical regions across four primary diagnostic categories: Alzheimer's disease (AD), Lewy body disease (LBD), Frontotemporal Lobar Degeneration with TDP-43 pathology (FTLD-TDP), and FTLD-tau. Pairwise differences between groups were assessed using likelihood-ratio tests adjusting for age at death, sex, education, and postmortem interval. Statistically significant pairwise comparisons with multiple test adjustment are denoted with *p<0.05, **p<0.01, ***p<0.001. In each boxplot, the center line indicates median, box edges represent 25th–75th percentiles (IQR), whiskers extend to 1.5×IQR, and overlaid points correspond to individuals.

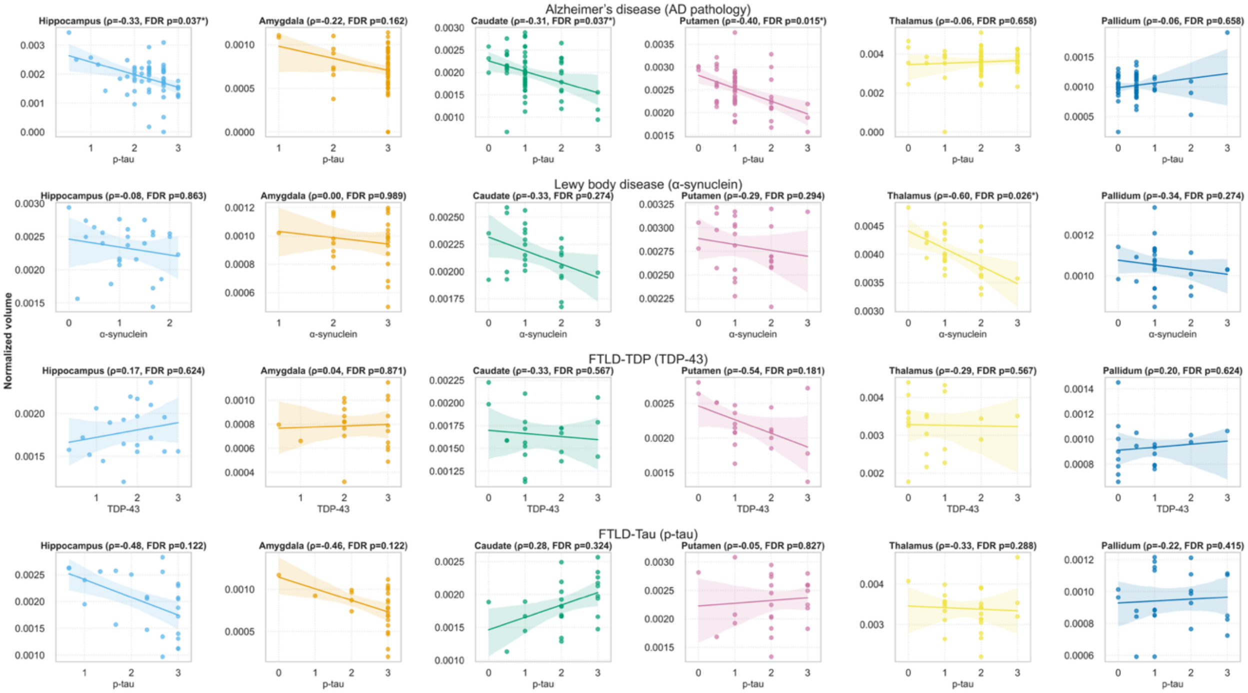


**Supplementary Figure S1.4. Partial Spearman correlations between regional postmortem MRI volumes and primary regional pathology burden across major neuropathological groups.** Scatter plots show covariate-adjusted partial Spearman correlations between ICV-normalized postmortem subcortical/limbic volumes and semi-quantitative pathology scores within each disease group: Alzheimer's disease (p-tau), Lewy body disease (α-synuclein), FTLD-TDP (TDP-43), and FTLD-Tau (p-tau). Each row corresponds to one diagnostic group, and each column to a distinct subcortical/limbic structure (hippocampus, amygdala, caudate, putamen, thalamus, and pallidum). Regression lines depict the direction of association, with raw p-values and significance levels after false-discovery-rate (FDR) correction shown in the subplot titles. All correlations were adjusted for age at death, sex, postmortem interval (PMI), and education. Pathology burden ratings were not available for the nucleus accumbens, and therefore this structure is not shown.

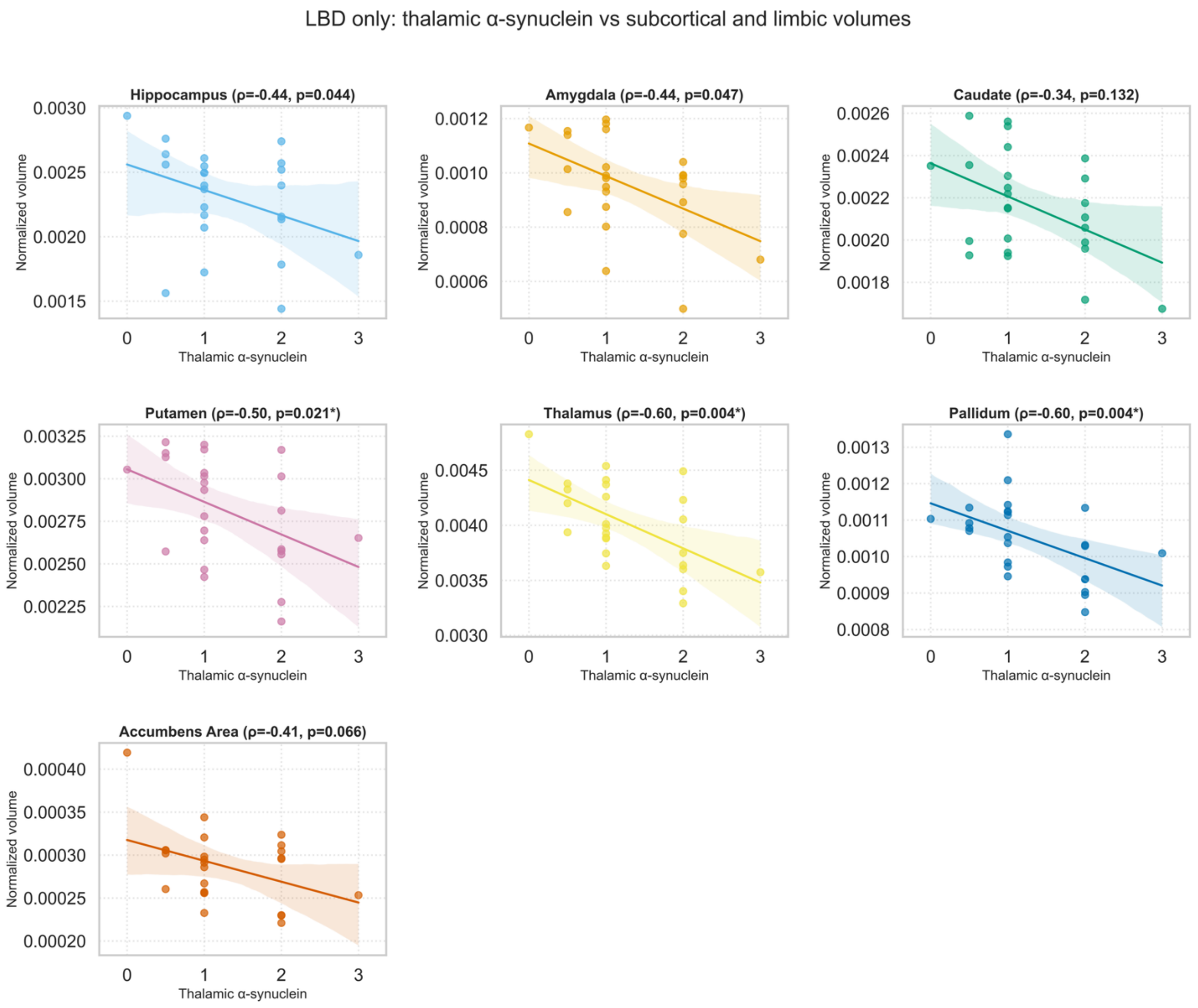


**Supplementary Figure S1.5. Thalamic α-synuclein severity correlates with subcortical volume loss in Lewy body disease.** Scatter plots show covariate-adjusted partial Spearman correlations between ICV-normalized postmortem subcortical/limbic volumes in LBD.

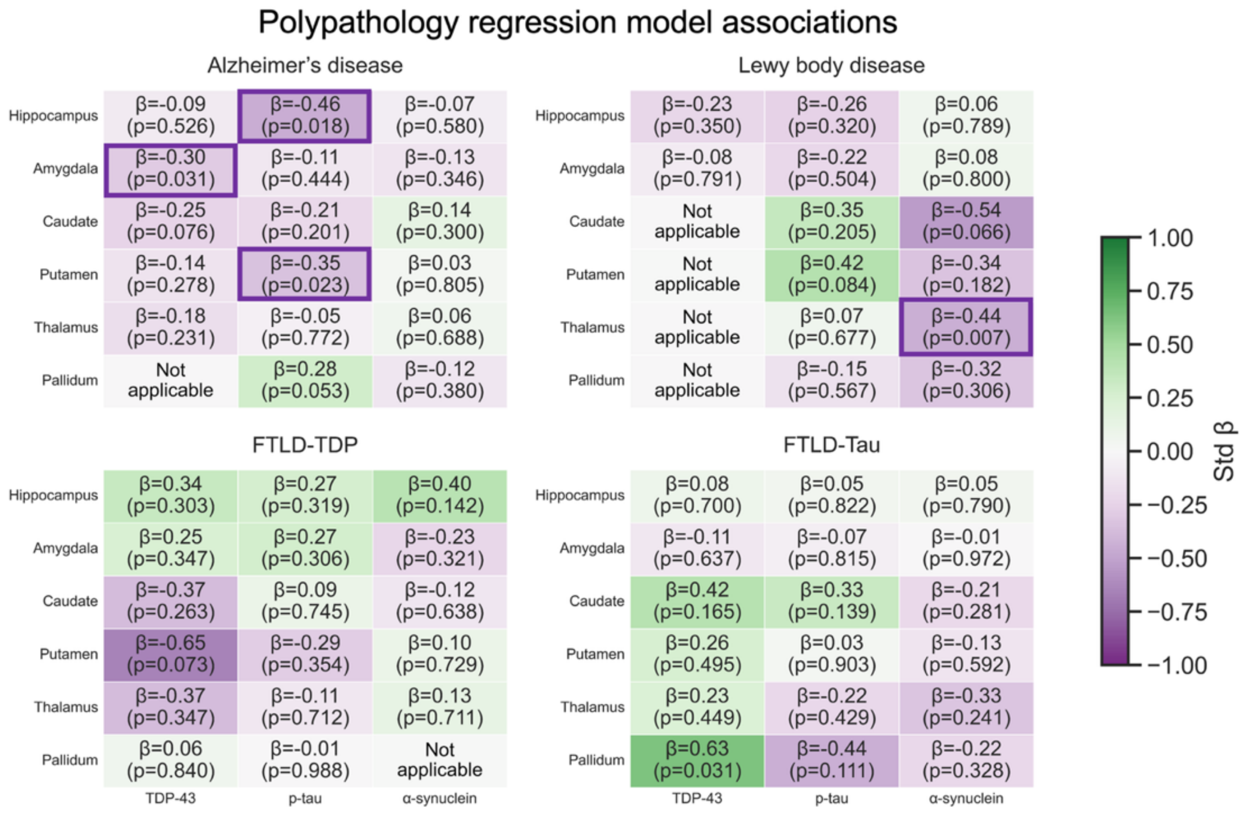


**Supplementary Figure S1.6. Polypathology ordinary least squares models for structure-pathology relationships.** Heatmaps show standardized β coefficients from OLS models quantifying the association between regional postmortem MRI volumes (rows) and multiple semi-quantitative pathology ratings (columns) across the four diagnostic groups (Alzheimer's disease, Lewy body disease, FTLD-TDP, FTLD-Tau). Each cell reports the standardized effect size (β) and the corresponding uncorrected p value; boxes highlight uncorrected p<0.05, and asterisks denote significance after false discovery rate correction using the Benjamini-Hochberg procedure applied separately within each diagnostic group across all structures and pathology predictors (*p<0.05 (also outlined), **p<0.01, ***p<0.001). All models included age at death, sex, years of education, postmortem interval, **gliosis and neuronal loss** as covariates. Pathology burden ratings were not available for the nucleus accumbens. Note: some predictors appear as "Not applicable" because several histology markers exhibited minimal or zero variability within disease groups. Predictors lacking sufficient within-group variance cannot yield interpretable regression estimates and may otherwise produce β values near zero with spuriously small p-values. Such predictors were removed prior to model fitting and excluded from FDR correction.

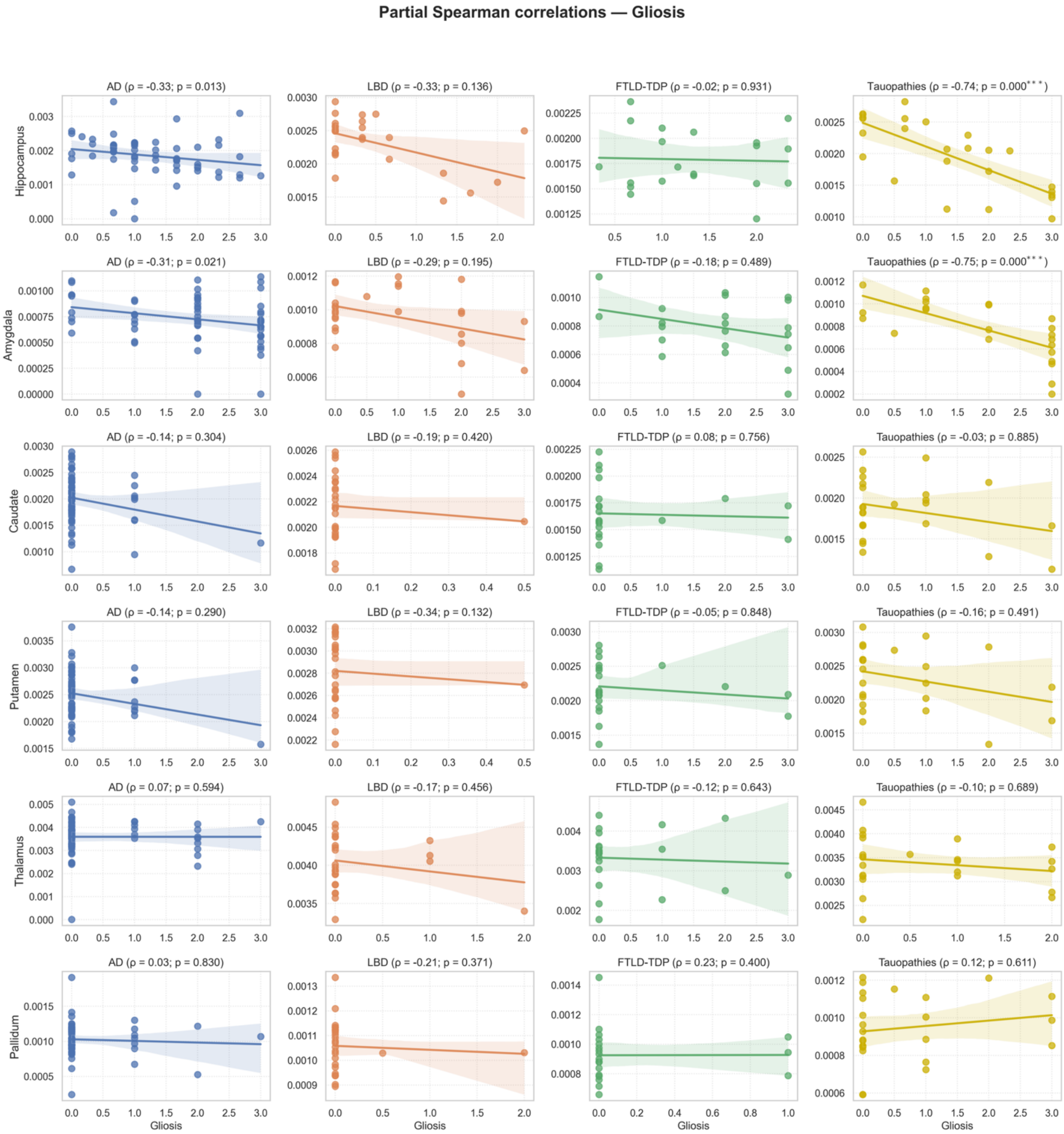
Partial Spearman correlations — Gliosis
AD (ρ = -0.33; p = 0.013)
LBD (ρ = -0.33; p = 0.136)
FTLD-TDP (ρ = -0.02; p = 0.931)
Tauopathies (ρ = -0.74; p = 0.000***)
Hippocampus
AD (ρ = -0.31; p = 0.021)
LBD (ρ = -0.29; p = 0.195)
FTLD-TDP (ρ = -0.18; p = 0.489)
Tauopathies (ρ = -0.75; p = 0.000***)
Amygdala
AD (ρ = -0.14; p = 0.304)
LBD (ρ = -0.19; p = 0.420)
FTLD-TDP (ρ = 0.08; p = 0.756)
Tauopathies (ρ = -0.03; p = 0.885)
Caudate
AD (ρ = -0.14; p = 0.290)
LBD (ρ = -0.34; p = 0.132)
FTLD-TDP (ρ = -0.05; p = 0.848)
Tauopathies (ρ = -0.16; p = 0.491)
Putamen
AD (ρ = 0.07; p = 0.594)
LBD (ρ = -0.17; p = 0.456)
FTLD-TDP (ρ = -0.12; p = 0.643)
Tauopathies (ρ = -0.10; p = 0.689)
Thalamus
AD (ρ = 0.03; p = 0.830)
LBD (ρ = -0.21; p = 0.371)
FTLD-TDP (ρ = 0.23; p = 0.400)
Tauopathies (ρ = 0.12; p = 0.611)
Pallidum
Gliosis
Gliosis
Gliosis
Gliosis

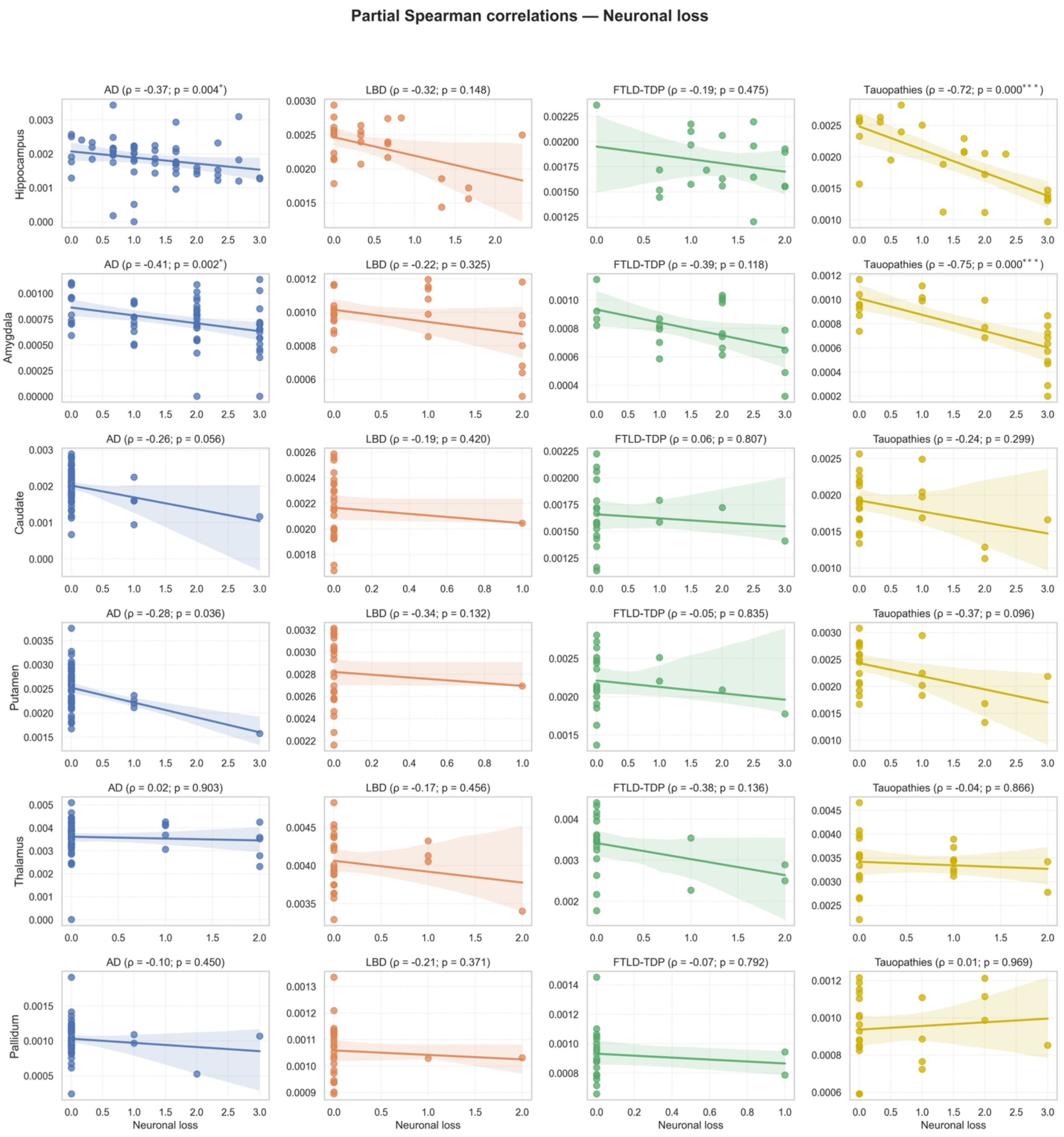


**Supplementary Figure S1.7. Structure-histology relationships for gliosis and neuronal loss.** Partial Spearman correlation between postmortem MRI volumes and regional mediation markers were calculated between postmortem subcortical/limbic volumes and semi-quantitative ratings of regional gliosis and neuronal loss for each group. All correlations were adjusted for age at death, sex, postmortem interval and education. Shown are the correlation strength (ρ) with the corresponding uncorrected p-value (in parentheses, boxes for $p<0.05$) with asterisks denoting significance after false discovery rate (FDR) correction applied within group across all structures (*$p<0.05$, **$p<0.01$, ***$p<0.001$). Pathology burden ratings were not available for the nucleus accumbens, and therefore this structure is not shown.

(A)

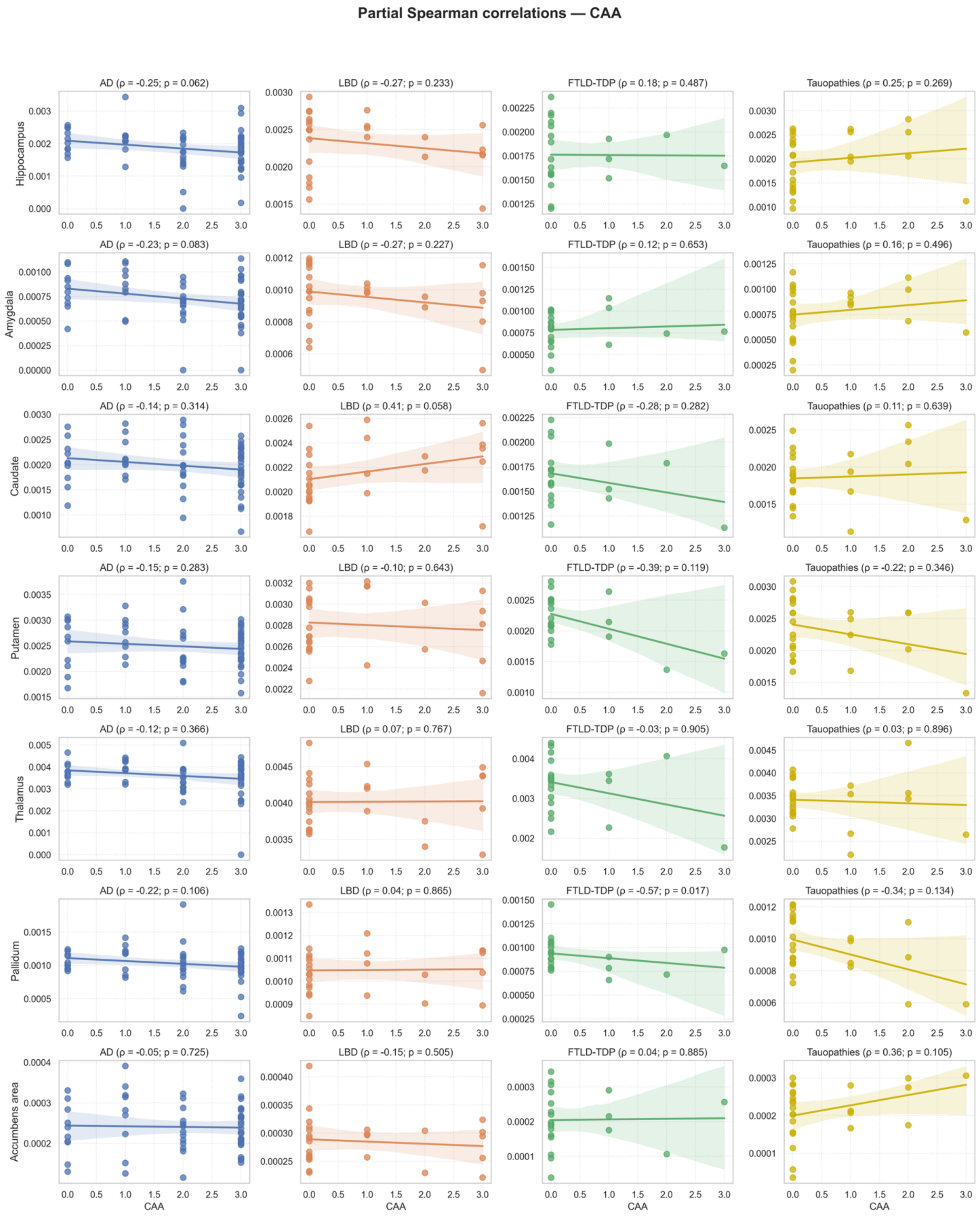
Partial Spearman correlations — CAA
AD (ρ = -0.25; p = 0.062)
LBD (ρ = -0.27; p = 0.233)
FTLD-TDP (ρ = 0.18; p = 0.487)
Tauopathies (ρ = 0.25; p = 0.269)
Hippocampus
AD (ρ = -0.23; p = 0.083)
LBD (ρ = -0.27; p = 0.227)
FTLD-TDP (ρ = 0.12; p = 0.653)
Tauopathies (ρ = 0.16; p = 0.496)
Amygdala
AD (ρ = -0.14; p = 0.314)
LBD (ρ = 0.41; p = 0.058)
FTLD-TDP (ρ = -0.28; p = 0.282)
Tauopathies (ρ = 0.11; p = 0.639)
Caudate
AD (ρ = -0.15; p = 0.283)
LBD (ρ = -0.10; p = 0.643)
FTLD-TDP (ρ = -0.39; p = 0.119)
Tauopathies (ρ = -0.22; p = 0.346)
Putamen
AD (ρ = -0.12; p = 0.366)
LBD (ρ = 0.07; p = 0.767)
FTLD-TDP (ρ = -0.03; p = 0.905)
Tauopathies (ρ = 0.03; p = 0.896)
Thalamus
AD (ρ = -0.22; p = 0.106)
LBD (ρ = 0.04; p = 0.865)
FTLD-TDP (ρ = -0.57; p = 0.017)
Tauopathies (ρ = -0.34; p = 0.134)
Pallidum
AD (ρ = -0.05; p = 0.725)
LBD (ρ = -0.15; p = 0.505)
FTLD-TDP (ρ = 0.04; p = 0.885)
Tauopathies (ρ = 0.36; p = 0.105)
Accumbens area
CAA
CAA
CAA
CAA

**Partial Spearman correlations — Arteriolosclerosis**

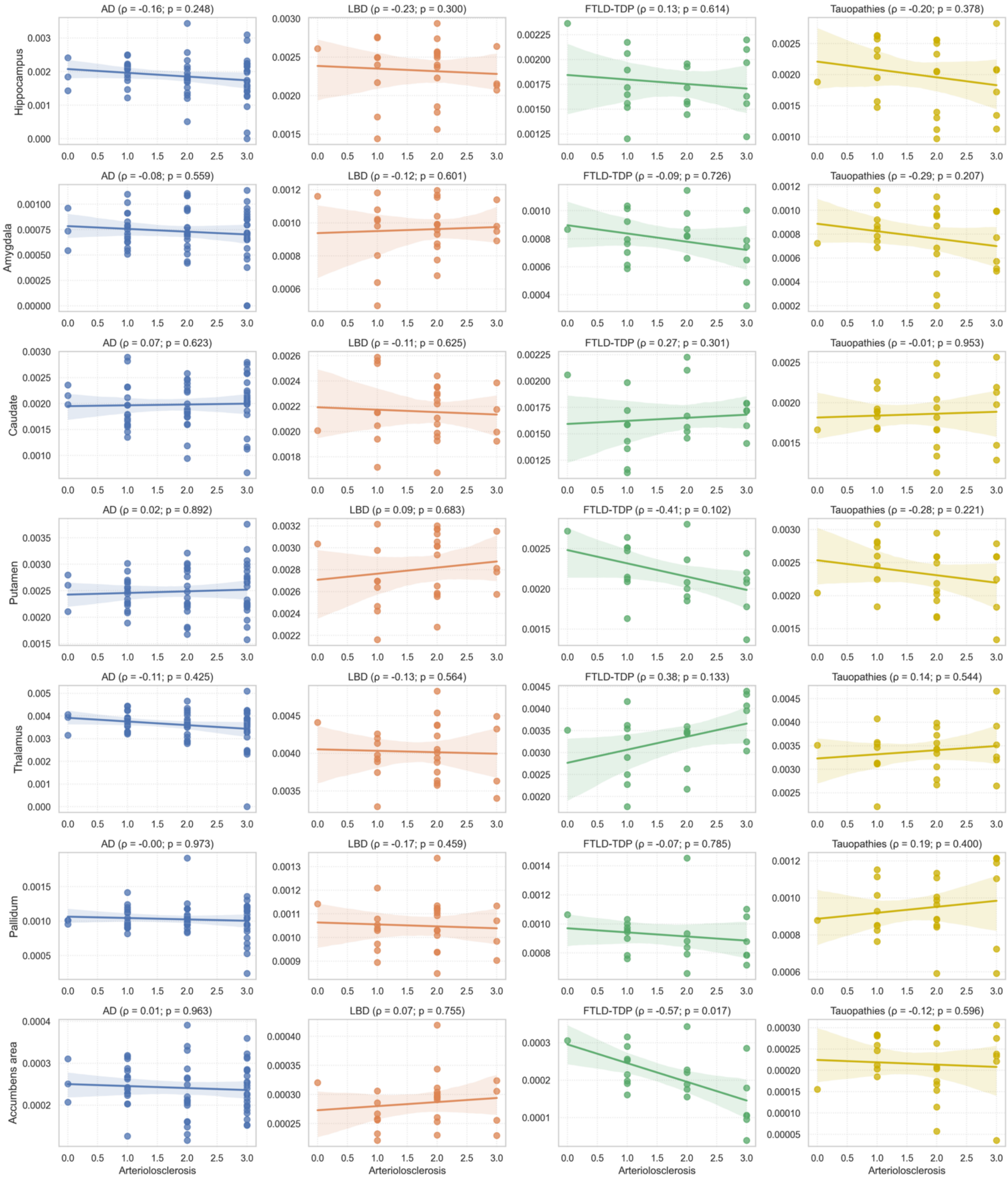

Partial Spearman correlations — Atherosclerosis

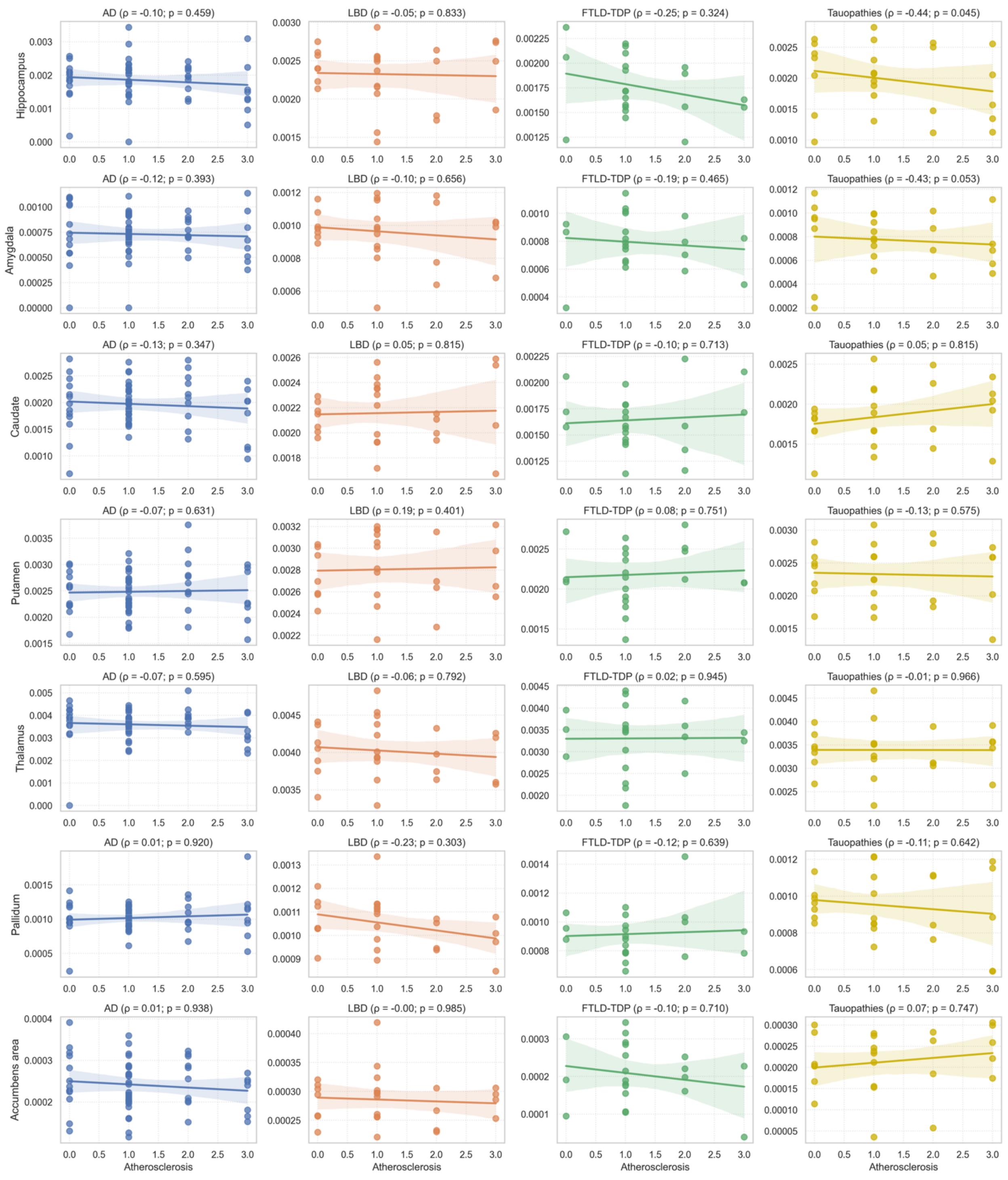

**(B)**

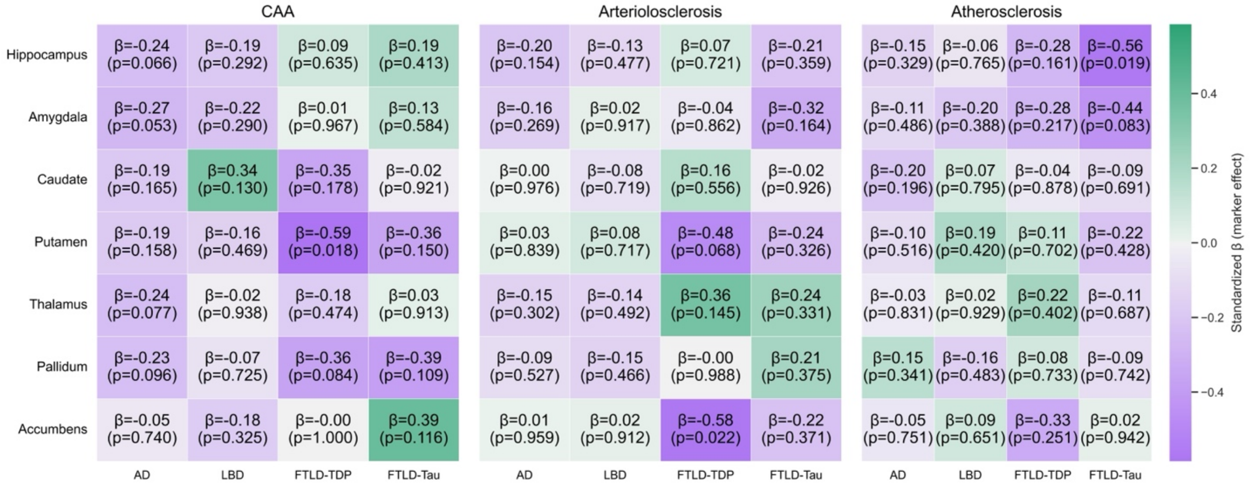


**Supplementary Figure S1.8. Partial Spearman correlations and ordinary least squares model between subcortical/limbic volumes and global vascular burden.** **(A)** Covariate-adjusted Spearman correlations (ρ) between ICV-normalized postmortem subcortical/limbic volumes and global semi-quantitative markers of cerebral amyloid angiopathy (CAA), and vascular pathology (arteriolosclerosis and atherosclerosis) across Alzheimer's disease, Lewy body disease, FTLD-TDP, and FTLD-Tau. Correlations are adjusted for age at death, sex, education, and PMI. Statistical significance is shown using FDR-corrected *p<0.05, **p<0.01, and ***p<0.001. **(B)** Heatmaps show standardized β coefficients from OLS models quantifying the association between subcortical/limbic postmortem MRI volumes and global semi-quantitative markers of cerebral amyloid angiopathy (CAA), and vascular pathology (arteriolosclerosis and atherosclerosis). Each cell reports the standardized effect size (β) and the corresponding uncorrected p value; boxes highlight uncorrected p<0.05, and asterisks denote significance after false discovery rate correction using the Benjamini–Hochberg procedure applied separately within each diagnostic group across all structures and pathology predictors (*p<0.05 (also outlined), **p<0.01, ***p<0.001). All models included age at death, sex, years of education, and postmortem interval as covariates.

**Supplementary Table S1.1. Postmortem subcortical/limbic volumes across neuropathological groups.** Pairwise differences between normalized (by ICV) groups (AD, LBD, FTLD-TDP and FTLD-Tau) were assessed using likelihood-ratio tests adjusting for age at death, sex, education, and postmortem interval. Covariate-adjusted adjusted mean differences (Δ), standard errors, and 95% confidence intervals are reported. FDR correction was applied globally across all pairwise tests. Scientific notation is used for all numeric values. Significant FDR-corrected p-values (p<0.05; p<0.01; *p<0.001).

##### Hippocampus

| Group comparison | Adjusted Means | Δ (Adj Diff) | SE | CI (95%) | LR | p (raw) | p (FDR) |
|---|---|---|---|---|---|---|---|
| LBD vs FTLD-TDP | LBD = 2.32<br>FTLD-TDP = 1.76 | 0.562 | 0.000 | [0.561, 0.562] | 32.6 | <0.0001 | <0.0001*** |
| AD vs LBD | AD = 1.87<br>LBD = 2.32 | -0.451 | 0.122 | [-0.691, -0.211] | 16.0 | <0.0001 | < 0.001*** |
| LBD vs FTLD-Tau | LBD = 2.32<br>FTLD-Tau = 1.98 | 0.348 | 0.143 | [0.066, 0.628] | 7.77 | 0.0053 | 0.010* |
| FTLD-TDP vs FTLD-Tau | FTLD-TDP = 1.76<br>FTLD-Tau = 1.98 | -0.215 | 0.135 | [-0.478, 0.049] | 2.98 | 0.084 | 0.127 |
| AD vs FTLD-TDP | AD = 1.87<br>FTLD-TDP = 1.76 | 0.111 | 0.136 | [-0.155, 0.377] | 1.12 | 0.289 | 0.349 |
| AD vs FTLD-Tau | AD = 1.87<br>FTLD-Tau = 1.98 | -0.103 | 0.139 | [-0.377, 0.170] | 0.458 | 0.499 | 0.537 |

##### Amygdala

| Group comparison | Adjusted Means | Δ (Adj Diff) | SE | CI (95%) | LR | p (raw) | p (FDR) |
|---|---|---|---|---|---|---|---|
| AD vs LBD | AD = 0.732<br>LBD = 0.959 | -0.228 | 0.053 | [-0.332, -0.124] | 17.500 | < 0.0001 | < 0.001*** |
| LBD vs FTLD-TDP | LBD = 0.959<br>FTLD-TDP = 0.790 | 0.169 | 0.000 | [0.168, 0.169] | 12.900 | < 0.001 | 0.001** |
| LBD vs FTLD-Tau | LBD = 0.959<br>FTLD-Tau = 0.772 | 0.187 | 0.065 | [0.058, 0.316] | 7.790 | 0.005 | 0.011* |
| AD vs FTLD-TDP | AD = 0.732<br>FTLD-TDP = 0.790 | -0.058 | 0.059 | [-0.176, 0.058] | 0.767 | 0.381 | 0.433 |
| AD vs FTLD-Tau | AD = 0.732<br>FTLD-Tau = 0.772 | -0.040 | 0.058 | [-0.155, 0.074] | 0.656 | 0.418 | 0.462 |
| FTLD-TDP vs FTLD-Tau | FTLD-TDP = 0.790<br>FTLD-Tau = 0.772 | 0.018 | 0.067 | [-0.114, 0.150] | 0.013 | 0.909 | 0.909 |

#### Nucleus Accumbens

| Group comparison | Adjusted Means | Δ (Adj Diff) | SE | CI (95%) | LR | p (raw) | p (FDR) |
|---|---|---|---|---|---|---|---|
| LBD vs FTLD-TDP | LBD = 0.285<br>FTLD-TDP = 0.205 | 0.080 | 0.00 | [0.080, 0.080] | 21.2 | < 0.0001 | < 0.0001*** |
| LBD vs FTLD-Tau | LBD = 0.285<br>FTLD-Tau = 0.214 | 0.071 | 0.017 | [0.036, 0.106] | 18.7 | < 0.0001 | < 0.0001*** |
| AD vs LBD | AD = 0.241<br>LBD = 0.285 | -0.045 | 0.013 | [-0.071, -0.018] | 13.4 | < 0.001 | < 0.001*** |
| AD vs FTLD-TDP | AD = 0.241<br>FTLD-TDP = 0.205 | 0.035 | 0.017 | [0.0014, 0.069] | 5.47 | 0.019 | 0.037* |
| AD vs FTLD-Tau | AD = 0.241<br>FTLD-Tau = 0.214 | 0.026 | 0.015 | [-0.0046, 0.057] | 3.22 | 0.073 | 0.113 |
| FTLD-TDP vs FTLD-Tau | FTLD-TDP = 0.205<br>FTLD-Tau = 0.214 | -0.009 | 0.024 | [-0.056, 0.038] | 0.205 | 0.651 | 0.683 |

#### Thalamus

| Group comparison | Adjusted Means | Δ (Adj Diff) | SE | CI (95%) | LR | p (raw) | p (FDR) |
|---|---|---|---|---|---|---|---|
| LBD vs FTLD-Tau | LBD = 4.02<br>FTLD-Tau = 3.39 | 0.627 | 0.134 | [0.363, 0.890] | 24.0 | < 0.0001 | < 0.0001*** |
| LBD vs FTLD-TDP | LBD = 4.02<br>FTLD-TDP = 3.31 | 0.712 | 0.00 | [0.711, 0.712] | 22.0 | < 0.0001 | < 0.0001*** |
| AD vs LBD | AD = 3.61<br>LBD = 4.02 | -0.412 | 0.154 | [-0.715, -0.110] | 81.9 | 0.004 | 0.009** |
| AD vs FTLD-TDP | AD = 3.61<br>FTLD-TDP = 3.31 | 0.300 | 0.189 | [-0.0706, 0.671] | 41.8 | 0.041 | 0.075 |
| AD vs FTLD-Tau | AD = 3.61<br>FTLD-Tau = 3.39 | 0.215 | 0.165 | [-0.108, 0.538] | 23.5 | 0.126 | 0.176 |
| FTLD-TDP vs FTLD-Tau | FTLD-TDP = 3.31<br>FTLD-Tau = 3.39 | -0.0855 | 0.192 | [-0.462, 0.291] | 0.127 | 0.722 | 0.739 |

#### Caudate

| Group comparison | Adjusted Means | Δ (Adj Diff) | SE | CI (95%) | LR | p (raw) | p (FDR) |
|---|---|---|---|---|---|---|---|
| LBD vs FTLD-TDP | LBD = 2.16<br>FTLD-TDP = 1.65 | 0.512 | 0.00 | [0.511, 0.512] | 29.1 | < 0.0001 | < 0.0001*** |
| AD vs FTLD-TDP | AD = 2.00<br>FTLD-TDP = 1.65 | 0.353 | 0.112 | [0.134, 0.573] | 10.4 | 0.00128 | 0.004** |
| LBD vs FTLD-Tau | LBD = 2.16<br>FTLD-Tau = 1.86 | 0.299 | 0.092 | [0.119, 0.478] | 10.3 | 0.00134 | 0.004** |
| FTLD-TDP vs FTLD-Tau | FTLD-TDP = 1.65<br>FTLD-Tau = 1.86 | -0.213 | 0.099 | [-0.408, -0.019] | 3.45 | 0.0633 | 0.111 |
| AD vs LBD | AD = 2.00<br>LBD = 2.16 | -0.158 | 0.099 | [-0.354, 0.037] | 2.56 | 0.110 | 0.159 |
| AD vs FTLD-Tau | AD = 2.00<br>FTLD-Tau = 1.86 | 0.140 | 0.106 | [-0.067, 0.347] | 1.91 | 0.167 | 0.213 |

#### Putamen

| Group comparison | Adjusted Means | Δ (Adj Diff) | SE | CI (95%) | LR | p (raw) | p (FDR) |
|---|---|---|---|---|---|---|---|
| LBD vs FTLD-TDP | LBD = 2.81<br>FTLD-TDP = 2.18 | 0.623 | 0.00 | [0.622, 0.623] | 32.2 | < 0.0001 | < 0.0001*** |
| LBD vs FTLD-Tau | LBD = 2.81<br>FTLD-Tau = 2.33 | 0.479 | 0.114 | [0.256, 0.703] | 19.2 | < 0.0001 | < 0.0001*** |
| AD vs LBD | AD = 2.49<br>LBD = 2.81 | -0.311 | 0.0965 | [-0.500, -0.122] | 10.4 | 0.00123 | 0.00353** |
| AD vs FTLD-TDP | AD = 2.49<br>FTLD-TDP = 2.18 | 0.312 | 0.109 | [0.099, 0.525] | 7.94 | 0.005 | 0.0106* |
| AD vs FTLD-Tau | AD = 2.49<br>FTLD-Tau = 2.33 | 0.168 | 0.105 | [-0.037, 0.373] | 3.30 | 0.069 | 0.112 |
| FTLD-TDP vs FTLD-Tau | FTLD-TDP = 2.18<br>Tau = 2.33 | -0.144 | 0.133 | [-0.404, 0.116] | 1.12 | 0.291 | 0.349 |

#### Pallidum

| Group comparison | Adjusted Means | Δ (Adj Diff) | SE | CI (95%) | LR | p (raw) | p (FDR) |
|---|---|---|---|---|---|---|---|
| LBD vs FTLD-TDP | LBD = 1.05<br>FTLD-TDP = 0.919 | 0.130 | 0.00 | [0.130, 0.131] | 14.5 | < 0.001 | < 0.001*** |
| LBD vs FTLD-Tau | LBD = 1.05<br>FTLD-Tau = 0.947 | 0.102 | 0.042 | [0.021, 0.018] | 8.80 | 0.003 | 0.007** |
| AD vs FTLD-TDP | AD = 1.01<br>FTLD-TDP = 0.919 | 0.0902 | 0.051 | [-0.009, 0.190] | 3.30 | 0.0695 | 0.112 |
| AD vs LBD | AD = 1.01<br>LBD = 1.05 | -0.040 | 0.043 | [-0.125, 0.044] | 1.97 | 0.160 | 0.211 |
| AD vs FTLD-Tau | AD = 1.01<br>FTLD-Tau = 0.947 | 0.062 | 0.046 | [-0.0287, 0. 153] | 1.97 | 0.161 | 0.211 |
| FTLD-TDP vs FTLD-Tau | FTLD-TDP = 0.919<br>FTLD-Tau = 0.947 | -0.028 | 0.054 | [-0.135, 0.079] | 0.767 | 0.381 | 0.433 |

**Supplementary Table S1.2. Primary pathology ordinary least squares model regression analysis between postmortem MRI volumes and primary neuropathological burden.** Regression models were fit to assess the relationship between ICV-normalized postmortem subcortical/limbic volumes and corresponding regional predominant molecular pathology measures within each diagnostic group: AD (p-tau), FTLD-Tau (p-tau), LBD (α-synuclein), and FTLD-TDP (TDP-43) adjusting for age at death, sex, postmortem interval, and years of education and corrected false discovery rate correction using the Benjamini–Hochberg procedure applied separately within each diagnostic group across all structures (*p<0.05, **p<0.01, ***p<0.001). Independent linear model was fit for each structure within each diagnostic group, including the region's predominant pathology measure together with covariates (age at death, sex, education, and postmortem interval). Standardized β coefficients, 95% CI, raw and FDR-corrected p-value (*p<0.05, **p<0.01, ***p<0.001). Pathology burden ratings were not available for the nucleus accumbens, and therefore this structure is not shown. Note that the columns: adjusted means, adjusted difference, standard error, confidence interval were multiplied by 1000 for an easier interpretation.

##### Alzheimer's disease

| Structure | Predictor | β | CI (95%) | p (raw) | p (FDR) |
|---|---|---|---|---|---|
| Amygdala | Age at death | -0.065 | [-0.336, 0.205] | 0.637 | 0.934 |
| | Sex | -0.042 | [-0.306, 0.223] | 0.759 | 0.934 |
| | Education | 0.002 | [-0.266, 0.270] | 0.989 | 0.989 |
| | PMI | 0.174 | [-0.090, 0.438] | 0.202 | 0.552 |
| | p-tau | -0.306 | [-0.578, -0.034] | 0.031 | 0.190 |
| Caudate | Age at death | -0.078 | [-0.340, 0.184] | 0.562 | 0.934 |
| | Sex | -0.032 | [-0.295, 0.230] | 0.810 | 0.934 |
| | Education | -0.060 | [-0.314, 0.194] | 0.644 | 0.934 |
| | PMI | 0.141 | [-0.117, 0.400] | 0.288 | 0.654 |
| | p-tau | -0.350 | [-0.614, -0.085] | 0.012 | 0.122 |
| Hippocampus | Age at death | -0.189 | [-0.429, 0.051] | 0.129 | 0.485 |
| | Sex | 0.050 | [-0.192, 0.293] | 0.685 | 0.934 |
| | Education | -0.021 | [-0.259, 0.217] | 0.862 | 0.934 |
| | PMI | 0.273 | [0.031, 0.515] | 0.031 | 0.190 |
| | p-tau | -0.388 | [-0.625, -0.150] | 0.002 | 0.034* |
| Pallidum | Age at death | -0.004 | [-0.273, 0.264] | 0.974 | 0.989 |
| | Sex | 0.143 | [-0.128, 0.415] | 0.305 | 0.654 |
| | Education | 0.043 | [-0.225, 0.310] | 0.756 | 0.934 |
| | PMI | 0.033 | [-0.236, 0.303] | 0.809 | 0.934 |
| | p-tau | 0.211 | [-0.053, 0.476] | 0.123 | 0.485 |
| Putamen | Age at death | 0.021 | [-0.223, 0.265] | 0.866 | 0.934 |
| | Sex | 0.028 | [-0.217, 0.273] | 0.823 | 0.934 |
| | Education | 0.131 | [-0.106, 0.368] | 0.284 | 0.654 |
| | PMI | 0.173 | [-0.069, 0.414] | 0.166 | 0.539 |
| | p-tau | -0.409 | [-0.656, -0.163] | 0.002 | 0.034* |
| Thalamus | Age at death | -0.219 | [-0.534, 0.097] | 0.179 | 0.539 |
| | Sex | -0.078 | [-0.347, 0.191] | 0.573 | 0.934 |
| | Education | 0.028 | [-0.237, 0.292] | 0.839 | 0.934 |
| | PMI | 0.220 | [-0.048, 0.487] | 0.114 | 0.485 |
| | p-tau | -0.025 | [-0.333, 0.283] | 0.872 | 0.934 |

#### Lewy body disease

| Structure | Predictor | β | CI (95%) | p (raw) | p (FDR) |
|---|---|---|---|---|---|
| Amygdala | Age at death | -0.171 | [-0.633, 0.291] | 0.476 | 0.826 |
| | Sex | -0.055 | [-0.511, 0.401] | 0.815 | 0.914 |
| | Education | -0.173 | [-0.640, 0.295] | 0.477 | 0.826 |
| | PMI | 0.407 | [-0.050, 0.865] | 0.095 | 0.479 |
| | α-synuclein | -0.170 | [-0.649, 0.310] | 0.495 | 0.826 |
| Caudate | Age at death | -0.089 | [-0.531, 0.353] | 0.696 | 0.914 |
| | Sex | -0.051 | [-0.478, 0.377] | 0.819 | 0.914 |
| | Education | -0.016 | [-0.485, 0.452] | 0.945 | 0.950 |
| | PMI | 0.059 | [-0.408, 0.527] | 0.806 | 0.914 |
| | α-synuclein | -0.376 | [-0.786, 0.034] | 0.087 | 0.479 |
| Hippocampus | Age at death | -0.287 | [-0.668, 0.095] | 0.156 | 0.521 |
| | Sex | 0.204 | [-0.163, 0.570] | 0.288 | 0.797 |
| | Education | -0.077 | [-0.478, 0.323] | 0.708 | 0.914 |
| | PMI | 0.502 | [0.088, 0.916] | 0.027 | 0.415 |
| | α-synuclein | -0.029 | [-0.405, 0.348] | 0.882 | 0.945 |
| Pallidum | Age at death | -0.460 | [-0.883, -0.036] | 0.046 | 0.460 |
| | Sex | 0.124 | [-0.285, 0.533] | 0.560 | 0.884 |
| | Education | 0.060 | [-0.387, 0.507] | 0.795 | 0.914 |
| | PMI | -0.235 | [-0.678, 0.209] | 0.312 | 0.797 |
| | α-synuclein | -0.179 | [-0.569, 0.211] | 0.379 | 0.797 |
| Putamen | Age at death | -0.372 | [-0.811, 0.066] | 0.111 | 0.479 |
| | Sex | 0.206 | [-0.218, 0.630] | 0.352 | 0.797 |
| | Education | -0.054 | [-0.518, 0.411] | 0.823 | 0.914 |
| | PMI | -0.015 | [-0.479, 0.449] | 0.950 | 0.950 |
| | α-synuclein | -0.184 | [-0.591, 0.223] | 0.386 | 0.797 |
| Thalamus | Age at death | -0.326 | [-0.659, 0.007] | 0.070 | 0.479 |
| | Sex | -0.139 | [-0.455, 0.177] | 0.398 | 0.797 |
| | Education | -0.093 | [-0.444, 0.258] | 0.608 | 0.913 |
| | PMI | 0.262 | [-0.079, 0.604] | 0.148 | 0.521 |
| | α-synuclein | -0.551 | [-0.856, -0.247] | 0.002 | 0.064 |

#### FTLD-TDP

| Structure | Predictor | β | CI (95%) | p (raw) | p (FDR) |
|---|---|---|---|---|---|
| Amygdala | Age at death | -0.543 | [-0.971, -0.115] | 0.025 | 0.267 |
| | Sex | -0.096 | [-0.540, 0.348] | 0.677 | 0.923 |
| | Education | 0.321 | [-0.234, 0.877] | 0.274 | 0.588 |
| | PMI | -0.578 | [-1.118, -0.039] | 0.053 | 0.370 |
| | TDP-43 | 0.128 | [-0.319, 0.574] | 0.583 | 0.886 |
| Caudate | Age at death | -0.387 | [-0.879, 0.105] | 0.145 | 0.397 |
| | Sex | -0.098 | [-0.516, 0.321] | 0.655 | 0.923 |
| | Education | -0.037 | [-0.491, 0.417] | 0.875 | 0.976 |
| | PMI | -0.751 | [-1.243, -0.260] | 0.009 | 0.267 |
| | TDP-43 | -0.415 | [-0.912, 0.082] | 0.124 | 0.397 |
| Hippocampus | Age at death | -0.514 | [-1.091, 0.063] | 0.102 | 0.397 |
| | Sex | 0.148 | [-0.325, 0.621] | 0.550 | 0.886 |
| | Education | -0.283 | [-0.698, 0.132] | 0.202 | 0.507 |
| | PMI | -0.070 | [-0.563, 0.423] | 0.784 | 0.941 |
| | TDP-43 | 0.020 | [-0.520, 0.560] | 0.944 | 0.976 |
| Pallidum | Age at death | -0.485 | [-1.040, 0.070] | 0.110 | 0.397 |
| | Sex | -0.361 | [-0.812, 0.091] | 0.141 | 0.397 |
| | Education | 0.455 | [-0.000, 0.910] | 0.072 | 0.370 |
| | PMI | -0.084 | [-0.593, 0.424] | 0.750 | 0.938 |
| | TDP-43 | 0.024 | [-0.499, 0.547] | 0.930 | 0.976 |
| Putamen | Age at death | -0.303 | [-0.820, 0.214] | 0.270 | 0.588 |
| | Sex | -0.128 | [-0.568, 0.311] | 0.576 | 0.886 |
| | Education | -0.233 | [-0.709, 0.244] | 0.354 | 0.664 |
| | PMI | -0.002 | [-0.518, 0.514] | 0.995 | 0.995 |
| | TDP-43 | -0.659 | [-1.181, -0.137] | 0.026 | 0.267 |
| Thalamus | Age at death | -0.152 | [-0.696, 0.391] | 0.591 | 0.886 |
| | Sex | 0.033 | [-0.437, 0.504] | 0.891 | 0.976 |
| | Education | -0.499 | [-1.006, 0.008] | 0.074 | 0.370 |
| | PMI | -0.105 | [-0.646, 0.436] | 0.709 | 0.925 |
| | TDP-43 | -0.279 | [-0.795, 0.237] | 0.307 | 0.614 |

#### FTLD-Tau

| Structure | Predictor | β | CI (95%) | p (raw) | p (FDR) |
|---|---|---|---|---|---|
| Amygdala | Age at death | 0.161 | [-0.191, 0.512] | 0.382 | 0.721 |
| | Sex | 0.073 | [-0.319, 0.466] | 0.718 | 0.847 |
| | Education | -0.208 | [-0.588, 0.173] | 0.299 | 0.690 |
| | PMI | -0.438 | [-0.795, -0.080] | 0.027 | 0.410 |
| | p-tau | -0.469 | [-0.805, -0.133] | 0.013 | 0.406 |
| Caudate | Age at death | 0.320 | [-0.062, 0.703] | 0.117 | 0.502 |
| | Sex | 0.421 | [-0.027, 0.868] | 0.081 | 0.450 |
| | Education | 0.185 | [-0.244, 0.614] | 0.409 | 0.721 |
| | PMI | -0.110 | [-0.480, 0.260] | 0.566 | 0.809 |
| | p-tau | 0.256 | [-0.174, 0.685] | 0.257 | 0.676 |
| Hippocampus | Age at death | 0.055 | [-0.318, 0.428] | 0.774 | 0.847 |
| | Sex | 0.089 | [-0.353, 0.530] | 0.698 | 0.847 |
| | Education | -0.089 | [-0.515, 0.337] | 0.687 | 0.847 |
| | PMI | -0.343 | [-0.720, 0.034] | 0.090 | 0.450 |
| | p-tau | -0.413 | [-0.784, -0.043] | 0.041 | 0.414 |
| Pallidum | Age at death | -0.266 | [-0.688, 0.156] | 0.231 | 0.676 |
| | Sex | 0.164 | [-0.344, 0.672] | 0.534 | 0.805 |
| | Education | -0.280 | [-0.762, 0.203] | 0.270 | 0.676 |
| | PMI | -0.339 | [-0.826, 0.148] | 0.188 | 0.676 |
| | p-tau | -0.260 | [-0.783, 0.263] | 0.341 | 0.721 |
| Putamen | Age at death | 0.058 | [-0.406, 0.521] | 0.810 | 0.847 |
| | Sex | 0.351 | [-0.191, 0.893] | 0.220 | 0.676 |
| | Education | 0.086 | [-0.435, 0.606] | 0.750 | 0.847 |
| | PMI | -0.154 | [-0.602, 0.294] | 0.508 | 0.805 |
| | p-tau | -0.062 | [-0.582, 0.459] | 0.818 | 0.847 |
| Thalamus | Age at death | 0.020 | [-0.410, 0.451] | 0.927 | 0.927 |
| | Sex | 0.498 | [-0.005, 1.001] | 0.067 | 0.450 |
| | Education | 0.154 | [-0.325, 0.632] | 0.536 | 0.805 |
| | PMI | -0.082 | [-0.525, 0.360] | 0.719 | 0.847 |
| | p-tau | -0.199 | [-0.639, 0.241] | 0.387 | 0.721 |

**Supplementary Table S1.3. Polypathology ordinary least squares models for structure-pathology relationships.** Shown are the standardized β coefficients from OLS models quantifying the association between regional postmortem MRI volumes and multiple semi-quantitative pathology ratings across the four diagnostic groups (AD, LBD, FTLD-TDP, FTLD-tau). Independent linear model was fit for each structure within each diagnostic group with multiple concomitant pathologies measures together with covariates (age at death, sex, education, and postmortem interval). Standardized β coefficients, standard error, 95% CI, raw and FDR-corrected p-value (*p<0.05, **p<0.01, ***p<0.001) using the Benjamini–Hochberg procedure. Pathology burden ratings were not available for the nucleus accumbens, and therefore this structure is not shown.

##### Alzheimer's disease

| Structure | Predictor | β | SE | CI (95%) | p(raw) | p(FDR) |
|---|---|---|---|---|---|---|
| Amygdala | Age at death | 0.021 | 0.135 | [-0.245, 0.286] | 0.879 | 0.974 |
| | Education | -0.053 | 0.129 | [-0.306, 0.200] | 0.683 | 0.974 |
| | PMI | 0.070 | 0.130 | [-0.186, 0.325] | 0.594 | 0.974 |
| | Sex | -0.005 | 0.127 | [-0.254, 0.244] | 0.968 | 0.991 |
| | TDP-43 | -0.349 | 0.131 | [-0.605, -0.093] | 0.010 | 0.141 |
| | p-tau | -0.193 | 0.138 | [-0.463, 0.078] | 0.168 | 0.636 |
| | α-synuclein | -0.160 | 0.135 | [-0.425, 0.105] | 0.241 | 0.676 |
| Caudate | Age at death | -0.044 | 0.141 | [-0.320, 0.232] | 0.757 | 0.974 |
| | Education | -0.081 | 0.133 | [-0.342, 0.180] | 0.546 | 0.955 |
| | PMI | 0.141 | 0.135 | [-0.124, 0.406] | 0.301 | 0.744 |
| | Sex | 0.015 | 0.138 | [-0.254, 0.285] | 0.912 | 0.980 |
| | TDP-43 | -0.261 | 0.135 | [-0.525, 0.003] | 0.057 | 0.486 |
| | p-tau | -0.249 | 0.140 | [-0.523, 0.025] | 0.081 | 0.538 |
| | α-synuclein | 0.141 | 0.134 | [-0.122, 0.404] | 0.300 | 0.744 |
| Hippocampus | Age at death | -0.172 | 0.127 | [-0.421, 0.077] | 0.182 | 0.636 |
| | Education | -0.024 | 0.126 | [-0.271, 0.223] | 0.847 | 0.974 |
| | PMI | 0.261 | 0.125 | [0.016, 0.507] | 0.041 | 0.440 |
| | Sex | 0.044 | 0.126 | [-0.203, 0.291] | 0.729 | 0.974 |
| | TDP-43 | -0.061 | 0.131 | [-0.317, 0.195] | 0.643 | 0.974 |
| | p-tau | -0.355 | 0.127 | [-0.603, -0.106] | 0.007 | 0.141 |
| | α-synuclein | -0.092 | 0.128 | [-0.344, 0.159] | 0.476 | 0.869 |
| Pallidum | Age at death | 0.000 | 0.137 | [-0.269, 0.270] | 0.999 | 0.999 |
| | Education | 0.052 | 0.138 | [-0.218, 0.322] | 0.707 | 0.974 |
| | PMI | 0.043 | 0.139 | [-0.229, 0.314] | 0.758 | 0.974 |
| | Sex | 0.126 | 0.141 | [-0.150, 0.403] | 0.375 | 0.788 |
| | TDP-43 | -0.000 | 0.000 | [-0.000, 0.000] | 0.102 | 0.538 |
| | p-tau | 0.231 | 0.138 | [-0.040, 0.502] | 0.100 | 0.538 |
| | α-synuclein | -0.105 | 0.139 | [-0.377, 0.167] | 0.453 | 0.869 |
| Putamen | Age at death | 0.020 | 0.132 | [-0.238, 0.278] | 0.881 | 0.974 |
| | Education | 0.113 | 0.125 | [-0.131, 0.357] | 0.369 | 0.788 |
| | PMI | 0.178 | 0.126 | [-0.069, 0.426] | 0.164 | 0.636 |
| | Sex | 0.056 | 0.129 | [-0.196, 0.308] | 0.664 | 0.974 |
| | TDP-43 | -0.155 | 0.126 | [-0.402, 0.092] | 0.224 | 0.673 |
| | p-tau | -0.392 | 0.131 | [-0.648, -0.136] | 0.004 | 0.141 |
| | α-synuclein | 0.026 | 0.126 | [-0.220, 0.272] | 0.838 | 0.974 |
| Thalamus | Age at death | -0.162 | 0.175 | [-0.504, 0.180] | 0.358 | 0.788 |
| | Education | 0.012 | 0.138 | [-0.259, 0.282] | 0.933 | 0.980 |
| | PMI | 0.206 | 0.138 | [-0.063, 0.476] | 0.139 | 0.636 |
| | Sex | -0.101 | 0.139 | [-0.373, 0.171] | 0.472 | 0.869 |
| | TDP-43 | -0.179 | 0.142 | [-0.457, 0.099] | 0.212 | 0.673 |
| | p-tau | -0.035 | 0.162 | [-0.352, 0.282] | 0.828 | 0.974 |
| | α-synuclein | 0.041 | 0.139 | [-0.231, 0.313] | 0.769 | 0.974 |

#### Lewy body disease

| Structure | Predictor | β | SE | CI (95%) | p(raw) | p(FDR) |
|---|---|---|---|---|---|---|
| Amygdala | Age at death | 0.031 | 0.263 | [-0.484, 0.546] | 0.907 | 0.967 |
| | Education | 0.011 | 0.271 | [-0.519, 0.542] | 0.967 | 0.967 |
| | PMI | 0.229 | 0.262 | [-0.284, 0.742] | 0.394 | 0.752 |
| | Sex | -0.047 | 0.236 | [-0.510, 0.416] | 0.845 | 0.967 |
| | TDP-43 | -0.265 | 0.223 | [-0.701, 0.171] | 0.250 | 0.597 |
| | p-tau | -0.351 | 0.268 | [-0.876, 0.173] | 0.207 | 0.597 |
| | α-synuclein | 0.049 | 0.284 | [-0.508, 0.605] | 0.866 | 0.967 |
| Caudate | Age at death | -0.077 | 0.224 | [-0.517, 0.362] | 0.734 | 0.967 |
| | Education | -0.052 | 0.239 | [-0.520, 0.417] | 0.831 | 0.967 |
| | PMI | 0.062 | 0.237 | [-0.402, 0.527] | 0.795 | 0.967 |
| | Sex | -0.089 | 0.219 | [-0.518, 0.341] | 0.690 | 0.966 |
| | TDP-43 | -0.000 | 0.000 | [-0.000, 0.000] | 0.120 | 0.559 |
| | p-tau | 0.286 | 0.252 | [-0.208, 0.780] | 0.270 | 0.597 |
| | α-synuclein | -0.541 | 0.253 | [-1.038, -0.044] | 0.046 | 0.559 |
| Hippocampus | Age at death | -0.199 | 0.190 | [-0.571, 0.174] | 0.310 | 0.651 |
| | Education | 0.086 | 0.207 | [-0.319, 0.491] | 0.682 | 0.966 |
| | PMI | 0.280 | 0.228 | [-0.167, 0.727] | 0.235 | 0.597 |
| | Sex | 0.234 | 0.192 | [-0.144, 0.611] | 0.241 | 0.597 |
| | TDP-43 | -0.326 | 0.188 | [-0.695, 0.043] | 0.100 | 0.559 |
| | p-tau | -0.312 | 0.238 | [-0.779, 0.156] | 0.208 | 0.597 |
| | α-synuclein | 0.019 | 0.220 | [-0.412, 0.450] | 0.932 | 0.967 |
| Pallidum | Age at death | -0.473 | 0.233 | [-0.929, -0.017] | 0.056 | 0.559 |
| | Education | 0.069 | 0.239 | [-0.399, 0.538] | 0.775 | 0.967 |
| | PMI | -0.222 | 0.241 | [-0.696, 0.251] | 0.369 | 0.738 |
| | Sex | 0.123 | 0.214 | [-0.296, 0.543] | 0.571 | 0.960 |
| | TDP-43 | -0.000 | 0.000 | [-0.000, 0.000] | 0.257 | 0.597 |
| | p-tau | -0.045 | 0.242 | [-0.518, 0.428] | 0.854 | 0.967 |
| | α-synuclein | -0.169 | 0.211 | [-0.582, 0.244] | 0.433 | 0.791 |
| Putamen | Age at death | -0.358 | 0.219 | [-0.787, 0.072] | 0.119 | 0.559 |
| | Education | -0.095 | 0.234 | [-0.554, 0.363] | 0.688 | 0.966 |
| | PMI | -0.011 | 0.232 | [-0.465, 0.442] | 0.962 | 0.967 |
| | Sex | 0.161 | 0.214 | [-0.259, 0.581] | 0.462 | 0.808 |
| | TDP-43 | -0.000 | 0.000 | [-0.000, 0.000] | 0.919 | 0.559 |
| | p-tau | 0.340 | 0.246 | [-0.142, 0.823] | 0.183 | 0.597 |
| | α-synuclein | -0.380 | 0.248 | [-0.865, 0.106] | 0.142 | 0.595 |
| Thalamus | Age at death | -0.346 | 0.169 | [-0.676, -0.015] | 0.055 | 0.559 |
| | Education | -0.084 | 0.177 | [-0.431, 0.263] | 0.641 | 0.966 |
| | PMI | 0.305 | 0.176 | [-0.040, 0.650] | 0.100 | 0.559 |
| | Sex | -0.073 | 0.168 | [-0.403, 0.257] | 0.671 | 0.966 |
| | TDP-43 | 0.000 | 0.000 | [-0.000, 0.000] | 0.912 | 0.967 |
| | p-tau | -0.204 | 0.170 | [-0.537, 0.128] | 0.244 | 0.597 |
| | α-synuclein | -0.516 | 0.156 | [-0.823, -0.210] | 0.003 | 0.168 |

#### FTLD-TDP

| Structure | Predictor | β | SE | CI (95%) | p(raw) | p(FDR) |
|---|---|---|---|---|---|---|
| Amygdala | Age at death | -0.453 | 0.218 | [-0.879, -0.026] | 0.057 | 0.486 |
| | Education | 0.176 | 0.303 | [-0.419, 0.771] | 0.572 | 0.874 |
| | PMI | -0.556 | 0.273 | [-1.091, -0.021] | 0.062 | 0.486 |
| | Sex | -0.034 | 0.263 | [-0.549, 0.482] | 0.900 | 0.945 |
| | TDP-43 | 0.112 | 0.225 | [-0.329, 0.552] | 0.628 | 0.910 |
| | p-tau | 0.207 | 0.225 | [-0.234, 0.649] | 0.374 | 0.786 |
| | α-synuclein | -0.303 | 0.214 | [-0.724, 0.117] | 0.181 | 0.662 |
| Caudate | Age at death | -0.332 | 0.276 | [-0.874, 0.209] | 0.252 | 0.662 |
| | Education | 0.044 | 0.268 | [-0.481, 0.568] | 0.873 | 0.945 |
| | PMI | -0.737 | 0.268 | [-1.263, -0.212] | 0.017 | 0.486 |
| | Sex | -0.065 | 0.238 | [-0.532, 0.403] | 0.791 | 0.945 |
| | TDP-43 | -0.382 | 0.272 | [-0.915, 0.151] | 0.186 | 0.662 |
| | p-tau | 0.143 | 0.242 | [-0.331, 0.617] | 0.565 | 0.874 |
| | α-synuclein | -0.138 | 0.237 | [-0.603, 0.327] | 0.572 | 0.874 |
| Hippocampus | Age at death | -0.203 | 0.291 | [-0.773, 0.367] | 0.499 | 0.873 |
| | Education | -0.319 | 0.190 | [-0.691, 0.053] | 0.119 | 0.557 |
| | PMI | 0.069 | 0.230 | [-0.381, 0.519] | 0.769 | 0.945 |
| | Sex | 0.536 | 0.271 | [0.005, 1.068] | 0.071 | 0.486 |
| | TDP-43 | 0.348 | 0.279 | [-0.199, 0.895] | 0.237 | 0.662 |
| | p-tau | 0.271 | 0.209 | [-0.138, 0.680] | 0.218 | 0.662 |
| | α-synuclein | 0.384 | 0.202 | [-0.011, 0.780] | 0.081 | 0.486 |
| Pallidum | Age at death | -0.363 | 0.363 | [-1.073, 0.348] | 0.337 | 0.748 |
| | Education | 0.533 | 0.276 | [-0.008, 1.074] | 0.077 | 0.486 |
| | PMI | -0.071 | 0.268 | [-0.595, 0.454] | 0.796 | 0.945 |
| | Sex | -0.282 | 0.275 | [-0.820, 0.256] | 0.325 | 0.748 |
| | TDP-43 | 0.068 | 0.285 | [-0.491, 0.626] | 0.817 | 0.945 |
| | p-tau | 0.184 | 0.325 | [-0.454, 0.821] | 0.583 | 0.874 |
| | α-synuclein | -0.000 | 0.000 | [-0.000, 0.000] | 0.464 | 0.848 |
| Putamen | Age at death | -0.358 | 0.285 | [-0.916, 0.201] | 0.233 | 0.662 |
| | Education | -0.335 | 0.276 | [-0.875, 0.206] | 0.248 | 0.662 |
| | PMI | -0.000 | 0.276 | [-0.542, 0.541] | 0.999 | 0.999 |
| | Sex | -0.200 | 0.246 | [-0.682, 0.282] | 0.431 | 0.823 |
| | TDP-43 | -0.692 | 0.280 | [-1.242, -0.143] | 0.029 | 0.486 |
| | p-tau | -0.249 | 0.249 | [-0.738, 0.240] | 0.338 | 0.748 |
| | α-synuclein | 0.093 | 0.245 | [-0.387, 0.573] | 0.710 | 0.935 |
| Thalamus | Age at death | -0.140 | 0.331 | [-0.788, 0.508] | 0.679 | 0.935 |
| | Education | -0.495 | 0.295 | [-1.073, 0.083] | 0.119 | 0.557 |
| | PMI | -0.114 | 0.303 | [-0.707, 0.479] | 0.712 | 0.935 |
| | Sex | 0.036 | 0.277 | [-0.508, 0.580] | 0.899 | 0.945 |
| | TDP-43 | -0.281 | 0.335 | [-0.938, 0.375] | 0.417 | 0.823 |
| | p-tau | 0.047 | 0.283 | [-0.507, 0.601] | 0.871 | 0.945 |
| | α-synuclein | -0.010 | 0.317 | [-0.631, 0.611] | 0.975 | 0.998 |

#### FTLD-Tau

| Structure | Predictor | β | SE | CI (95%) | p(raw) | p(FDR) |
|---|---|---|---|---|---|---|
| Amygdala | Age at death | 0.164 | 0.192 | [-0.213, 0.541] | 0.408 | 0.634 |
| | Education | -0.393 | 0.271 | [-0.924, 0.139] | 0.167 | 0.390 |
| | PMI | -0.490 | 0.194 | [-0.870, -0.111] | 0.022 | 0.233 |
| | Sex | -0.010 | 0.221 | [-0.443, 0.423] | 0.965 | 0.988 |
| | TDP-43 | -0.222 | 0.239 | [-0.691, 0.247] | 0.368 | 0.618 |
| | p-tau | -0.504 | 0.181 | [-0.860, -0.149] | 0.013 | 0.233 |
| | α-synuclein | 0.084 | 0.184 | [-0.276, 0.445] | 0.654 | 0.855 |
| Caudate | Age at death | 0.264 | 0.176 | [-0.081, 0.608] | 0.152 | 0.390 |
| | Education | 0.582 | 0.245 | [0.102, 1.062] | 0.029 | 0.243 |
| | PMI | -0.102 | 0.175 | [-0.445, 0.241] | 0.567 | 0.779 |
| | Sex | 0.734 | 0.235 | [0.273, 1.195] | 0.006 | 0.233 |
| | TDP-43 | 0.499 | 0.197 | [0.112, 0.886] | 0.021 | 0.233 |
| | p-tau | 0.315 | 0.197 | [-0.071, 0.700] | 0.128 | 0.390 |
| | α-synuclein | -0.218 | 0.178 | [-0.566, 0.130] | 0.236 | 0.471 |
| Hippocampus | Age at death | 0.072 | 0.206 | [-0.332, 0.476] | 0.731 | 0.877 |
| | Education | -0.044 | 0.309 | [-0.649, 0.561] | 0.888 | 0.982 |
| | PMI | -0.357 | 0.211 | [-0.770, 0.056] | 0.108 | 0.379 |
| | Sex | 0.114 | 0.265 | [-0.405, 0.633] | 0.672 | 0.855 |
| | TDP-43 | 0.019 | 0.250 | [-0.470, 0.508] | 0.940 | 0.988 |
| | p-tau | -0.423 | 0.200 | [-0.816, -0.030] | 0.050 | 0.298 |
| | α-synuclein | -0.086 | 0.218 | [-0.514, 0.342] | 0.699 | 0.863 |
| Pallidum | Age at death | -0.315 | 0.206 | [-0.720, 0.089] | 0.145 | 0.390 |
| | Education | 0.073 | 0.285 | [-0.487, 0.632] | 0.802 | 0.911 |
| | PMI | -0.382 | 0.244 | [-0.860, 0.096] | 0.136 | 0.390 |
| | Sex | 0.518 | 0.296 | [-0.062, 1.098] | 0.098 | 0.375 |
| | TDP-43 | 0.475 | 0.240 | [0.004, 0.947] | 0.064 | 0.302 |
| | p-tau | -0.352 | 0.255 | [-0.851, 0.147] | 0.185 | 0.390 |
| | α-synuclein | -0.239 | 0.218 | [-0.666, 0.188] | 0.288 | 0.526 |
| Putamen | Age at death | 0.012 | 0.231 | [-0.441, 0.465] | 0.959 | 0.988 |
| | Education | 0.451 | 0.322 | [-0.180, 1.082] | 0.179 | 0.390 |
| | PMI | -0.131 | 0.230 | [-0.582, 0.319] | 0.575 | 0.779 |
| | Sex | 0.631 | 0.309 | [0.026, 1.237] | 0.056 | 0.298 |
| | TDP-43 | 0.477 | 0.259 | [-0.031, 0.985] | 0.083 | 0.351 |
| | p-tau | -0.002 | 0.258 | [-0.509, 0.505] | 0.994 | 0.994 |
| | α-synuclein | -0.152 | 0.233 | [-0.609, 0.305] | 0.524 | 0.779 |
| Thalamus | Age at death | -0.060 | 0.223 | [-0.497, 0.377] | 0.792 | 0.911 |
| | Education | 0.351 | 0.308 | [-0.254, 0.955] | 0.272 | 0.519 |
| | PMI | -0.145 | 0.236 | [-0.606, 0.317] | 0.548 | 0.779 |
| | Sex | 0.713 | 0.309 | [0.108, 1.319] | 0.034 | 0.243 |
| | TDP-43 | 0.225 | 0.263 | [-0.290, 0.740] | 0.404 | 0.634 |
| | p-tau | -0.232 | 0.230 | [-0.682, 0.218] | 0.327 | 0.572 |
| | α-synuclein | -0.327 | 0.237 | [-0.791, 0.136] | 0.186 | 0.390 |

**Supplementary Table S1.4. Ordinary least squares (OLS) models for structure/histology relationships.** Shown are the standardized β coefficients from OLS models quantifying the association between volumes and regional gliosis and neuronal loss semi-quantitative pathology ratings across the two diagnostic groups (AD and FTLD-Tau). Independent linear model was fit for each structure within each diagnostic group with multiple concomitant pathologies measures together with covariates (age at death, sex, education, and postmortem interval). Standardized β coefficients, standard error, 95% CI, raw and FDR-corrected p-value (*p<0.05, **p<0.01, ***p<0.001) using the Benjamini–Hochberg procedure. Pathology burden ratings were not available for the nucleus accumbens, and therefore this structure is not shown. Tables shown for the groups which survived corrections for multiple comparisons.

**Alzheimer's disease: Gliosis**

| Structure | Predictor | β | SE | CI (95%) | p(raw) | p(FDR) |
|---|---|---|---|---|---|---|
| Amygdala | Age at death | 0.022 | 0.132 | [-0.236, 0.280] | 0.867 | 0.936 |
| | Education | -0.136 | 0.135 | [-0.400, 0.128] | 0.318 | 0.850 |
| | PMI | 0.134 | 0.134 | [-0.128, 0.397] | 0.319 | 0.850 |
| | Sex | 0.031 | 0.138 | [-0.240, 0.301] | 0.824 | 0.936 |
| | Gliosis | -0.313 | 0.136 | [-0.579, -0.046] | 0.025 | 0.151 |
| Caudate | Age at death | 0.041 | 0.135 | [-0.224, 0.306] | 0.760 | 0.936 |
| | Education | -0.044 | 0.134 | [-0.307, 0.219] | 0.743 | 0.936 |
| | PMI | 0.143 | 0.138 | [-0.127, 0.412] | 0.304 | 0.850 |
| | Sex | 0.003 | 0.137 | [-0.266, 0.273] | 0.981 | 0.981 |
| | Gliosis | -0.219 | 0.134 | [-0.482, 0.044] | 0.108 | 0.168 |
| Hippocampus | Age at death | -0.082 | 0.132 | [-0.340, 0.176] | 0.535 | 0.900 |
| | Education | -0.097 | 0.132 | [-0.357, 0.162] | 0.465 | 0.900 |
| | PMI | 0.292 | 0.131 | [0.035, 0.550] | 0.030 | 0.724 |
| | Sex | 0.083 | 0.132 | [-0.176, 0.342] | 0.533 | 0.900 |
| | Gliosis | -0.214 | 0.133 | [-0.475, 0.046] | 0.112 | 0.168 |
| Pallidum | Age at death | -0.022 | 0.142 | [-0.299, 0.256] | 0.878 | 0.936 |
| | Education | 0.074 | 0.139 | [-0.200, 0.347] | 0.600 | 0.900 |
| | PMI | 0.018 | 0.142 | [-0.260, 0.297] | 0.897 | 0.936 |
| | Sex | 0.129 | 0.141 | [-0.148, 0.406] | 0.365 | 0.877 |
| | Gliosis | -0.060 | 0.139 | [-0.333, 0.214] | 0.669 | 0.789 |
| Putamen | Age at death | 0.155 | 0.130 | [-0.100, 0.409] | 0.238 | 0.850 |
| | Education | 0.148 | 0.129 | [-0.105, 0.401] | 0.255 | 0.850 |
| | PMI | 0.181 | 0.132 | [-0.078, 0.440] | 0.177 | 0.850 |
| | Sex | 0.074 | 0.132 | [-0.185, 0.333] | 0.579 | 0.900 |
| | Gliosis | -0.216 | 0.129 | [-0.468, 0.037] | 0.099 | 0.168 |
| Thalamus | Age at death | -0.207 | 0.135 | [-0.471, 0.057] | 0.130 | 0.850 |
| | Education | 0.025 | 0.135 | [-0.239, 0.290] | 0.852 | 0.936 |
| | PMI | 0.223 | 0.137 | [-0.047, 0.492] | 0.111 | 0.850 |
| | Sex | -0.081 | 0.138 | [-0.352, 0.189] | 0.557 | 0.900 |
| | Gliosis | 0.036 | 0.133 | [-0.224, 0.295] | 0.789 | 0.789 |

#### Alzheimer's disease: Neuronal loss

| Structure | Predictor | β | SE | CI (95%) | p(raw) | p(FDR) |
|---|---|---|---|---|---|---|
| Amygdala | Age at death | 0.059 | 0.128 | [-0.191, 0.310] | 0.644 | 0.909 |
| | Education | -0.127 | 0.129 | [-0.380, 0.126] | 0.329 | 0.808 |
| | PMI | 0.125 | 0.129 | [-0.128, 0.379] | 0.336 | 0.808 |
| | Sex | 0.018 | 0.132 | [-0.239, 0.276] | 0.888 | 0.983 |
| | Neuronal loss | -0.390 | 0.128 | [-0.641, -0.139] | 0.003 | 0.021* |
| Caudate | Age at death | 0.027 | 0.131 | [-0.230, 0.284] | 0.837 | 0.983 |
| | Education | -0.045 | 0.131 | [-0.302, 0.213] | 0.735 | 0.980 |
| | PMI | 0.133 | 0.135 | [-0.131, 0.397] | 0.329 | 0.808 |
| | Sex | 0.010 | 0.134 | [-0.253, 0.273] | 0.939 | 0.983 |
| | Neuronal loss | -0.290 | 0.130 | [-0.545, -0.036] | 0.029 | 0.058 |
| Hippocampus | Age at death | -0.071 | 0.131 | [-0.328, 0.186] | 0.592 | 0.901 |
| | Education | -0.100 | 0.131 | [-0.356, 0.157] | 0.449 | 0.901 |
| | PMI | 0.285 | 0.131 | [0.029, 0.541] | 0.033 | 0.804 |
| | Sex | 0.096 | 0.132 | [-0.162, 0.354] | 0.470 | 0.901 |
| | Neuronal loss | -0.248 | 0.133 | [-0.508, 0.012] | 0.067 | 0.101 |
| Pallidum | Age at death | -0.010 | 0.139 | [-0.283, 0.263] | 0.942 | 0.983 |
| | Education | 0.085 | 0.138 | [-0.187, 0.356] | 0.543 | 0.901 |
| | PMI | 0.003 | 0.141 | [-0.274, 0.280] | 0.983 | 0.983 |
| | Sex | 0.142 | 0.141 | [-0.134, 0.418] | 0.316 | 0.808 |
| | Neuronal loss | -0.143 | 0.138 | [-0.413, 0.128] | 0.305 | 0.367 |
| Putamen | Age at death | 0.141 | 0.126 | [-0.105, 0.387] | 0.264 | 0.808 |
| | Education | 0.148 | 0.126 | [-0.098, 0.394] | 0.244 | 0.808 |
| | PMI | 0.169 | 0.129 | [-0.083, 0.421] | 0.195 | 0.808 |
| | Sex | 0.080 | 0.128 | [-0.171, 0.331] | 0.535 | 0.901 |
| | Neuronal loss | -0.299 | 0.124 | [-0.542, -0.055] | 0.019 | 0.058 |
| Thalamus | Age at death | -0.201 | 0.135 | [-0.466, 0.064] | 0.143 | 0.808 |
| | Education | 0.029 | 0.135 | [-0.235, 0.293] | 0.829 | 0.983 |
| | PMI | 0.216 | 0.137 | [-0.052, 0.484] | 0.119 | 0.808 |
| | Sex | -0.073 | 0.138 | [-0.344, 0.198] | 0.600 | 0.901 |
| | Neuronal loss | -0.034 | 0.133 | [-0.294, 0.226] | 0.798 | 0.798 |

#### FTLD-Tau: Gliosis

| Structure | Predictor | β | SE | CI (95%) | p(raw) | p(FDR) |
|---|---|---|---|---|---|---|
| Amygdala | Age at death | -0.020 | 0.158 | [-0.330, 0.290] | 0.899 | 0.968 |
| | Education | -0.229 | 0.161 | [-0.544, 0.087] | 0.173 | 0.627 |
| | PMI | -0.417 | 0.152 | [-0.714, -0.119] | 0.013 | 0.319 |
| | Sex | -0.260 | 0.188 | [-0.628, 0.107] | 0.183 | 0.627 |
| | Gliosis | -0.733 | 0.165 | [-1.056, -0.411] | 0.000 | 0.001** |
| Caudate | Age at death | 0.377 | 0.193 | [-0.000, 0.755] | 0.065 | 0.520 |
| | Education | 0.047 | 0.242 | [-0.427, 0.522] | 0.846 | 0.968 |
| | PMI | -0.104 | 0.194 | [-0.485, 0.276] | 0.598 | 0.968 |
| | Sex | 0.442 | 0.251 | [-0.051, 0.934] | 0.094 | 0.568 |
| | Gliosis | -0.122 | 0.215 | [-0.544, 0.300] | 0.576 | 0.692 |
| Hippocampus | Age at death | -0.036 | 0.143 | [-0.316, 0.245] | 0.805 | 0.968 |
| | Education | -0.015 | 0.158 | [-0.324, 0.295] | 0.927 | 0.968 |
| | PMI | -0.342 | 0.142 | [-0.620, -0.063] | 0.026 | 0.319 |
| | Sex | 0.040 | 0.166 | [-0.285, 0.365] | 0.810 | 0.968 |
| | Gliosis | -0.685 | 0.139 | [-0.958, -0.411] | 0.000 | 0.000*** |
| Pallidum | Age at death | -0.250 | 0.219 | [-0.678, 0.179] | 0.267 | 0.802 |
| | Education | -0.197 | 0.250 | [-0.688, 0.294] | 0.441 | 0.968 |
| | PMI | -0.194 | 0.225 | [-0.634, 0.247] | 0.399 | 0.968 |
| | Sex | 0.117 | 0.256 | [-0.385, 0.619] | 0.652 | 0.968 |
| | Gliosis | 0.127 | 0.215 | [-0.294, 0.549] | 0.561 | 0.692 |
| Putamen | Age at death | 0.026 | 0.222 | [-0.409, 0.462] | 0.906 | 0.968 |
| | Education | -0.021 | 0.279 | [-0.569, 0.526] | 0.940 | 0.968 |
| | PMI | -0.139 | 0.224 | [-0.578, 0.300] | 0.543 | 0.968 |
| | Sex | 0.195 | 0.290 | [-0.373, 0.763] | 0.508 | 0.968 |
| | Gliosis | -0.242 | 0.249 | [-0.729, 0.245] | 0.342 | 0.684 |
| Thalamus | Age at death | -0.009 | 0.227 | [-0.454, 0.436] | 0.968 | 0.968 |
| | Education | 0.152 | 0.282 | [-0.400, 0.704] | 0.596 | 0.968 |
| | PMI | -0.039 | 0.232 | [-0.493, 0.416] | 0.869 | 0.968 |
| | Sex | 0.433 | 0.299 | [-0.152, 1.019] | 0.164 | 0.627 |
| | Gliosis | -0.027 | 0.251 | [-0.518, 0.464] | 0.914 | 0.914 |

#### FTLD-Tau: Neuronal loss

| Structure | Predictor | β | SE | CI (95%) | p(raw) | p(FDR) |
|---|---|---|---|---|---|---|
| Amygdala | Age at death | 0.044 | 0.151 | [-0.252, 0.340] | 0.774 | 0.963 |
| | Education | -0.140 | 0.158 | [-0.450, 0.170] | 0.387 | 0.891 |
| | PMI | -0.409 | 0.149 | [-0.701, -0.117] | 0.013 | 0.264 |
| | Sex | -0.186 | 0.177 | [-0.533, 0.161] | 0.307 | 0.891 |
| | Neuronal loss | -0.718 | 0.156 | [-1.024, -0.412] | 0.000 | 0.000*** |
| Caudate | Age at death | 0.384 | 0.187 | [0.017, 0.751] | 0.054 | 0.326 |
| | Education | 0.074 | 0.212 | [-0.342, 0.489] | 0.731 | 0.963 |
| | PMI | -0.063 | 0.193 | [-0.442, 0.316] | 0.749 | 0.963 |
| | Sex | 0.458 | 0.220 | [0.026, 0.889] | 0.051 | 0.326 |
| | Neuronal loss | -0.217 | 0.188 | [-0.585, 0.151] | 0.261 | 0.392 |
| Hippocampus | Age at death | -0.065 | 0.147 | [-0.354, 0.224] | 0.664 | 0.963 |
| | Education | -0.022 | 0.161 | [-0.338, 0.293] | 0.891 | 0.963 |
| | PMI | -0.362 | 0.145 | [-0.646, -0.077] | 0.022 | 0.264 |
| | Sex | 0.014 | 0.171 | [-0.321, 0.348] | 0.937 | 0.963 |
| | Neuronal loss | -0.682 | 0.144 | [-0.965, -0.400] | 0.000 | 0.000*** |
| Pallidum | Age at death | -0.259 | 0.220 | [-0.690, 0.172] | 0.253 | 0.891 |
| | Education | -0.211 | 0.250 | [-0.700, 0.278] | 0.408 | 0.891 |
| | PMI | -0.209 | 0.223 | [-0.646, 0.228] | 0.360 | 0.891 |
| | Sex | 0.104 | 0.256 | [-0.397, 0.605] | 0.687 | 0.963 |
| | Neuronal loss | 0.087 | 0.213 | [-0.332, 0.505] | 0.689 | 0.827 |
| Putamen | Age at death | 0.040 | 0.210 | [-0.372, 0.452] | 0.851 | 0.963 |
| | Education | 0.040 | 0.238 | [-0.426, 0.507] | 0.866 | 0.963 |
| | PMI | -0.070 | 0.217 | [-0.495, 0.356] | 0.752 | 0.963 |
| | Sex | 0.240 | 0.247 | [-0.244, 0.725] | 0.343 | 0.891 |
| | Neuronal loss | -0.375 | 0.211 | [-0.788, 0.039] | 0.091 | 0.183 |
| Thalamus | Age at death | -0.011 | 0.229 | [-0.460, 0.439] | 0.963 | 0.963 |
| | Education | 0.166 | 0.259 | [-0.342, 0.674] | 0.530 | 0.963 |
| | PMI | -0.036 | 0.231 | [-0.489, 0.416] | 0.876 | 0.963 |
| | Sex | 0.451 | 0.270 | [-0.077, 0.980] | 0.111 | 0.535 |
| | Neuronal loss | 0.008 | 0.226 | [-0.434, 0.450] | 0.971 | 0.971 |

**Supplementary Table S1.5. Mediation analyses for gliosis and neuronal loss.** For each diagnostic group, mediation analyses were performed with regional pathology as the predictor, gliosis or neuronal loss as mediators, and the ICV-normalized volume as the outcome. Standardized β coefficients for the direct and indirect effects, raw and FDR-corrected p-value (*p<0.05, **p<0.01, ***p<0.001) using the Benjamini–Hochberg procedure along with the proportion mediated and the 95% confidence interval are shown. Pathology burden ratings were not available for the nucleus accumbens, and therefore this structure is not shown.

#### Alzheimer's disease

| Structure | Mediator | Direct effect (β) | p (raw) Direct effect | p (FDR) Direct effect | Indirect effect (β) | p (raw) Indirect effect | p (FDR) Indirect effect | Proportion mediated | CI (95%) |
|---|---|---|---|---|---|---|---|---|---|
| Hippocampus | Gliosis | -0.467 | 0.011 | 0.092 | 0.067 | 0.603 | 0.804 | -0.169 | [-0.187, 0.335] |
| | Neuronal loss | -0.430 | 0.023 | 0.092 | 0.029 | 0.832 | 0.953 | -0.074 | [-0.238, 0.309] |
| Amygdala | Gliosis | -0.203 | 0.153 | 0.205 | -0.104 | 0.046 | 0.276 | 0.339 | [-0.250, -0.001] |
| | Neuronal loss | -0.159 | 0.271 | 0.326 | -0.148 | 0.017 | 0.211 | 0.484 | [-0.317, -0.023] |
| Caudate | Gliosis | -0.261 | 0.088 | 0.193 | -0.063 | 0.363 | 0.724 | 0.197 | [-0.217, 0.074] |
| | Neuronal loss | -0.222 | 0.151 | 0.205 | -0.103 | 0.182 | 0.468 | 0.317 | [-0.275, 0.049] |
| Putamen | Gliosis | -0.329 | 0.022 | 0.092 | -0.051 | 0.422 | 0.724 | 0.136 | [-0.192, 0.077] |
| | Neuronal loss | -0.285 | 0.049 | 0.148 | -0.095 | 0.177 | 0.468 | 0.253 | [-0.256, 0.044] |
| Thalamus | Gliosis | -0.077 | 0.059 | 0.617 | 0.000 | 0.983 | 0.983 | -0.008 | [-0.059, 0.064] |
| | Neuronal loss | -0.073 | 0.617 | 0.617 | -0.003 | 0.873 | 0.953 | 0.045 | [-0.060, 0.047] |
| Pallidum | Gliosis | 0.203 | 0.143 | 0.205 | -0.024 | 0.484 | 0.726 | -0.140 | [-0.117, 0.043] |
| | Neuronal loss | 0.227 | 0.096 | 0.193 | -0.049 | 0.195 | 0.468 | -0.278 | [-0.158, 0.019] |

#### FTLD-Tau

| Structure | Mediator | Direct effect (β) | p (raw) Direct effect | p (FDR) Direct effect | Indirect effect (β) | p (raw) Indirect effect | p (FDR) Indirect effect | Proportion mediated | CI (95%) |
|---|---|---|---|---|---|---|---|---|---|
| Hippocampus | Gliosis | -0.006 | 0.972 | 0.984 | -0.483 | 0.00 | 0.000*** | 0.991 | [-0.869, -0.176] |
| | Neuronal loss | -0.064 | 0.713 | 0.984 | -0.425 | 0.009 | 0.004** | 0.872 | [-0.799, -0.135] |
| Amygdala | Gliosis | -0.059 | 0.819 | 0.984 | -0.458 | 0.020 | 0.060 | 0.892 | [-0.922, -0.072] |
| | Neuronal loss | -0.136 | 0.532 | 0.984 | -0.380 | 0.010 | 0.041* | 0.739 | [-0.772, -0.081] |
| Caudate | Gliosis | 0.326 | 0.175 | 0.984 | 0.006 | 0.950 | 0.974 | 0.018 | [-0.173, 0.196] |
| | Neuronal loss | 0.364 | 0.112 | 0.984 | -0.032 | 0.758 | 0.974 | -0.096 | [-0.285, 0.195] |
| Putamen | Gliosis | -0.044 | 0.871 | 0.984 | 0.008 | 0.943 | 0.974 | -0.235 | [-0.223, 0.250] |
| | Neuronal loss | 0.005 | 0.984 | 0.984 | -0.041 | 0.750 | 0.974 | 1.180 | [-0.359, 0.242] |
| Thalamus | Gliosis | -0.117 | 0.676 | 0.984 | -0.014 | 0.878 | 0.974 | 0.110 | [-0.260, 0.225] |
| | Neuronal loss | -0.136 | 0.631 | 0.984 | 0.004 | 0.974 | 0.974 | -0.030 | [-0.239, 0.261] |
| Pallidum | Gliosis | -0.245 | 0.415 | 0.984 | 0.043 | 0.699 | 0.974 | -0.216 | [-0.176, 0.331] |
| | Neuronal loss | -0.219 | 0.462 | 0.984 | 1.740 | 0.876 | 0.974 | -0.086 | [-0.203, 0.269] |
