## Supplementary File 2 for "Postmortem brain MRI reveals differential associations of subcortical and limbic volumes with cortical thinning and neurodegenerative pathologies"

### Supplementary File 2: Five disease diagnostic groups

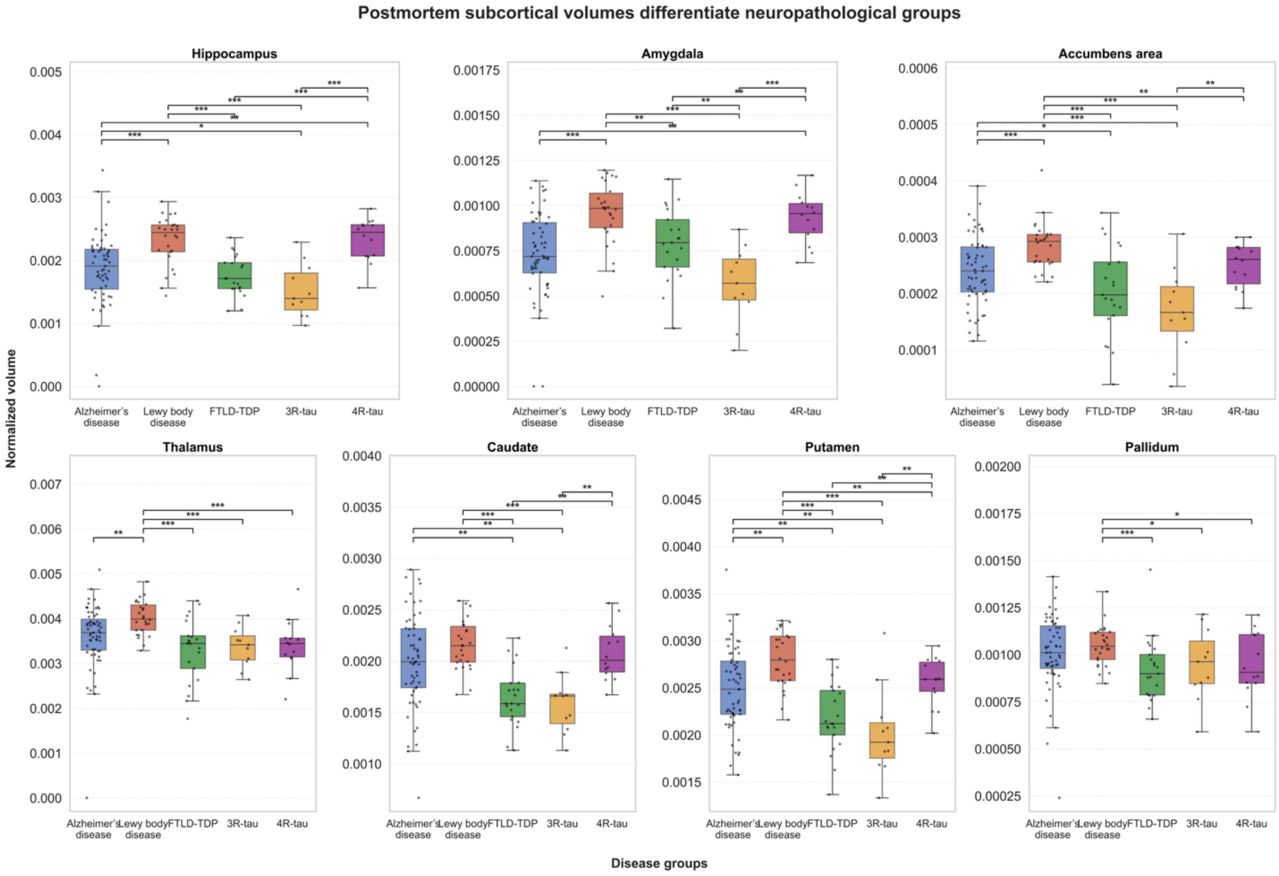


**Supplementary Figure S2.1. Postmortem limbic and subcortical volumes across neuropathological groups.** Boxplots show ICV-normalized postmortem hemisphere volumes for the subcortical/limbic structures across the five diagnostic categories: Alzheimer's disease (AD), Lewy body disease (LBD), Frontotemporal Lobar Degeneration with TDP-43 pathology (FTLD-TDP), 3R (Pick's disease) and 4R (PSP and CBD) tauopathies. Pairwise differences between groups were assessed using likelihood-ratio tests adjusting for age at death, sex, education, and postmortem interval. Horizontal bars indicate statistically significant pairwise contrasts ($p<0.05$; p<0.01; *p<0.001) after False Discovery Rate (FDR) correction using the Benjamini–Hochberg procedure, applied independently within each diagnostic group's set of pairwise comparisons.

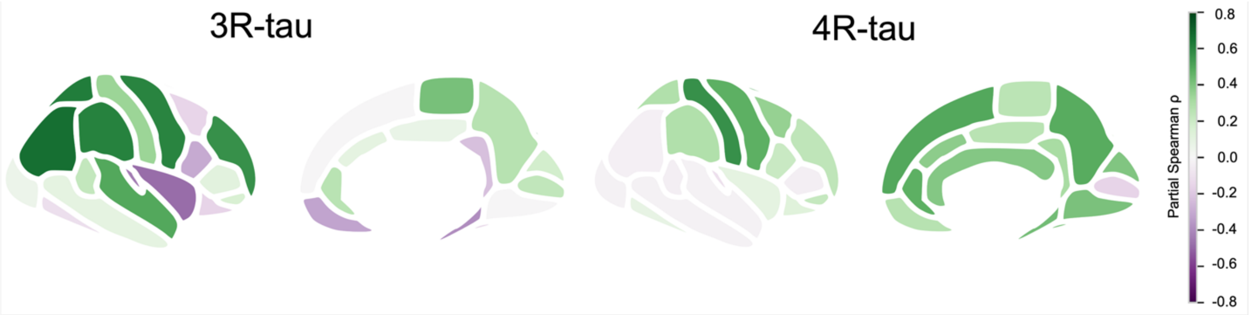


**Supplementary Figure S2.2. Associations between Desikan-Killiany-Tourville (DKT) atlas-based regional mean thickness and weighted subcortical and limbic volumes.** A weighted composite across the seven nuclei (hippocampus, amygdala, caudate, putamen, pallidum, thalamus, and nucleus accumbens), with weights proportional to each structure's native volume. The composite DGM value was further ICV-normalized by antemortem intracranial volume (ICV). For each neuropathological group (Alzheimer's disease, Lewy body disease, FTLD-TDP, 3R tauopathy, and 4R tauopathy), partial Spearman correlations were calculated between mean ROI-level Desikan-Killiany-Tourville (DKT) cortical thickness and the normalized weighted DGM volume, adjusting for age at death, sex, education, and postmortem interval. Reported are the correlation coefficients (ρ), with regions outlined in black indicating associations that survived false discovery rate correction using the Benjamini–Hochberg procedure applied within each diagnostic group.

**(A)**

Partial Spearman correlation between postmortem volume and regional pathology burden

| | Alzheimer's disease (p-tau) | Lewy body disease (α-synuclein) | FTLD-TDP (TDP-43) | 3R-tauopathy (p-tau) | 4R-tauopathy (p-tau) |
|---|---|---|---|---|---|
| Hippocampus | ρ = -0.33* (p = 0.013) | ρ = -0.08 (p = 0.720) | ρ = 0.17 (p = 0.520) | ρ = 0.46 (p = 0.301) | ρ = 0.09 (p = 0.811) |
| Amygdala | ρ = -0.22 (p = 0.108) | ρ = 0.00 (p = 0.989) | ρ = 0.04 (p = 0.871) | Not Applicable | ρ = 0.08 (p = 0.827) |
| Caudate | ρ = -0.31* (p = 0.018) | ρ = -0.33 (p = 0.137) | ρ = -0.33 (p = 0.208) | ρ = 0.54 (p = 0.206) | ρ = 0.50 (p = 0.138) |
| Putamen | ρ = -0.40* (p = 0.002) | ρ = -0.29 (p = 0.196) | ρ = -0.54 (p = 0.030) | ρ = 0.78 (p = 0.039) | ρ = -0.14 (p = 0.691) |
| Thalamus | ρ = -0.06 (p = 0.650) | ρ = -0.60* (p = 0.004) | ρ = -0.29 (p = 0.284) | ρ = 0.15 (p = 0.741) | ρ = -0.09 (p = 0.804) |
| Pallidum | ρ = -0.06 (p = 0.658) | ρ = -0.34 (p = 0.122) | ρ = 0.20 (p = 0.468) | ρ = -0.31 (p = 0.493) | ρ = -0.22 (p = 0.538) |

Partial Spearman ρ: 1.00, 0.75, 0.50, 0.25, 0.00, −0.25, −0.50, −0.75, −1.00

**(B)**

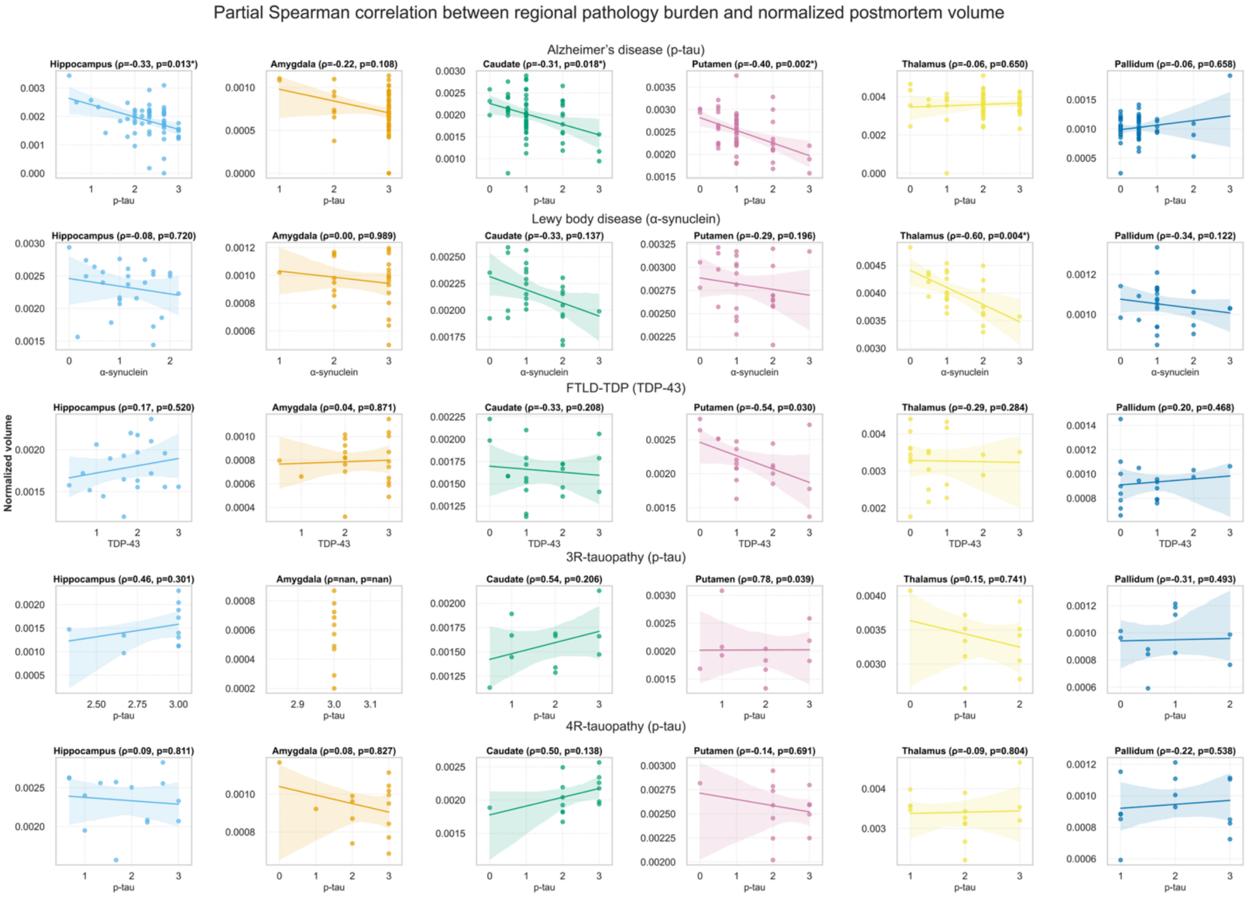


**Supplementary Figure S2.3. Ordinary least squares model analysis and partial Spearman correlations between subcortical/limbic volumes and primary neuropathological burden.** (**A**) Heatmaps show standardized β coefficients from OLS models quantifying the association between subcortical/limbic postmortem MRI volumes and regional semi-quantitative ratings of primary molecular pathology within each diagnostic group (Alzheimer's disease, Lewy body disease, FTLD-TDP, 3R tauopathy, and 4R tauopathy). Each cell reports the standardized effect size (β) and the corresponding uncorrected p value; boxes highlight uncorrected p<0.05, and asterisks denote significance after false discovery rate correction using the Benjamini–Hochberg procedure applied separately within each diagnostic group across all structures and pathology predictors (*p<0.05 (also outlined), **p<0.01, ***p<0.001). All models included age at death, sex, years of education, and postmortem interval as covariates. (**B**) shows partial Spearman correlation coefficients between ICV-normalized postmortem structure volumes and their corresponding molecular pathology ratings within each diagnostic group. Correlations were adjusted for age at death, sex, postmortem interval, and years of education and corrected for multiple comparisons.

Note: Some entries appear as "Not Applicable" because certain region-specific pathology markers exhibited zero variability within specific diagnostic groups (for example, in Panel B we notice that the amygdala pathology ratings in the 3R tauopathy group were confined to a single value). When a predictor lacks sufficient within-group variance, neither partial Spearman correlations nor OLS regression coefficients can be estimated reliably. Such cases would otherwise yield β values near zero with spuriously small p-values. These predictors were therefore excluded from model fitting and reported as not applicable. For the OLS analysis, an independent linear model was fit for each structure within each diagnostic group, including the region's predominant pathology measure together with covariates (age at death, sex, education, and postmortem interval). After removing predictors with zero variance, all retained predictors across all structures within a diagnostic group were corrected for multiple comparisons using the Benjamini–Hochberg FDR procedure

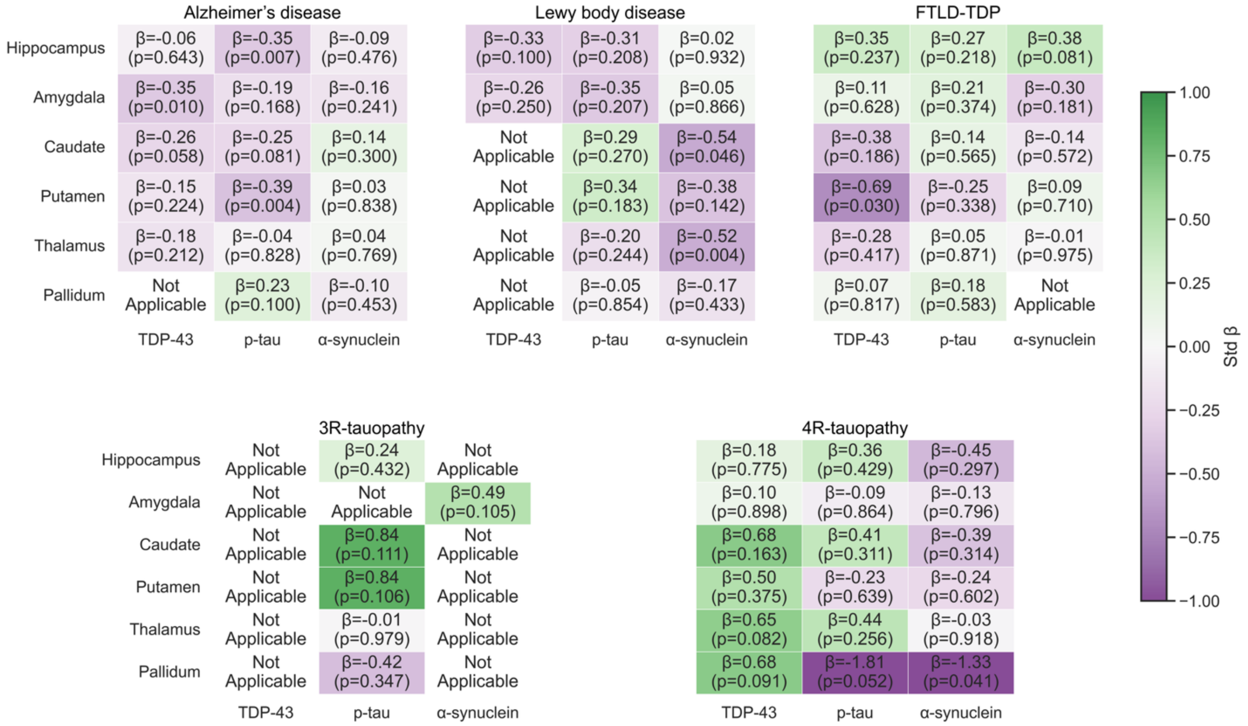


**Supplementary Figure S2.4. Polypathology ordinary least squares models for structure-pathology relationships.** Heatmaps show standardized β coefficients from OLS models quantifying the association between regional postmortem volumes (rows) and multiple semi-quantitative pathology ratings (columns) across the five diagnostic groups (Alzheimer's disease, Lewy body disease, FTLD-TDP, 3R tauopathy, and 4R tauopathy). Each cell reports the standardized effect size (β) and the corresponding uncorrected p-value; boxes highlight uncorrected $p<0.05$, and asterisks denote significance after false discovery rate correction using the Benjamini–Hochberg procedure applied separately within each diagnostic group across all structures and pathology predictors (*$p<0.05$, **$p<0.01$, ***$p<0.001$). All models included age at death, sex, years of education, and postmortem interval as covariates.

Some predictors appear as "Not applicable" because several pathology markers exhibited minimal or zero variability within particular disease groups. Predictors lacking sufficient within-group variance cannot yield interpretable regression estimates and may otherwise produce β values near zero with spuriously small p-values. Consistent with the approach described for Figure S2.3, such predictors were removed prior to model fitting and excluded from FDR correction. Pathology burden ratings were not available for the nucleus accumbens, and therefore this structure is not shown.

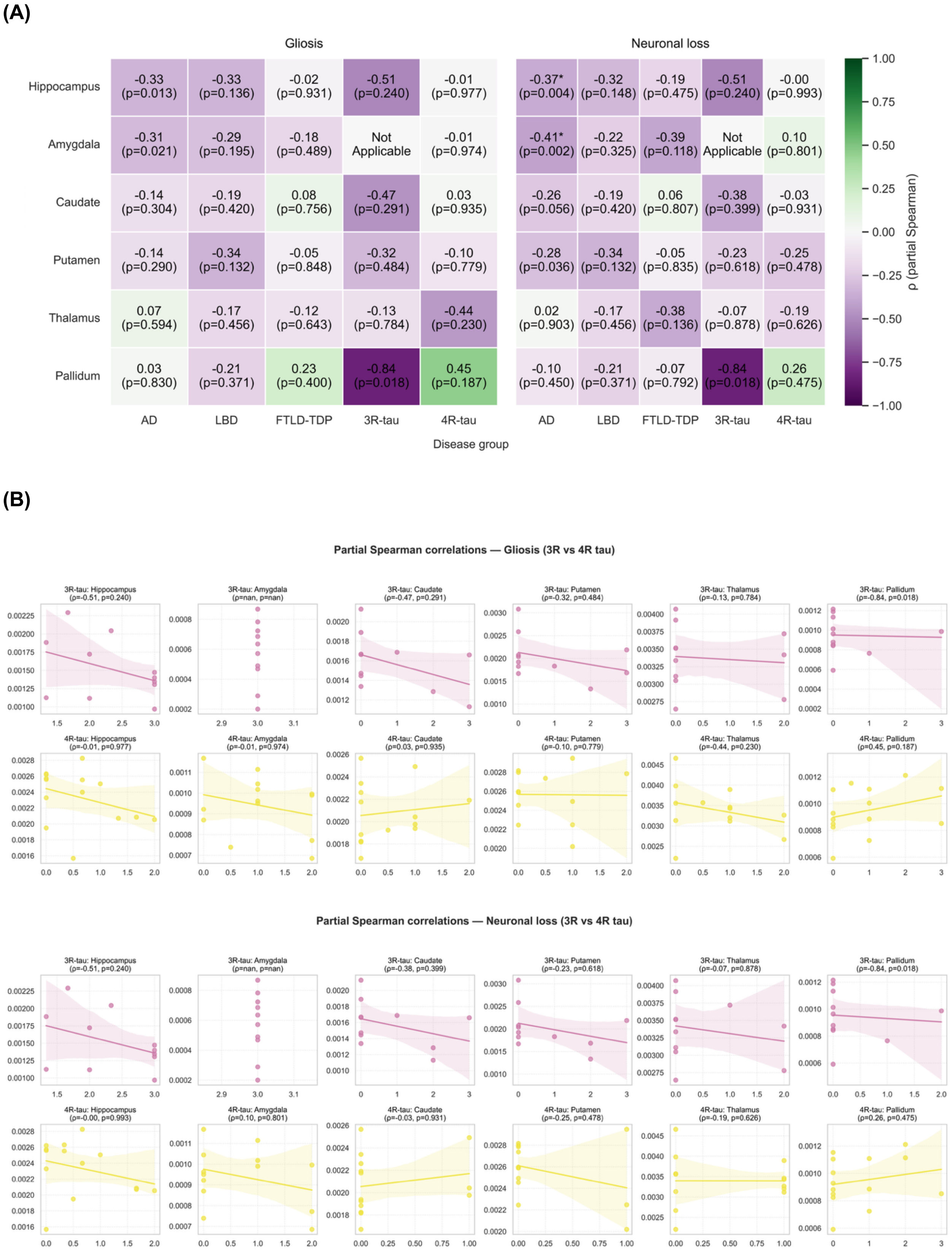

(A)
Gliosis
Neuronal loss
Hippocampus
Amygdala
Caudate
Putamen
Thalamus
Pallidum
-0.33 (p=0.013)
-0.33 (p=0.136)
-0.02 (p=0.931)
-0.51 (p=0.240)
-0.01 (p=0.977)
-0.31 (p=0.021)
-0.29 (p=0.195)
-0.18 (p=0.489)
Not Applicable
-0.01 (p=0.974)
-0.14 (p=0.304)
-0.19 (p=0.420)
0.08 (p=0.756)
-0.47 (p=0.291)
0.03 (p=0.935)
-0.14 (p=0.290)
-0.34 (p=0.132)
-0.05 (p=0.848)
-0.32 (p=0.484)
-0.10 (p=0.779)
0.07 (p=0.594)
-0.17 (p=0.456)
-0.12 (p=0.643)
-0.13 (p=0.784)
-0.44 (p=0.230)
0.03 (p=0.830)
-0.21 (p=0.371)
0.23 (p=0.400)
-0.84 (p=0.018)
0.45 (p=0.187)
-0.37* (p=0.004)
-0.32 (p=0.148)
-0.19 (p=0.475)
-0.51 (p=0.240)
-0.00 (p=0.993)
-0.41* (p=0.002)
-0.22 (p=0.325)
-0.39 (p=0.118)
Not Applicable
0.10 (p=0.801)
-0.26 (p=0.056)
-0.19 (p=0.420)
0.06 (p=0.807)
-0.38 (p=0.399)
-0.03 (p=0.931)
-0.28 (p=0.036)
-0.34 (p=0.132)
-0.05 (p=0.835)
-0.23 (p=0.618)
-0.25 (p=0.478)
0.02 (p=0.903)
-0.17 (p=0.456)
-0.38 (p=0.136)
-0.07 (p=0.878)
-0.19 (p=0.626)
-0.10 (p=0.450)
-0.21 (p=0.371)
-0.07 (p=0.792)
-0.84 (p=0.018)
0.26 (p=0.475)
AD
LBD
FTLD-TDP
3R-tau
4R-tau
Disease group
ρ (partial Spearman)
1.00
0.75
0.50
0.25
0.00
−0.25
−0.50
−0.75
−1.00
(B)
Partial Spearman correlations — Gliosis (3R vs 4R tau)
3R-tau: Hippocampus (ρ=-0.51, p=0.240)
3R-tau: Amygdala (ρ=nan, p=nan)
3R-tau: Caudate (ρ=-0.47, p=0.291)
3R-tau: Putamen (ρ=-0.32, p=0.484)
3R-tau: Thalamus (ρ=-0.13, p=0.784)
3R-tau: Pallidum (ρ=-0.84, p=0.018)
4R-tau: Hippocampus (ρ=-0.01, p=0.977)
4R-tau: Amygdala (ρ=-0.01, p=0.974)
4R-tau: Caudate (ρ=0.03, p=0.935)
4R-tau: Putamen (ρ=-0.10, p=0.779)
4R-tau: Thalamus (ρ=-0.44, p=0.230)
4R-tau: Pallidum (ρ=0.45, p=0.187)
Partial Spearman correlations — Neuronal loss (3R vs 4R tau)
3R-tau: Hippocampus (ρ=-0.51, p=0.240)
3R-tau: Amygdala (ρ=nan, p=nan)
3R-tau: Caudate (ρ=-0.38, p=0.399)
3R-tau: Putamen (ρ=-0.23, p=0.618)
3R-tau: Thalamus (ρ=-0.07, p=0.878)
3R-tau: Pallidum (ρ=-0.84, p=0.018)
4R-tau: Hippocampus (ρ=-0.00, p=0.993)
4R-tau: Amygdala (ρ=0.10, p=0.801)
4R-tau: Caudate (ρ=-0.03, p=0.931)
4R-tau: Putamen (ρ=-0.25, p=0.478)
4R-tau: Thalamus (ρ=-0.19, p=0.626)
4R-tau: Pallidum (ρ=0.26, p=0.475)

**Supplementary Figure S2.5. Structure-histology relationships for gliosis and neuronal loss.** (**A**) Heatmaps show standardized β coefficients from OLS models quantifying the association between subcortical/limbic postmortem MRI volumes and regional semi-quantitative ratings for gliosis and neuronal loss markers across the five diagnostic groups (Alzheimer's disease, Lewy body disease, FTLD-TDP, 3R and 4R tauopathies). Each cell reports the standardized effect size (β) and the corresponding uncorrected p value; boxes highlight uncorrected $p<0.05$, and asterisks denote significance after false discovery rate correction using the Benjamini–Hochberg procedure applied separately within each diagnostic group across all structures and pathology predictors (*$p<0.05$ (also outlined), **$p<0.01$, ***$p<0.001$). All models included age at death, sex, years of education, and postmortem interval as covariates. Pathology burden ratings were not available for the nucleus accumbens, so this structure is not shown. *Mediation analysis could not be performed for the 3R and 4R groups as the direct effect did not reach significance.* **(B)** Partial Spearman correlation between postmortem MRI volumes and regional mediation markers were calculated between postmortem subcortical MRI volumes and semi-quantitative ratings of gliosis and neuronal loss for each group. All correlations were adjusted for age at death, sex, postmortem interval and education. Heatmap colors reflect correlation strength (ρ), and values within each cell indicate correlation coefficients with the corresponding uncorrected p-value (in parentheses, boxes for $p<0.05$) with asterisks denoting significance after false discovery rate (FDR) correction applied within group across all structures (*$p<0.05$, **$p<0.01$, ***$p<0.001$).

Some predictors appear as "Not applicable" because several histology markers exhibited minimal or zero variability within disease groups. Predictors lacking sufficient within-group variance cannot yield interpretable regression estimates and may otherwise produce β values near zero with spuriously small p-values. Consistent with the approach described for Figure S2.3, such predictors were removed prior to model fitting and excluded from FDR correction. Pathology burden ratings were not available for the nucleus accumbens, and therefore this structure is not shown.

(A)

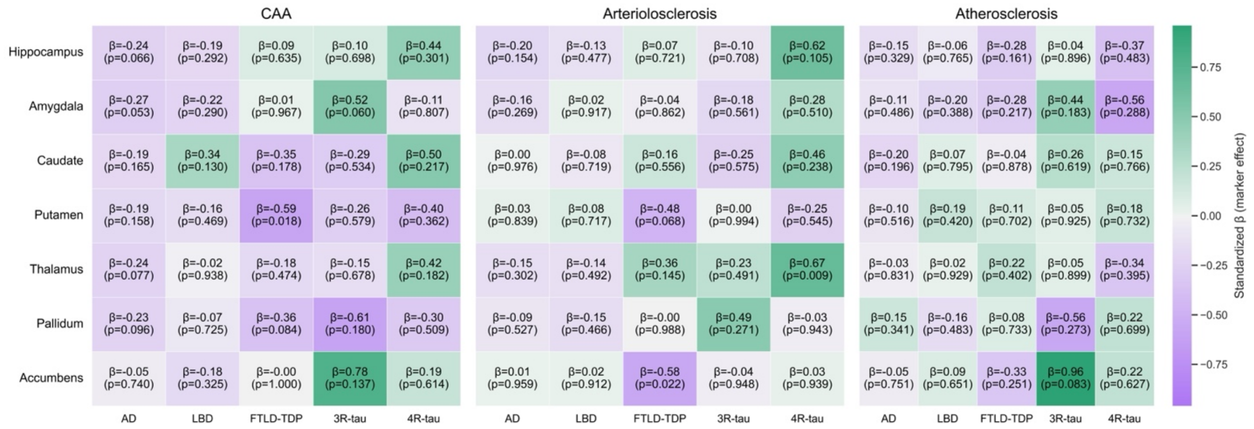
CAA
Arteriolosclerosis
Atherosclerosis
Hippocampus
Amygdala
Caudate
Putamen
Thalamus
Pallidum
Accumbens
AD
LBD
FTLD-TDP
3R-tau
4R-tau
Standardized β (marker effect)
0.75
0.50
0.25
0.00
−0.25
−0.50
−0.75
β=-0.24 (p=0.066) β=-0.19 (p=0.292) β=0.09 (p=0.635) β=0.10 (p=0.698) β=0.44 (p=0.301)
β=-0.27 (p=0.053) β=-0.22 (p=0.290) β=0.01 (p=0.967) β=0.52 (p=0.060) β=-0.11 (p=0.807)
β=-0.19 (p=0.165) β=0.34 (p=0.130) β=-0.35 (p=0.178) β=-0.29 (p=0.534) β=0.50 (p=0.217)
β=-0.19 (p=0.158) β=-0.16 (p=0.469) β=-0.59 (p=0.018) β=-0.26 (p=0.579) β=-0.40 (p=0.362)
β=-0.24 (p=0.077) β=-0.02 (p=0.938) β=-0.18 (p=0.474) β=-0.15 (p=0.678) β=0.42 (p=0.182)
β=-0.23 (p=0.096) β=-0.07 (p=0.725) β=-0.36 (p=0.084) β=-0.61 (p=0.180) β=-0.30 (p=0.509)
β=-0.05 (p=0.740) β=-0.18 (p=0.325) β=-0.00 (p=1.000) β=0.78 (p=0.137) β=0.19 (p=0.614)
β=-0.20 (p=0.154) β=-0.13 (p=0.477) β=0.07 (p=0.721) β=-0.10 (p=0.708) β=0.62 (p=0.105)
β=-0.16 (p=0.269) β=0.02 (p=0.917) β=-0.04 (p=0.862) β=-0.18 (p=0.561) β=0.28 (p=0.510)
β=0.00 (p=0.976) β=-0.08 (p=0.719) β=0.16 (p=0.556) β=-0.25 (p=0.575) β=0.46 (p=0.238)
β=0.03 (p=0.839) β=0.08 (p=0.717) β=-0.48 (p=0.068) β=0.00 (p=0.994) β=-0.25 (p=0.545)
β=-0.15 (p=0.302) β=-0.14 (p=0.492) β=0.36 (p=0.145) β=0.23 (p=0.491) β=0.67 (p=0.009)
β=-0.09 (p=0.527) β=-0.15 (p=0.466) β=-0.00 (p=0.988) β=0.49 (p=0.271) β=-0.03 (p=0.943)
β=0.01 (p=0.959) β=0.02 (p=0.912) β=-0.58 (p=0.022) β=-0.04 (p=0.948) β=0.03 (p=0.939)
β=-0.15 (p=0.329) β=-0.06 (p=0.765) β=-0.28 (p=0.161) β=0.04 (p=0.896) β=-0.37 (p=0.483)
β=-0.11 (p=0.486) β=-0.20 (p=0.388) β=-0.28 (p=0.217) β=0.44 (p=0.183) β=-0.56 (p=0.288)
β=-0.20 (p=0.196) β=0.07 (p=0.795) β=-0.04 (p=0.878) β=0.26 (p=0.619) β=0.15 (p=0.766)
β=-0.10 (p=0.516) β=0.19 (p=0.420) β=0.11 (p=0.702) β=0.05 (p=0.925) β=0.18 (p=0.732)
β=-0.03 (p=0.831) β=0.02 (p=0.929) β=0.22 (p=0.402) β=0.05 (p=0.899) β=-0.34 (p=0.395)
β=0.15 (p=0.341) β=-0.16 (p=0.483) β=0.08 (p=0.733) β=-0.56 (p=0.273) β=0.22 (p=0.699)
β=-0.05 (p=0.751) β=0.09 (p=0.651) β=-0.33 (p=0.251) β=0.96 (p=0.083) β=0.22 (p=0.627)

(B)

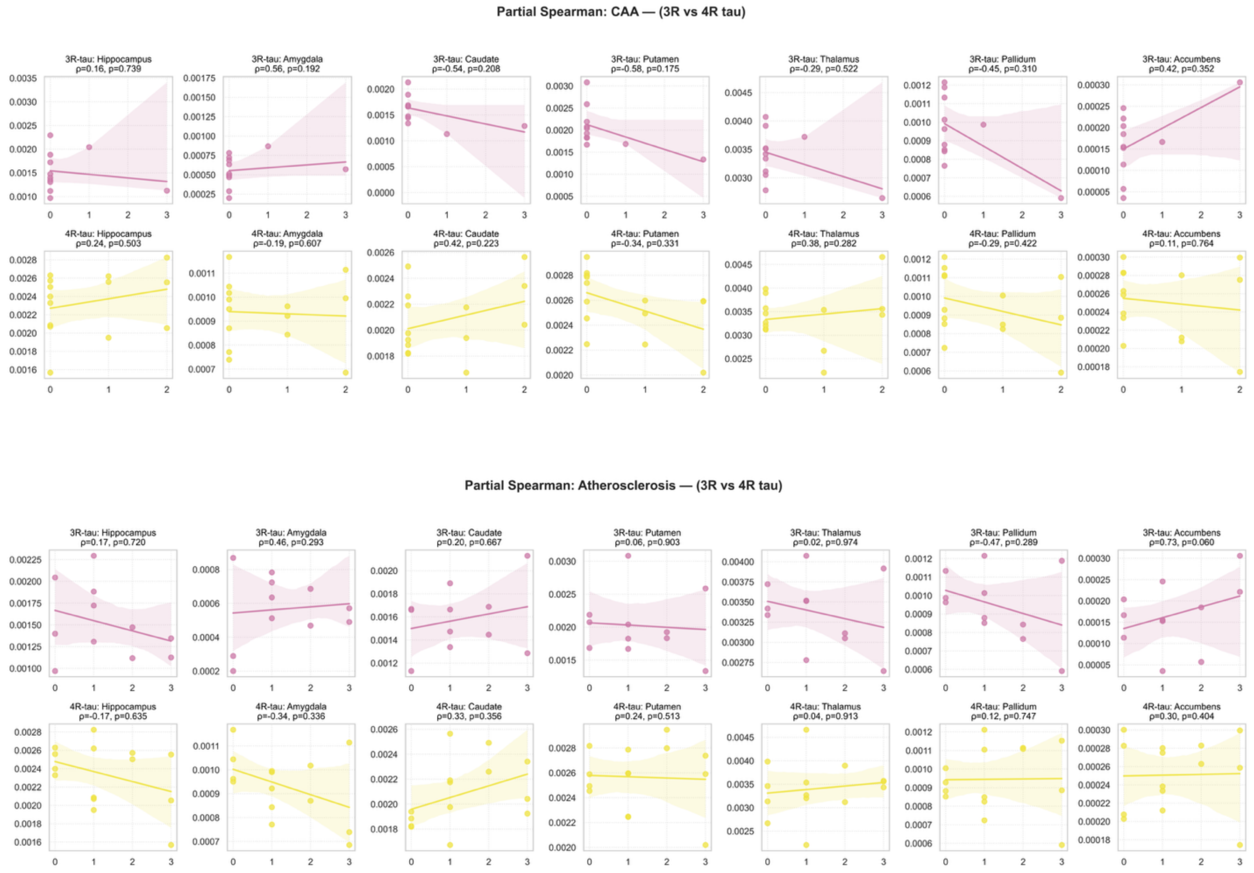
Partial Spearman: CAA — (3R vs 4R tau)
3R-tau: Hippocampus ρ=0.16, p=0.739
3R-tau: Amygdala ρ=0.56, p=0.192
3R-tau: Caudate ρ=-0.54, p=0.208
3R-tau: Putamen ρ=-0.58, p=0.175
3R-tau: Thalamus ρ=-0.29, p=0.522
3R-tau: Pallidum ρ=-0.45, p=0.310
3R-tau: Accumbens ρ=0.42, p=0.352
4R-tau: Hippocampus ρ=0.24, p=0.503
4R-tau: Amygdala ρ=-0.19, p=0.607
4R-tau: Caudate ρ=0.42, p=0.223
4R-tau: Putamen ρ=-0.34, p=0.331
4R-tau: Thalamus ρ=0.38, p=0.282
4R-tau: Pallidum ρ=-0.29, p=0.422
4R-tau: Accumbens ρ=0.11, p=0.764
Partial Spearman: Atherosclerosis — (3R vs 4R tau)
3R-tau: Hippocampus ρ=0.17, p=0.720
3R-tau: Amygdala ρ=0.46, p=0.293
3R-tau: Caudate ρ=0.20, p=0.667
3R-tau: Putamen ρ=0.06, p=0.903
3R-tau: Thalamus ρ=0.02, p=0.974
3R-tau: Pallidum ρ=-0.47, p=0.289
3R-tau: Accumbens ρ=0.73, p=0.060
4R-tau: Hippocampus ρ=-0.17, p=0.635
4R-tau: Amygdala ρ=-0.34, p=0.336
4R-tau: Caudate ρ=0.33, p=0.356
4R-tau: Putamen ρ=0.24, p=0.513
4R-tau: Thalamus ρ=0.04, p=0.913
4R-tau: Pallidum ρ=0.12, p=0.747
4R-tau: Accumbens ρ=0.30, p=0.404

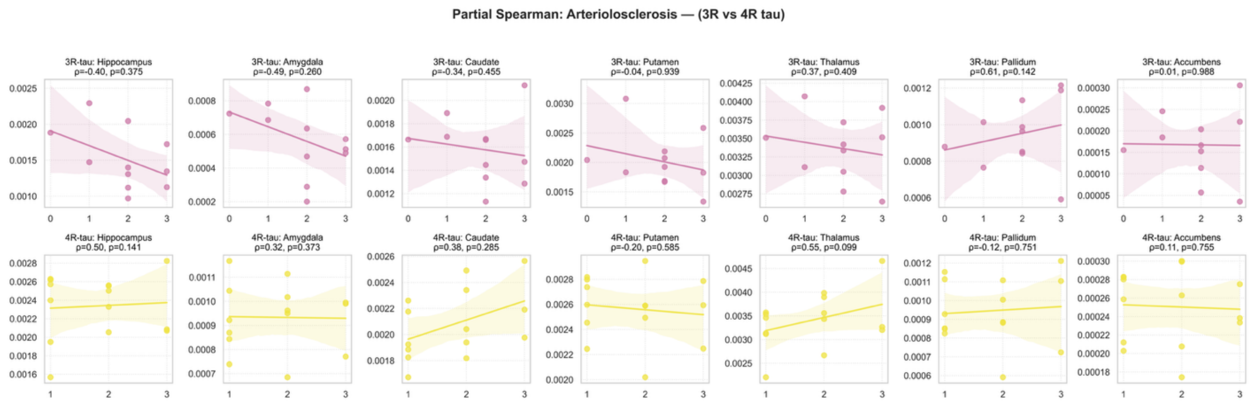


**Supplementary Figure 2.6. Partial Spearman correlations and ordinary least squares model between postmortem volumes and vascular burden.** (**A**) Heatmaps show standardized β coefficients from OLS models quantifying the association between subcortical/limbic postmortem MRI volumes and global semi-quantitative markers of cerebral amyloid angiopathy (CAA), and vascular pathology (arteriolosclerosis and atherosclerosis) across Alzheimer's disease, Lewy body disease, FTLD-TDP, 3R and 4R tauopathies. Each cell reports the standardized effect size (β) and the corresponding uncorrected p value; boxes highlight uncorrected p<0.05, and asterisks denote significance after false discovery rate correction using the Benjamini–Hochberg procedure applied separately within each diagnostic group across all structures and pathology predictors (*p<0.05 (also outlined), **p<0.01, ***p<0.001). All models included age at death, sex, years of education, and postmortem interval as covariates. (**B**) Partial Spearman correlation coefficients between ICV-normalized postmortem structure volumes and their corresponding histology marker ratings within each diagnostic group. Correlations were adjusted for age at death, sex, postmortem interval, and years of education and corrected for multiple comparisons.
